# Regioselective biosynthesis of oligoamides as precursors for sequence-controlled co-polyamides

**DOI:** 10.1101/2025.11.03.685623

**Authors:** Liangyu Qian, Isaiah T. Dishner, Nikolas Capra, Dana L. Carper, Nduka Ogbonna, Vilmos Kertesz, John F. Cahill, Jeffrey C. Foster, Joshua K. Michener

**Affiliations:** Biosciences, Oak Ridge National Laboratory, Oak Ridge, TN; Chemical Sciences, Oak Ridge National Laboratory, Oak Ridge, TN; Neutron Scattering Divisions, Oak Ridge National Laboratory, Oak Ridge, TN

## Abstract

Polyamides are important natural and synthetic polymers, best exemplified by proteins and nylons respectively. Proteins demonstrate that novel polymers with emergent properties can be generated by combining diverse monomers in precisely defined sequences. However, commercial polyamides represent only a small fraction of the potential diversity in polyamide sequences, due to the synthetic challenges of sequence-controlled polymerization. Amide synthetases have been shown to synthesize a broad array of nylon-relevant diads, but the generation of novel sequenced copolyamides requires enzymes capable of acting with longer and more diverse substrates. In this study, we demonstrated that NRPS-independent siderophore (NIS) synthetases, represented by DesD, can ligate oligomeric substrates. A simultaneous enzyme cascade using DesD enabled the synthesis of an oligotriad directly from unprotected substrates. Moreover, the regioselectivity of DesD allowed the selective synthesis of a sequenced amide tetrad, the precursor to a novel sequenced-defined polyamide with properties superior to nylon 66. This study establishes a direct biocatalytic route for the facile synthesis of sequenced oligoamides from unprotected bifunctional substrates, opening new possibilities to synthesize sequenced copolyamides.

## Introduction

Nylons, the most widely produced synthetic polyamides, are extensively used in applications ranging from textiles to automotive parts due to their excellent mechanical and thermal properties. A variety of nylon backbones are commercially available, but due to practical synthetic considerations are typically limited to either homopolymers of a single ⍵-amino acid (e.g. nylon 6), combinations of one diacid and one diamine (nylon 66), or block copolymers of the same. While those nylons provide different properties based on the repeat units of the polymer, studies have increasingly explored the incorporation of complex and multi-component repeat units to develop polymers with novel properties.^1,2^ However, achieving precise sequence control during polymerization remains a significant challenge, limiting the ability to tailor material properties.^3,4^ Very few examples of sequenced synthetic polyamides have been described.^5^

Nature provides a powerful model for sequence-controlled polyamide, as exemplified by proteins (Figure 1). Proteins are natural polyamides synthesized by ribosomes and primarily composed of twenty natural α-amino acids connected by peptide (amide) bonds and arranged in specific sequences. The resulting polyamides demonstrate emergent properties based on structured interactions between sequenced monomers (Figure 1). However, ribosomes are largely limited to canonical α-amino acids as substrates, and it still remains a significant engineering challenge to expand their substrate scope.^6–11^

**Figure 1.**
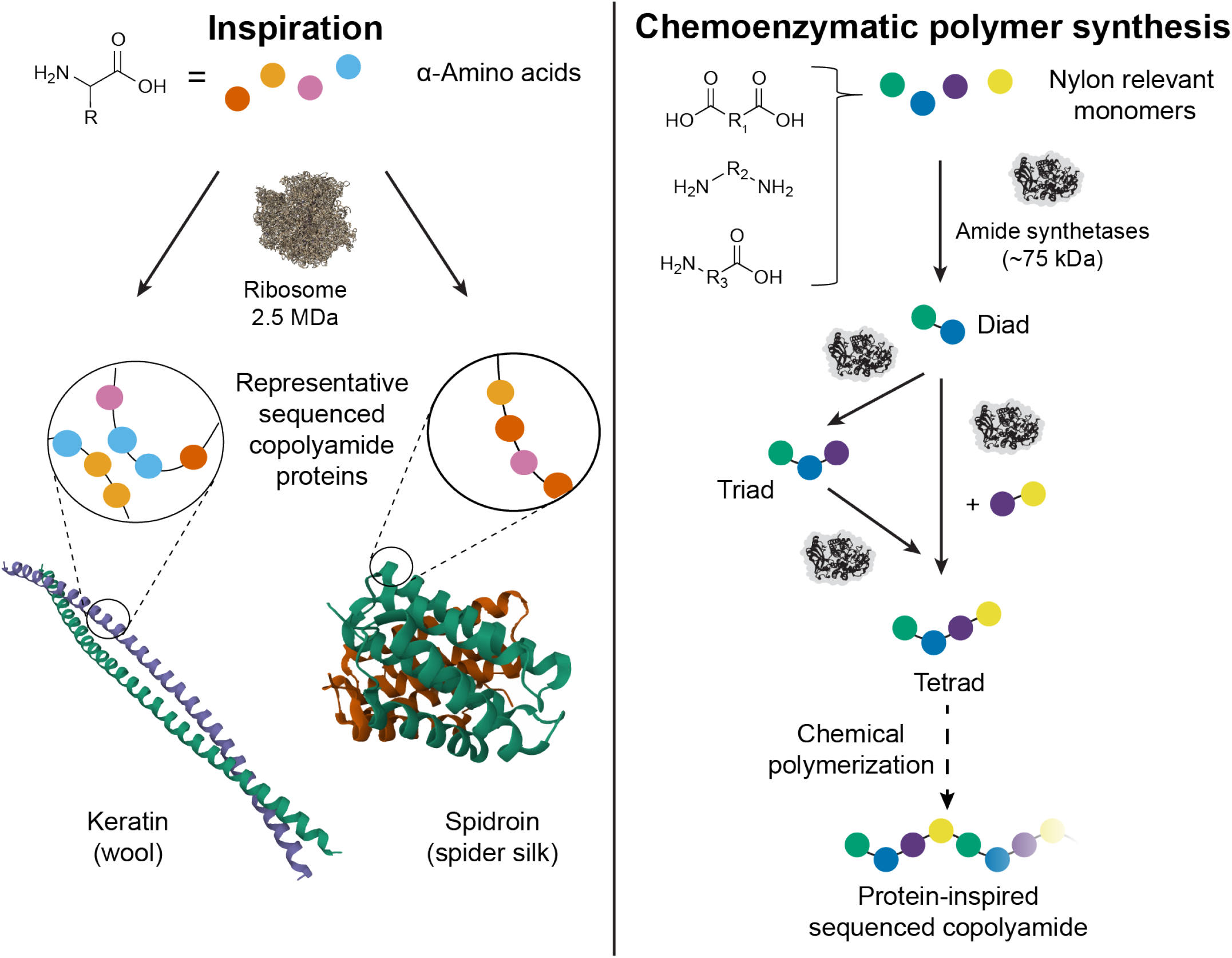
Enzymatic synthesis of sequenced polyamides. The emergent properties of ribosomally-synthesized sequenced copolyamide proteins provides inspiration for the use of amide synthetases to synthesize nylon-relevant oligoamides that can be chemically polymerized into sequenced copolyamides.

Alternatively, using enzymes to selectively couple nylon-like substrates could lead to polymeric materials with novel properties. Our previous work demonstrated that amide synthetases can couple a variety of polymer-relevant substrates to synthesize amide diads that can serve as nylon precursors.^12^ However, synthesizing multi-component oligoamides, such as triads and tetrads, requires enzymes capable of ligating longer and more diverse substrates (Figure 1). Non-ribosomal peptide synthetase-independent siderophore (NIS) synthetases, which catalyze ATP-dependent ligation of carboxylates with nucleophiles in siderophore biosynthesis, are promising candidates due to their ability to natively ligate longer substrates.^13–16^ NIS synthetases are classified into three types based on substrate specificity: Type A (citrate),^17,18^ Type B (α-ketoglutarate),^19,20^ and Type C (citrate or succinate derivatives).^21–23^ These three enzyme classes exhibit low sequence identity (20–30%), reflecting their diverse substrate preferences. However, their potential for ligation of nylon-relevant substrates remained unexplored.

In this study, we demonstrated that NIS synthetases are active with polymer-relevant substrates and provide unique activities compared to previously characterized enzymes. Notably, DesD, a Type C NIS synthetase, enabled the synthesis of diverse oligoamides. Using an enzyme cascade involving two enzymes, we synthesized an ⍵-amino acid triad directly from unprotected substrates. Furthermore, DesD catalyzed regioselective ligation of asymmetric oligomeric substrates, providing the basis of sequence control over oligoamide synthesis. The resulting oligoamides can be further polymerized into sequenced polymers with properties superior to nylon 66. Our study not only provides a viable method for enzymatically synthesizing multi-component nylon oligomers, but also facilitates the characterization of polyamide sequence-function relationships and the design of new polymers.

## Results and discussion

### NIS synthetases are active with polymer-relevant substrates

Our previous work demonstrated that DdaG and SfaB can couple diacid and diamine substrates to form a variety of ⍵-amino acid diads.^12^ However, these enzymes were unable to synthesize other diad types, such as diamine diads, which prompted us to identify enzymes with broader substrate scope. Four NIS synthetases, namely AsbA, AsbB, AcsA and DesD, were selected and purified as hexahistidine-tagged proteins (Figure S1) to test their activities *in vitro*. We first validated the published activities of AsbA to ligate citrate and spermidine,^17^ and DesD to dimerize *N*-succinylcadaverine,^23^ detecting oligomeric products using IDOT/OPSI-MS analysis.^24^ Next, we tested these enzymes, as well as previously identified amide synthetases DdaG^25^ and SfaB^26^, with polymer-relevant substrates. Results from direct injection immediate droplet-on-demand/open port sampling interface-mass spectrometry (IDOT/OPSI-MS) and MS^2^ analysis indicated that these NIS synthetases could accept nonnative substrates and exhibited broadened substrate scope compared to previously-reported enzymes. Notably, AcsA demonstrated approximately 8-fold higher activity than SfaB when ligating *p*-xylylenediamine (**X**) and adipic acid (**A**) to form the **XA** diad (Figures 2A and S2) while maintaining comparable activity with SfaB when synthesizing the **MA** diad from hexamethylenediamine (**M**) and adipic acid (**A**) (Figures S3 and S4). DesD displayed no higher activity than SfaB for activation of **A** but exhibited more than 7.5-fold higher activity when ligating hexamethylenediamine (**M**) and azelaic acid (**Z**) to form the **MZ** diad (Figures 2B and S5). In contrast, SfaB still exhibited the highest activity towards glutaric acid (Figure S3). These data suggested that AcsA and DesD may possess larger substrate-binding pockets than DdaG and SfaB to accommodate longer diacid (i.e., azelaic acid) and aromatic diamine (i.e., *p*-xylylenediamine) substrates.

**Figure 2.**
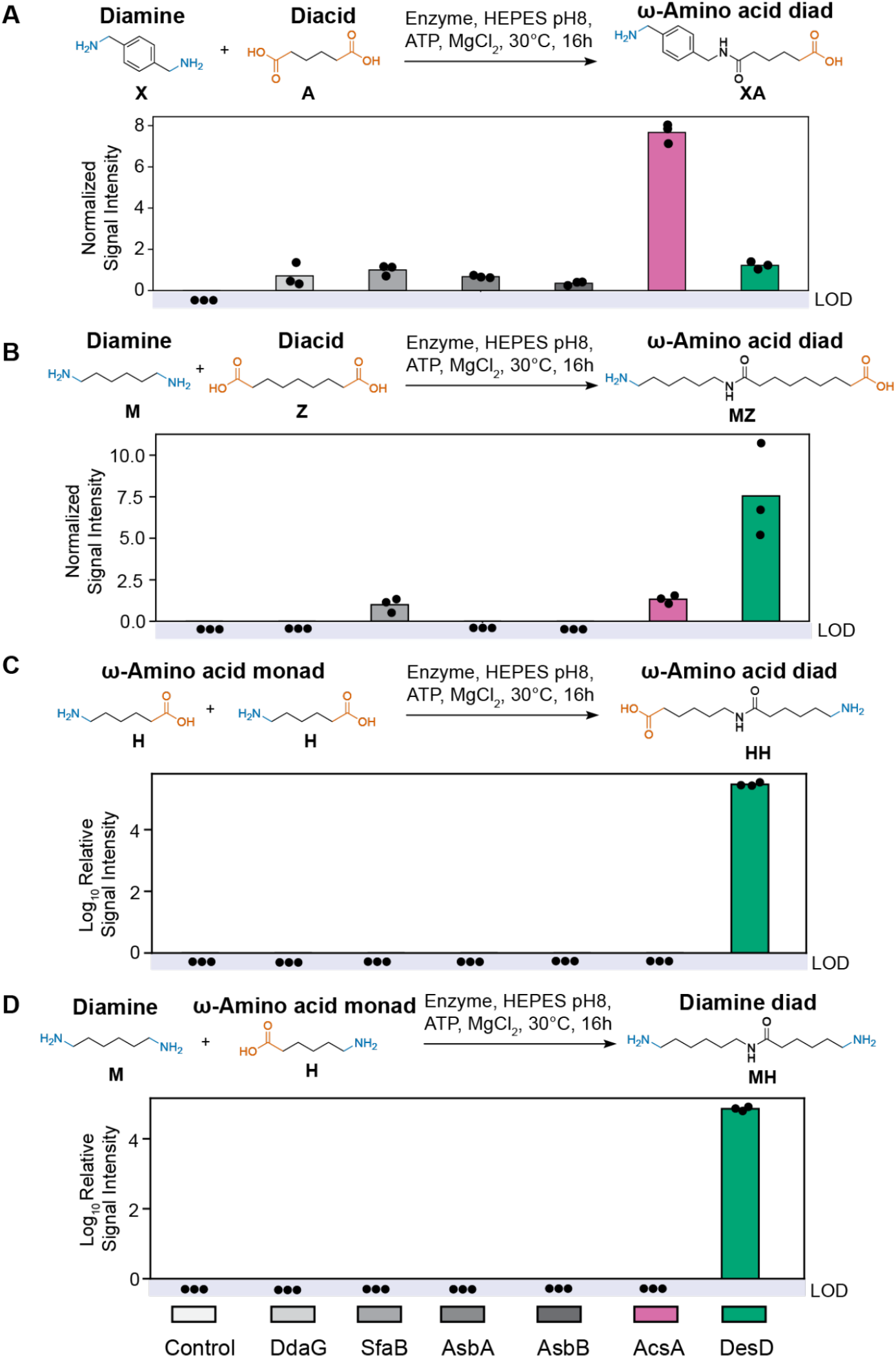
NIS synthetases can exhibit enhanced and unique diad-forming capabilities. A) *In vitro* biochemical assays were conducted by incubating **X** with **A** in the presence of enzymes (i.e., DdaG, SfaB, AsbA, AsbB, AcsA and DesD) or no-enzyme control. The MS signal intensity was normalized to SfaB. B) *In vitro* biochemical assays were conducted by incubating **M** with **Z** in the presence of enzymes (i.e., DdaG, SfaB, AsbA, AsbB, AcsA and DesD) or no-enzyme control. The MS signal intensity was normalized to SfaB. C) *In vitro* biochemical assays were conducted by incubating **H** in the presence of enzymes (i.e., DdaG, SfaB, AsbA, AsbB, AcsA and DesD) or no-enzyme control. D) *In vitro* biochemical assays were conducted by incubating **M** with **H** in the presence of enzymes (i.e., DdaG, SfaB, AsbA, AsbB, AcsA and DesD) or no-enzyme control. Products were assayed using OPSI-MS. All replicates shown are biological replicates. Sample size is *N* = 3. LOD: limit of detection. **A**: adipic acid; **X**: *p*-xylylenediamine; **Z**: azelaic acid; **M**: hexamethylenediamine; **H**: 6-aminocaproic acid.

Additionally, DesD uniquely catalyzed the dimerization of the ⍵-amino acid 6-aminocaproic acid (**H**) to the **HH** diad (Figures 2B and S6). Consistent with a previous study,^23^ we confirmed its activity for dimerization of 8-aminooctanoic acid and further demonstrated dimerization of 7-aminoheptanoic acid, whereas 5-aminovalerate was not accepted (Figures S7A and S8-9). Notably, among the enzymes tested, DesD was the only synthetase tested that formed diamine diads, for example ligating the ⍵-amino acid 6-aminocaproic acid (**H**) with the diamine hexamethylenediamine (**M**) to form the **MH** diad (Figures 2C and S10). DesD could also couple **H** or 8-aminooctanoic acid with longer diamines such as 1,8-octanediamine, but was inactive towards 5-aminovalerate with either **M** or 1,8-octanediamine (Figure S3B and S11-12). These findings suggest that DesD may possess a large binding pocket that does not efficiently accommodate shorter carboxylic acid donors.

### Structural analysis of DesD with non-native substrates

DesD displayed activity with diverse larger carboxylic acid donors, including azelaic acid (**Z**, diacid monad) and 8-aminooctanoate (⍵-amino acid monad), but was inactive towards shorter donors such as 5-aminovalerate. Therefore, we next employed structural analysis to investigate the molecular basis of this selectivity on substrate size.

For members of the adenylate-forming superfamily that carry out a similar reaction, such as acyl-CoA ligases, the substrate size-selectivity is ascribed to the size and shape of the acyl-binding pocket.^27–29^ To assess whether this principle applies to DesD, we conducted structural analysis on the structures of DesD and its homolog AcsD, which natively acts on the small substrate citrate and is the only available homolog with a ligand-bound structure.^30^ Comparing the adenyl-*N*^1^-hydroxy-*N*^1^-succinyl-cadaverine (HSC)-bound DesD structure (PDB 7TGM) and the ligand-bound structure of AcsD (PDB 2W03) revealed significant differences in the acyl-binding pocket (Figure 3A). The catalytic residues, His153/His440/Arg303 for DesD and His170/His444/Arg305 for AcsD, are conserved. Conversely, the residues defining the acyl-binding pocket differed both in size and hydrophobicity which resulted in a more hydrophobic environment in DesD. For instance, residues Asp497, Ser 556, Leu558, and Met569 in DesD are replaced by Tyr504, Lys563, Asn565 and Arg576 in AcsD, respectively, creating a smaller and more polar pocket in AcsD to accommodate binding of shorter substrates such as citrate. Pocket volume calculations showed a volume of 816 Å^3^ for AcsD and 2497 Å^3^ for DesD (Figure 3B). Superimposition of the adenosine moiety and citrate from the AcsD structure in the DesD acyl-binding pocket revealed the excess space and the lack of stabilizing contacts in DesD. Similarly, superimposition of the inhibitor HSC-AMS from the DesD structure in the AcsD structure indicated not only an excess of polar groups for a hydrophobic substrate but also the steric hindrance by Lys563 and Arg576 to prevent the binding of the substrate, with Arg576 clashing on the HSC-AMS model. These results suggested that pocket volume and polarity play a concerted role for DesD’s selectivity on substrate size.

**Figure 3.**
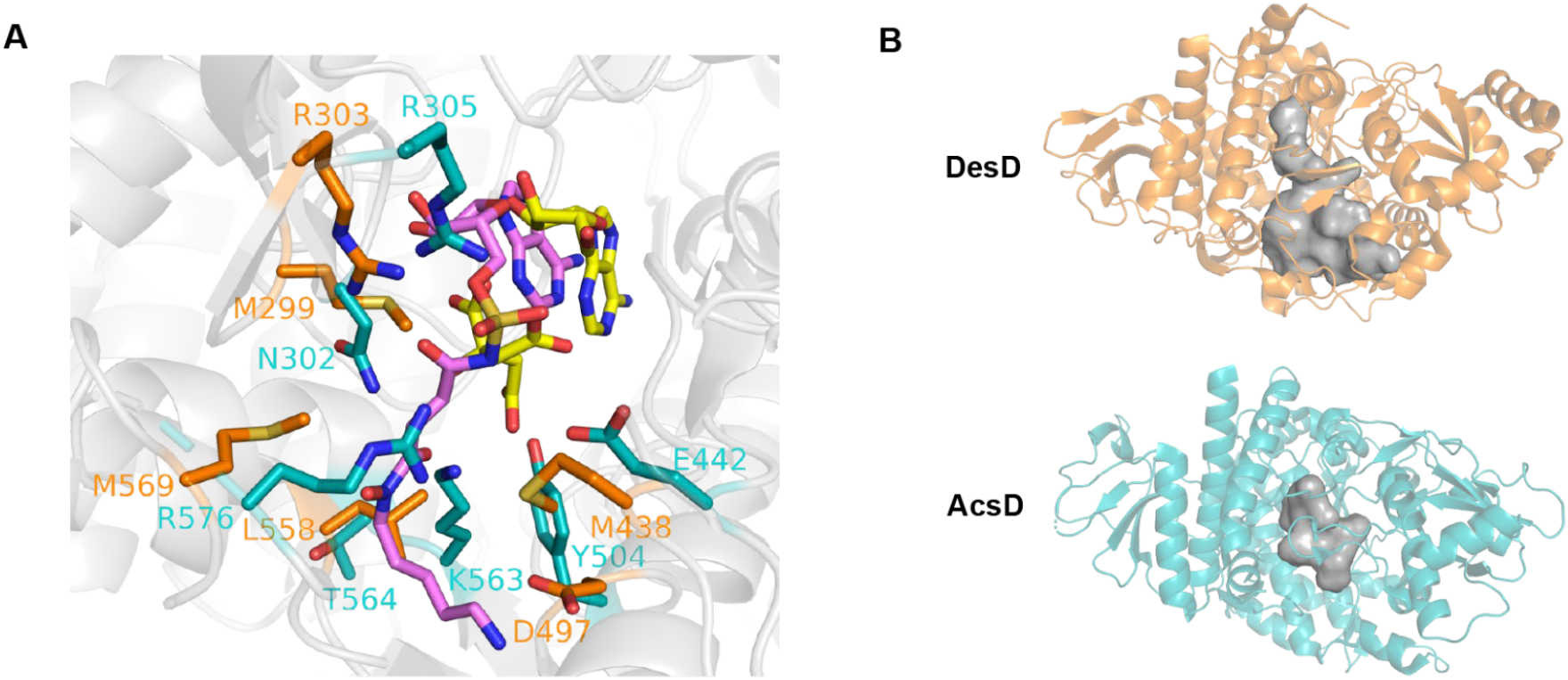
Structural comparison between DesD and its homolog AcsD. A) Comparison of active sites of DesD (orange) bound to the HSC-acyl sulfamoyl adenosine analog (HSC-AMS) (violet) and AcsD (teal) bound to AMP and citrate (yellow). B) Comparison of binding pockets of DesD and AcsD. DesD is in orange, AcsD is in teal and their pockets are shown in grey.

### Amide synthetases can synthesize longer oligoamides

While we have shown that amide synthetases can form diverse nylon-relevant diads, synthesis of novel sequenced copolyamides will require generating longer oligoamides (Figure 1). DesD is natively involved in late stages of desferrioxamine biosynthesis, where it catalyzes the formation of amide oligomers.^23^ Our results also showed that it acted on longer substrates and possessed a relatively larger substrate binding pocket. Therefore, we next tested whether amide diads can serve as substrates for triad formation. We selected two chemically synthesized diads (i.e., the polyamide-65 (PA65) and polyamide-66 (PA66) monomers, **MG** and **MA**) and tested whether the enzymes with the highest activity ligating monads (e.g. DdaG, SfaB, AcsA and DesD) could ligate these diads with other polymer-relevant substrates. Analysis by I.DOT/OPSI-MS demonstrated that both diads could be ligated with diacid monads (i.e., succinic acid (**S**), glutaric acid (**G**) and adipic acid (**A**)) to form symmetric or asymmetric diacid triads (e.g. **GMG** and **SMG**) (Figures 4A and S13-17). The enzymes also maintained their previously-described substrate preferences, with DdaG preferentially ligating **S** and SfaB preferring **G** and **A**.^12^ AcsA also formed diacid triads with all the tested diacids, exhibiting activities comparable to SfaB with **M** and **A** to SfaB, but about 1.5-fold lower activities with **S** relative to DdaG. Similarly, DesD showed no higher activities with **G** and **A** and was inactive with **S**.

**Figure 4.**
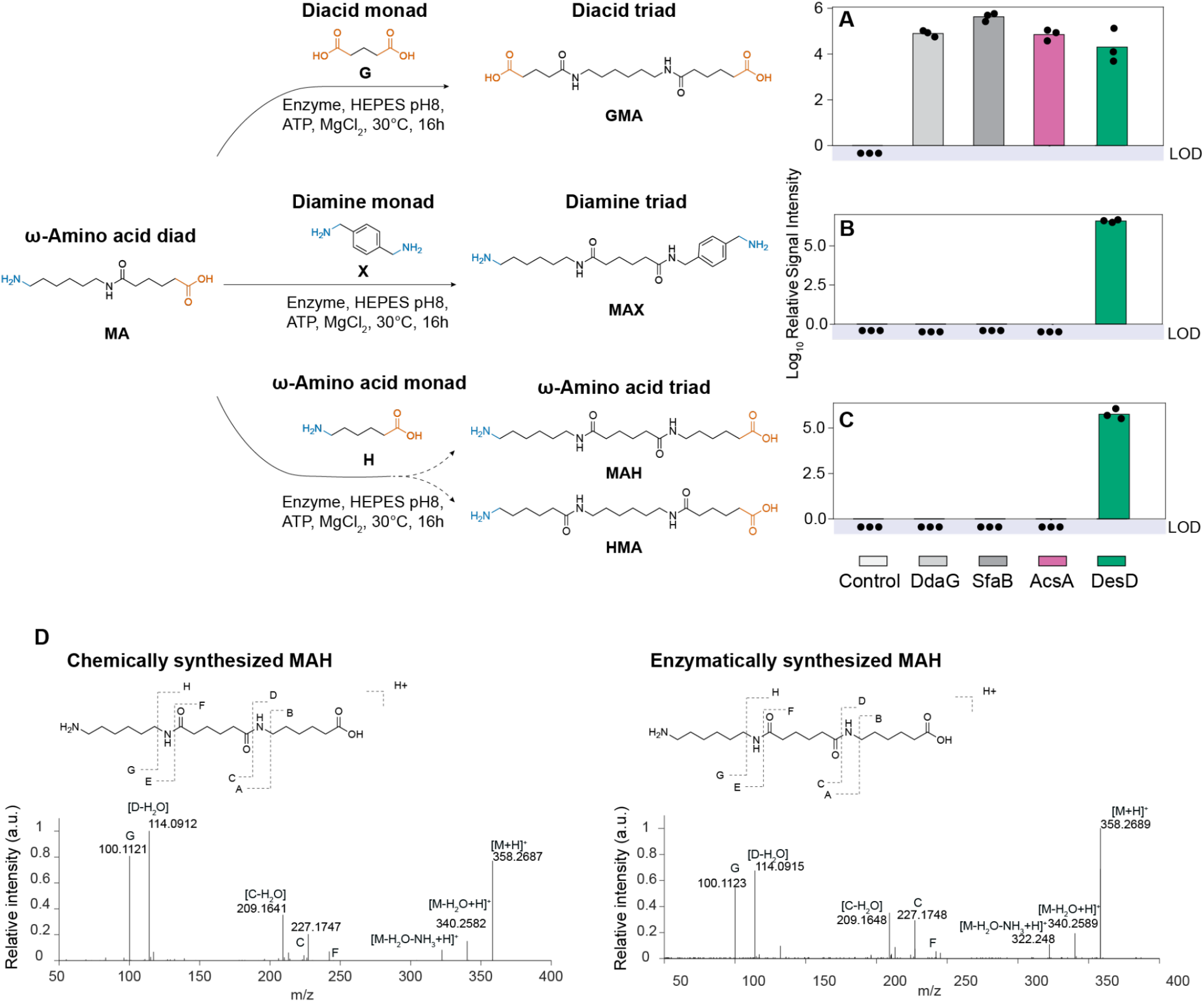
Amide synthetases can form oligotriads. *In vitro* biochemical assays were conducted by incubating ⍵-amino acid diad (i.e., **MA**) with different monads in the presence of enzymes (i.e., DdaG, SfaB, AcsA and DesD) or no-enzyme control. A) Diacid triad (i.e., **GMA**) formation between ⍵-amino acid diad (i.e., **MA)** and diacid monad (i.e., **G).** B) Diamine triads formation (i.e., **MAX**) formation between ⍵-amino acid diad (i.e., **MA**) and diamine monad (i.e., **X**). C) Two possible omega triads formation (i.e., **MAH** and **HMA**) formation between ⍵-amino acid diad (i.e., **MA**) and ⍵-amino acid monad (i.e., **H**). Products were assayed using OPSI-MS. All replicates shown are biological replicates. Sample size is *N* = 3. LOD: limit of detection. D) I.DOT/OPSI-MS^2^ (CE=25eV) of enzymatically synthesized product (right). The fragmentation spectrum closely aligns with the chemically synthesized **MAH** standard (left). **MA**: hexamethylene diamine-adipic acid; **G**: glutaric acid; **X**: *p*-xylylenediamine; **H**: 6-aminocaproic acid.

In addition, DesD exhibited distinctive activities in ligating ⍵-amino acid diads with diamine or ⍵-amino acid monads to form diamine or ⍵-amino acid triads. Specifically, using I.DOT/OPSI-MS and MS^2^, we demonstrated that DesD can ligate an ⍵-amino acid diad (i.e. the PA66 monomer, **MA**) with aliphatic (e.g., cadaverine, **C** and 1,8-octanediamine) (Figures S18-20) or aromatic diamines (e.g., *p*-xylylenediamine, **X**) to form asymmetric diamine triads (Figures 4B and S21). Interestingly, we observed that DesD could regioselectively ligate **MA** with spermidine, where it preferentially formed an amide bond at the amine group located near the secondary amine (Figure S22). However, this regioselectivity of DesD is different from that of AsbA, which favors ligation at the distal primary amine in the reaction of citrate acid and spermidine, suggesting that enzyme selection or engineering could determine product regiochemistry.

Similarly, DesD can also ligate **MA** with **H** to form an ⍵-amino acid triad (Figure 4C). When ligating an ⍵-amino acid diad with an ⍵-amino acid monad, both substrates are bifunctional and can serve as either a carboxylic donor or an amine acceptor, leading to two possible regioisomers. For example, the reaction of **MA** and **H** could yield **MAH** or **HMA**. To determine the regioselectivity of DesD in ⍵-amino acid triad synthesis, we chemically synthesized both possible triad standards, **MAH** and **HMA** (Figures S23-39), and used MS^2^ to identify their unique fragment ions. The MS^2^ spectrum of the enzymatic product matched only the **MAH** standard, indicating that DesD selectively forms a single ⍵-amino acid triad product (Figure S40). This observation aligns with the previously reported N-to-C condensation mechanism on HSC-HSC and HSC of DesD in desferrioxamine biosynthesis.^31^ The resulting sequenced **MAH** triad could potentially be polymerized into a precisely sequenced PA6/66 polymer, as an alternative to commercial block co-polymers of PA6/66 that have randomly distributed PA66 and PA6 segments.

Motivated by the success of amide triad formation, we next assessed the use of these enzymes to form amide tetrads. We chemically synthesized one symmetrical diacid triad, **AMA** (adipic acid-hexamethylenediamine-adipic acid) and two symmetrical diamine triads, **MSM** (hexamethylenediamine-succinic acid-hexamethylenediamine) and **MAM** (hexamethylenediamine-adipic acid-hexamethylenediamine) and tested for ligation with either diamine or diacid monads (Figures S41-54). We determined from I.DOT/OPSI-MS that DdaG and SfaB were capable of forming tetrads by coupling the diamine triads **MSM** or **MAM** with aliphatic diacids including **S, G**, and **A** while preserving their diacid preferences. AcsA exhibited comparable activities with SfaB, while DesD was active with **G** and **A** but not **S** (Figures 5A and S55-58). Notably, DesD and AcsA enabled the ligation of the diacid triad **AMA** with hexamethylenediamine (**M**) to form an amide tetrad (Figures 5B and S59). Since **AMA** is structurally analogous to a linear C20 diacid, this finding again suggests that DesD may possess a larger substrate binding pocket to accommodate longer substrates. Furthermore, we found that only DesD was able to dimerize the synthesized ⍵-amino acid diad (i.e., the PA66 monomer **MA**) to form ⍵-amino acid tetrad (Figures 5C and S60), highlighting its versatile potential in synthesizing complex, sequence-controlled oligoamides.

**Figure 5.**
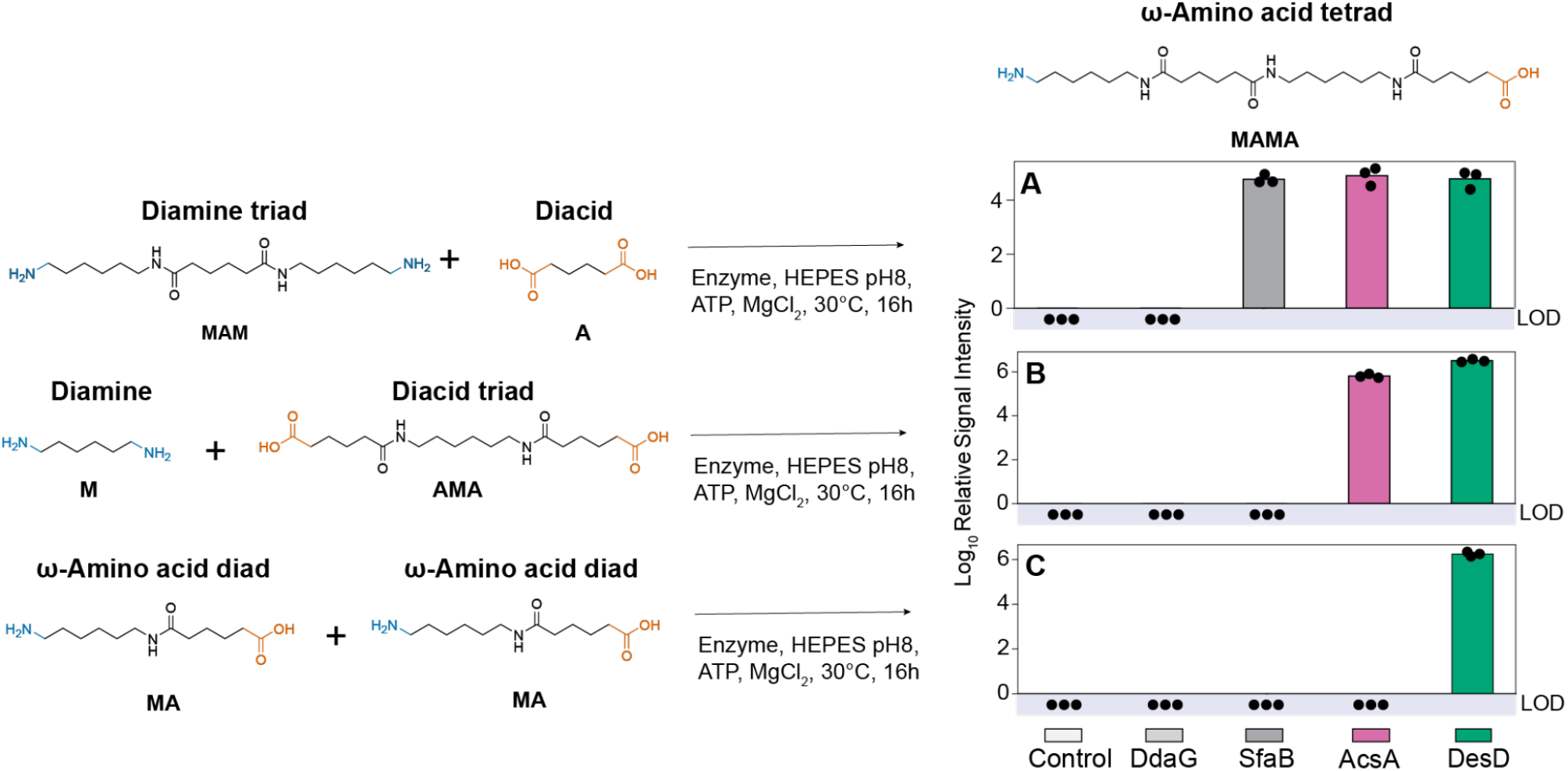
Amide synthetases can form oligotetrads. *In vitro* biochemical assays were conducted by incubating different carboxylic donors (diacid monad, diacid triad or ⍵-amino acid diad) with different amine acceptors (diamine monad, diamine triad or ⍵-amino acid diad) to synthesize the ⍵-amino acid tetrad **MAMA** in the presence of enzymes (i.e., DdaG, SfaB, AcsA and DesD) or no-enzyme control. A) Combining a diamine triad (i.e., **MAM**) with a diacid monad (i.e., **A**). B) Combining a diacid triad (i.e., **AMA**) with a diamine monad (i.e., **M).** C) Dimerization of two ⍵-amino acid diads (i.e., **MA**). Products were assayed using OPSI-MS. All replicates shown are biological replicates. Sample size is *N* = 3. LOD: limit of detection.

Since DesD can form tetrads through multiple routes, we further examined whether it could directly extend monads into longer oligoamides. We reanalyzed the reaction of DesD with **M** and **A** but did not detect any triads or tetrads (**MAMA, MAM** or **AMA**). Given that DesD exhibited more than 2-fold lower activity on **MA** production compared SfaB (Figure S3), and SfaB achieved only ∼3% conversion of this reaction,^12^ we infer that limited **MA** formation could constrain DesD’s capability to directly extend monads into higher-order oligoamides.

### Enzyme cascade enables triad formation from monads

The high catalytic activity of DdaG to ligate **S** with multiple diamines,^12^ combined with DesD’s unique ability to form diverse amide oligomers, motivated us to explore an enzymatic cascade to synthesize sequenced oligomers directly from unprotected substrates. To test the feasibility of this approach, we constructed a model two-enzyme cascade. Since DdaG can efficiently couple **S** and **M**, and DesD can ligate ⍵-amino acid diads with ⍵-amino acid monads, we conducted a simultaneous one-pot reaction to assess whether DdaG and DesD could cooperatively form an ⍵-amino acid triad from **S, M**, and 7-aminoheptanoic acid (**P**). Similar to the reaction with **MA** and **H**, when DesD ligates **MS** with **P**, two products (i.e., **MSP** and **PMS**) could be theoretically formed because both substrates are bifunctional. Consistent with DesD’s regioselectivity in synthesizing **MAH**, I.DOT/OPSI-MS and MS^2^ analysis, compared to the chemically-synthesized standard (Figures S61-68), confirmed the formation of only the target ⍵-amino acid amide triad **MSP** (Figure 6). In addition to the desired product, we also observed a range of side products formed, including **MS, MP, PS** and **SMS**, likely due to excess substrate and limited substrate specificity. This observation highlights the need for future enzyme engineering to enhance enzyme efficiency and substrate specificity towards sequence-controlled oligoamide synthesis. Nevertheless, this simultaneous cascade provides an enzymatic pathway for a protecting-group-free biocatalytic route to diverse oligoamides from polymer-relevant substrates.

**Figure 6.**
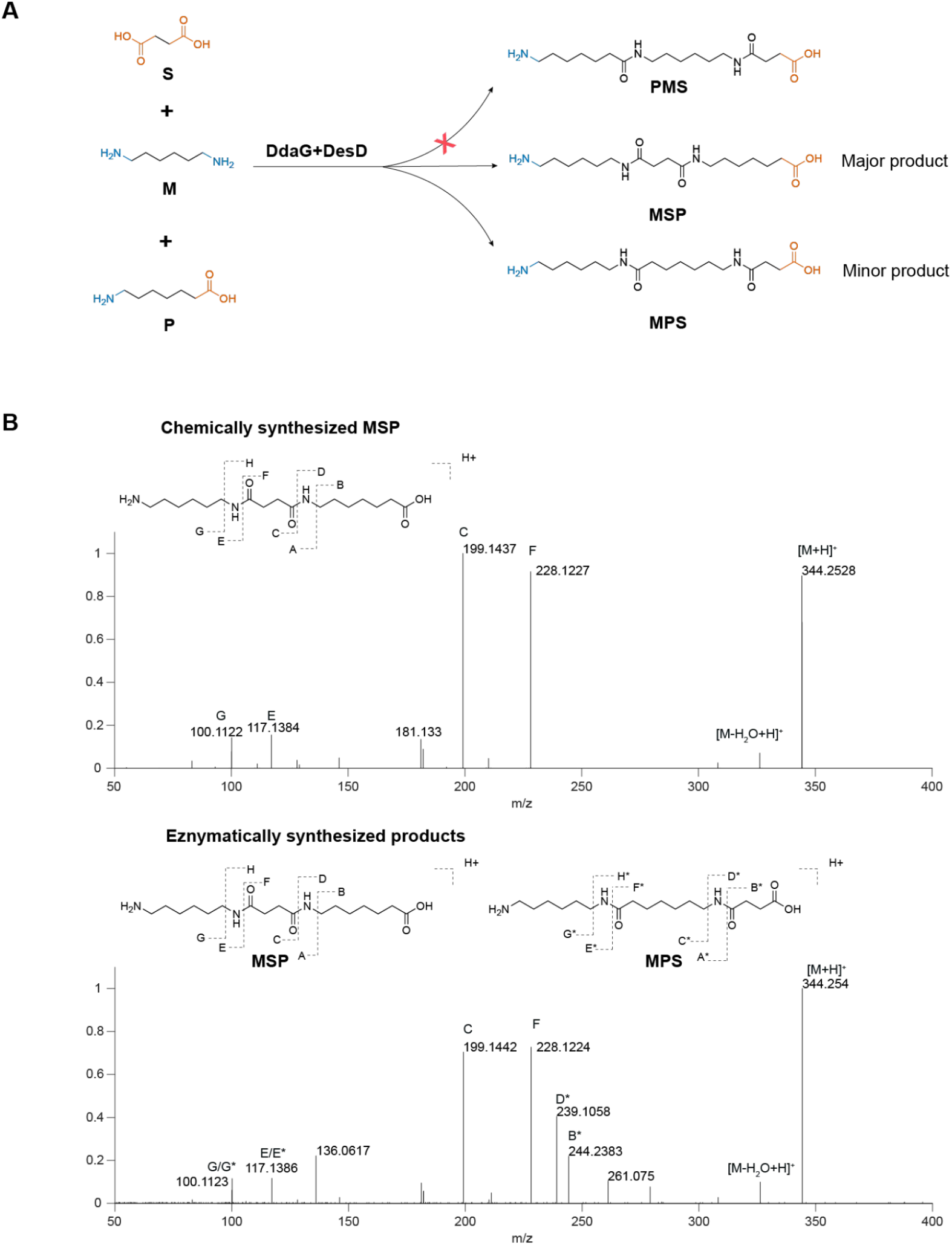
Enzyme cascade can synthesize ⍵-amino acid triad from unprotected monomers. A) One-pot simultaneous reaction of **S, M**, and **P** with DdaG and DesD forming a single ⍵-amino acid triad. Of the two possible ligation products **MSP** and **PMS**, only **MSP** was detected. B) MS^2^ spectrum (30 eV) comparison between enzymatically synthesized products and chemically synthesized standard. **S**: succinic acid; **M**: hexamethylenediamine; **P**: 7-aminoheptanoic acid.

**Figure 7.**
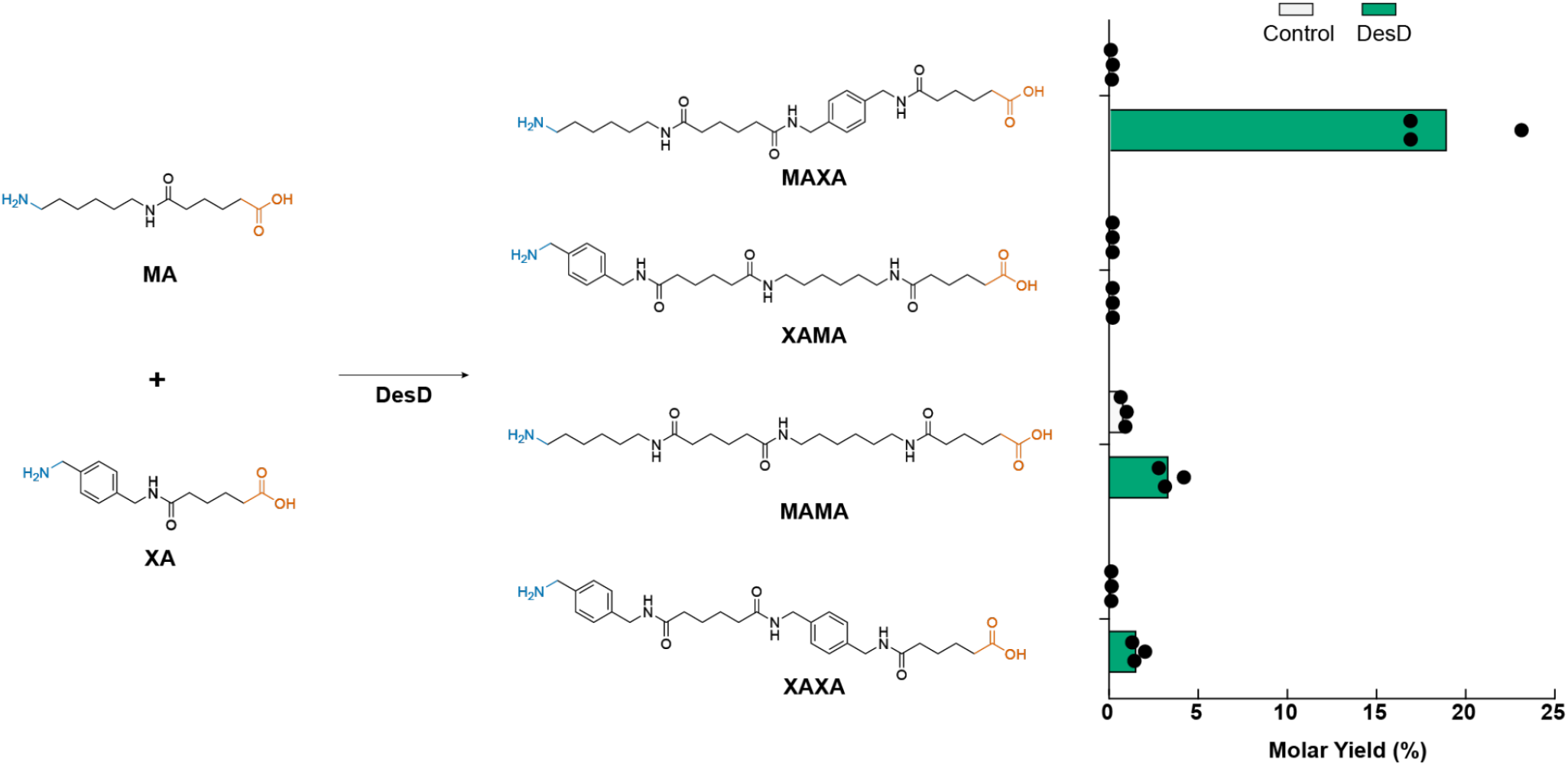
DesD regioselectively synthesizes oligotetrads from asymmetric ω-amino diads for sequence-defined polyamide production. *In vitro* biochemical assays were conducted by incubating **XA** and **MA** in the presence of DesD or no-enzyme control. Products were quantified by calibration with chemically-synthesized standards using I.DOT/OPSI-MS. Sample size is N = 3.

### MAXA synthesis

Novel sequence-defined polyamides can exhibit superior material properties compared to conventional nylons. For example, the recently described poly(hexamethylenediamine—adipic acid—*p*-xylylenediamine—adipic acid), also described as poly[**MAXA]**, displayed a higher melting temperature and reduced water uptake compared to PA66. Since DesD has been observed to dimerize ω-amino acid diads (i.e., **MA** and **HH**) into homotetrads (i.e., **MAMA** and **HHHH**), we next examined its activity to ligate the chemically synthesized standards **MA** (hexamethylenediamine—adipic acid) and **XA** (*p*-xylylenediamine—adipic acid) (Figures S69-76). Remarkably, DesD catalyzed this ligation with high product regioselectivity. Because both **MA** and **XA** contain carboxylic acid and amine termini, the ligation products could contain a mixture of heterotetrads (i.e., **MAXA** and/or **XAMA)** and homotetrads (i.e., **MAMA** and/or **XAXA**). However, by comparison to chemically-synthesized standards (Figures S77-90), the yield of **MAXA** was 15%, while only a trace amount of **XAMA** was detected (0.009%). In addition to **MAXA**, low levels of the homotetrads **MAMA** (3%) and **XAXA** (2%) were also detected; however, their yields were significantly lower than that of **MAXA**.

Motivated by this finding, we next assembled an *in vitro* enzymatic cascade using AcsA, SfaB, and DesD to synthesize **MAXA** directly from monads, leveraging AcsA to synthesize **XA**, SfaB to synthesize **MA**, and DesD to synthesize **MAXA**. However, this cascade did not generate detectable amounts of **MAXA**. Quantification of separate reactions showed that SfaB produced **MA** at a yield of 6% and AcsA produced **XA** at a yield of 1%, indicating that the cascade failure was likely due to the low production of these diads and highlighting the need for future enzyme engineering. Nevertheless, the selective enzymatic synthesis of **MAXA** demonstrates the potential of biocatalysts to efficiently generate sequence-defined oligoamides for advanced polymers.

## Conclusion

In summary, our study demonstrated that NIS synthetases, represented by DesD, can regioselectively ligate oligomeric substrates to sequenced oligoamides. Using an enzyme cascade, we directly assembled an amide triad from unprotected monomers, bypassing the need for protecting functional groups or multistep chemical synthesis. This biocatalytic approach offers a practical route to sequence-defined oligoamides and expands the diversity of accessible polyamide sequences beyond conventional nylons. Moreover, our findings provide a foundation for exploring sequence–property relationships in polymers and advancing the design of tailored polyamides.

## Acknowledgements

Research was sponsored by the Laboratory Directed Research and Development Program of Oak Ridge National Laboratory, managed by UT-Battelle, LLC under Contract No. DE-AC05-00OR22725, for the U. S. Department of Energy. LQ, ITD, JCF, and JKM are inventors on a patent application related to this work. We thank Stephen J. Koehler and C. Adrian Figg for providing the chemically synthesized MAMA standard.

## Author contributions

L.Q: Biochemistry, writing

I.T.D.: Synthesis of analytical standards

N.C.: Structural analysis, docking simulations, writing

D.L.C.: Analytical characterization

N.O.: Synthesis of analytical standards

V.K.: Analytical methods development and analysis

J.F.C.:Analytical methods development and analysis, supervision of analytical research.

J.C.F.: Conceptualization, supervision of chemistry research

J.K.M.: Conceptualization; funding acquisition; supervision of biochemistry research, writing.

## Methods

### Materials

LB agar, LB medium and Terrific Broth were purchased from BD Difco. All chemicals were purchased from ThermoFisher, Alfa Aesar, or Tokyo Chemical Industry (TCI).

### Synthesis and expression of enzymes

Genes encoding AsbA, AsbB, AcsA and DesD were synthesized by GenScript (Piscataway, NJ) with a N-terminal His_6_ tag for purification and cloned into pET28a(+). Plasmids were transformed into chemically competent BL21 (DE3) *Escherichia coli* cells (Sigma-Aldrich, St Louis, MO) according to the manufacturer’s specifications.

Expression and purification of His-DdaG, His-SfaB, His-AsbA, His-AsbB, His-AcsA and His-DesD were conducted following previously published methods.^32^ A colony or glycerol stock of BL21(DE3) *E. coli* containing plasmid DNA was used to inoculate 3 mL of LB medium supplemented with appropriate antibiotics. The culture was incubated at 37 ℃ and 250 rpm overnight. The overnight culture was then diluted 100-fold into 200 mL of Terrific Broth. The subculture was incubated at 37 °C at 250 rpm until an OD of 0.4-0.6 was reached. Protein expression was induced by adding IPTG to a final concentration of 0.2 mM. The temperature was decreased to 20 °C and the culture was grown with shaking overnight. The cells were harvested by centrifugation (4000 rpm, 40 min, 4 °C), resuspended in 5 mL of lysis buffer (50 mM HEPES, 300 mM NaCl, 10 mM imidazole, 10mM MgCl_2_, 10% glycerol, pH 8.0), and sonicated on ice. Cell debris was removed by centrifugation (10,000 rpm, 30 min, 4 °C). The supernatant was then filtered with an MCE filter (0.22 μm) and loaded at 3 mL/min onto an ÄKTA Start FPLC (Cytiva Marlburough, MA) equipped with a 5-mL His-Trap™ column (Cytiva). The column was washed with wash buffer (50 mM HEPES, 300 mM NaCl, 30 mM imidazole, 10 mM MgCl_2_, 10% glycerol, pH 8.0) before eluting with elution buffer (50 mM HEPES, 300 mM NaCl, 250 mM imidazole, 10 mM MgCl_2_, 10% glycerol, pH 8.0) at 3 mL/min. Purified proteins were concentrated with an Amicon® Ultra centrifugal filter at 10 kDa cutoff, and buffer exchanged into exchange buffer (50 mM HEPES, pH 8.0, 100 mM NaCl, 10mM MgCl_2_, 10% glycerol). The final purified product was analyzed by SDS-PAGE and enzyme concentration was measured using a NanoDrop™ 1000 Spectrophotometer (Thermo Scientific) with exchange buffer as a blank, and the extinction coefficient of each protein was calculated using ProtParam from the ExPASy Proteomics Server. The molecular weight of each protein was calculated using ProtParam from the ExPASy Proteomics Server, and further confirmed by the SDS-PAGE method. Pure enzymes were stored at −80 °C for subsequent activity assays.

### In vitro enzyme activity assays

A typical enzymatic reaction was carried at 50 μL containing 100 mM HEPES (pH 8.0), 10 mM ATP, 10 mM MgCl_2_, 10 μM enzyme, 5 mM carboxylic group-containing compounds and 5 mM amine group-containing compounds and was incubated at 30 °C for 16 h with shaking at 200 rpm. The reaction was quenched by adding 50 μL methanol. The samples were centrifuged at 4,000 rpm for 10 min, and the supernatant was subjected to MS analysis to determine the formation of corresponding products.

### Enzyme cascade assays

For one pot synthesis, the reaction was carried at 100 μL containing 100 mM HEPES (pH 8.0), 40 mM ATP, 40 mM MgCl_2_, 10 μM DdaG, 20 μM DesD, 5 mM succinic acid, 5 mM hexamethylenediamine, and 5 mM 1,8-octanoic amino acid and was incubated at 30

°C for 16 h with shaking at 200 rpm. The reaction was quenched by adding 100 μL methanol. The samples were centrifuged at 4,000 rpm for 10 min, and the supernatant was subjected to MS analysis to determine the formation of corresponding products.

### I.DOT/OPSI-MS analysis of enzyme activity

The immediate drop-on-demand technology (Dispendix GmbH, Stuttgart, Germany) coupled with open port sampling interface mass spectrometry (I.DOT/OPSI-MS) was used for targeted quantification, untargeted analysis by mass, and collection of tandem mass spectra.^12,24,33^ All reactions were diluted 1:100 (v/v) in HPLC grade water containing 500 nM propranolol which served as an internal standard for droplet detection and droplet normalization. 40 µL of diluted reactions were transferred to I.DOT S.100 96-well plates and analyzed immediately. For analysis, 20 nL of sample was ejected into the OPSI-MS having a flow of 75/25/0.1 (v/v/v) acetonitrile/water/formic acid. Samples were transported to the electrospray ion source of a Sciex 7500 triple quadrupole mass spectrometer (SCIEX, Concord, Ontario, Canada) for targeted quantitation of oligotetrads. Quantification used a 12-point calibration curve for each oligotetrad (0-500 µg/mL). Each sample was measured in triplicate, with the means averaged across sample replicates for reported means and standard deviations. The 7500 was operated in positive ion mode with flow = 150 µL/min, gas setting 1 = 90, gas setting 2 = 60, electrospray voltage = 5.5 kV, capillary temperature = 200 °C, and dwell time = 20 ms. Multiple reaction monitoring (MRM) used the following transitions: 260.1→183.0; collision energy (CE)=26 eV (propranolol), 491.32→248.00; CE=50 eV (MAXA), 491.32→337.00; CE=31 eV (XAMA), 471.35→100.10; CE=24 eV (MAMA), and 511.29→248.13; CE=35 eV (XAXA). In-house developed softwares (ORNL I.DOT-MS Coupler v2.50) were used for control of the IDOT system, extraction of data from vendor file formats, peak finding, and peak integration. Each droplet signal was background subtracted and normalized to propranolol signal. For untargeted and tandem mass spectrometry analyses, a Thermo Q-Exactive HF mass spectrometer (ThermoFisher Scientific) with sheath gas = 80, auxiliary gas = 40, electrospray voltage = 4 kV, ion injection time = 50 ms, automatic gain control = 3e^6^, capillary temperature = 200 °C, mass/charge (*m/z*) range = 100-1000 *m/z*, and OPSI solvent flow of 250 µL/min was used in positive ion mode with a mass resolution of 60,000. Tandem mass spectra were acquired using the same system, with a 1 *m/z* isolation window, 30,000 mass resolution, and CEs between 10-80 eV. Putative fragment identifications are given for each compound using hypothesized fragmented pathways. Additional CEs were used to help with fragment identifications as necessary (data not shown).

#### Pocket volume calculation

Pocket volumes were calculated with the PyMOL plugin PyVOL^34^, increasing the minimum radius to 1.8 Å and decreasing the maximum to 3.0 Å. The search was restricted to the binding pocket, including the AMP-binding pocket, by selecting the residues up to 5 Å away from the ligands, which were then omitted. Figures were generated with PyMOL^35^.

