## Supplemental Information for "Regioselective biosynthesis of oligoamides as precursors for sequence-controlled co-polyamides"

**Notice: This manuscript has been authored by UT-Battelle, LLC under Contract No. DE-AC05-00OR22725 with the U.S. Department of Energy. The United States Government retains and the publisher, by accepting the article for publication, acknowledges that the United States Government retains a non-exclusive, paid-up, irrevocable, world-wide license to publish or reproduce the published form of this manuscript, or allow others to do so, for United States Government purposes. DOE will provide public access to these results of federally sponsored research in accordance with the DOE Public Access Plan (http://energy.gov/downloads/doe-public-access-plan).**

**Chemical synthesis materials and methods:**

All chemicals were purchased from commercial sources and used without further purification unless otherwise stated. Methyl 6-((6-*N*-boc-aminohexyl)amino)-6-oxohexanoate (Boc-MA-OMe), 6-(6-*N*-boc-aminohexyl)amino)-6-oxohexanoic acid (Boc-MA), and 4-((6-*N*-boc-amino)hexyl)amino)-4-oxobutanoic acid (Boc-MS) were prepared as described previously.[1]

*Nuclear magnetic resonance (NMR) spectroscopy.* $^1$H and $^{13}$C NMR spectra were recorded on a Bruker Avance III spectrometer operating at 400 MHZ ($^1$H) or 100 MHz ($^{13}$C). Chemical shifts are reported in parts per million (ppm) relative to residual protonated solvent.

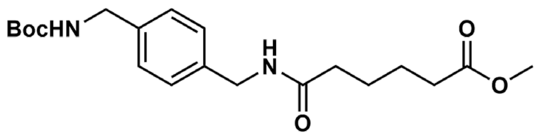


*Methyl 6-((4-(((t-butoxycarbonyl)amino)methyl)benzyl)amino)-6-oxohexanoate (Boc-XA-OMe).* A 1000 mL rbf was charged with monomethyl adipate (10 g, 62.4 mmol, 1 equiv.) and 450 mL dichloromethane (DCM). 1-Ethyl-3-(3-dimethylaminopropyl)carbodiimide hydrochloride (EDC·HCl) (13.16 g, 68.6 mmol, 1.1 equiv.) was added and the RBF was stirred at r.t. for 5 min until dissolved. 1-(*N*-Boc-aminomethyl)-4-(aminomethyl)benzene (16.22 g, 68.6 mmol, 1.1 equiv) was then added to the reaction mixture and the reaction was stirred at r.t. overnight. The reaction was washed with 2×250 mL 1 M HCl, 3×250 mL mL saturated $NaHCO_3$ solution, and 250 mL brine. The organic layer was dried over sodium sulfate, filtered, and concentrated using a rotary evaporator. The product (14.43 g, 61.1%) was dried overnight in a vacuum oven and used in subsequent reactions without further purification. $^1$H NMR (400 MHz, DMSO-$d_6$) δ 8.26 (t, 1H), 7.35 (t, J = 6.3 Hz, 1H), 7.16 (s, 4H), 4.21 (d, J = 5.9 Hz, 2H), 4.08 (d, J = 6.2 Hz, 2H), 3.58 (s, 3H), 2.30 (s, 2H), 2.12 (s, 2H), 1.52 (d, J = 3.6 Hz, 4H), 1.38 (s, 9H). $^{13}$C NMR (101 MHz, DMSO-$d_6$) δ 173.7, 172.2, 156.2, 139.1, 138.5, 127.6, 127.3, 78.2, 51.7, 43.6, 42.2, 35.4, 33.5, 28.7, 25.2, 24.6.

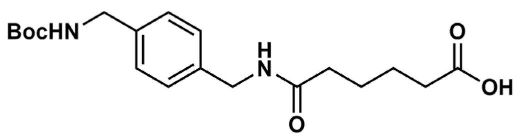


*6-((4-(((t-Butoxycarbonyl)amino)methyl)benzyl)amino)-6-oxohexanoic acid (Boc-XA).* In a 500 mL round bottom flask (RBF), Boc-XA-OMe (5 g, 13.2 mmol) was dissolved in 50 mL THF. The RBF was charged with a 1 M LiOH solution (60 mL) and subsequently stirred at r.t. for 3.5 h. The reaction was acidified to pH ~4 by addition of 1 M HCl. THF was removed *in vacuo*, and the resulting white precipitate was filtered and dried under vacuum to provide the product as a white solid (4.54 g, 94%). $^1$H NMR (400 MHz, DMSO-$d_6$) δ 11.99 (s, 1H), 8.26 (t, J = 5.9 Hz, 1H), 7.35 (t, J = 6.2 Hz, 1H), 7.17 (s, 4H), 4.21 (d, J = 5.8 Hz, 2H), 4.08 (d, J = 6.3 Hz, 2H), 2.50 (p, J = 1.8

Hz, 4H), 2.21 (t, J = 6.9 Hz, 2H), 2.12 (t, J = 6.9 Hz, 2H), 1.50 (td, J = 12.4, 6.0 Hz, 4H), 1.38 (m, 9H). $^{13}$C NMR (101 MHz, DMSO-d$_6$) δ 174.9, 172.3, 156.2, 139.1, 138.5, 127.6, 127.4, 78.2, 43.6, 42.2, 35.5, 33.9, 28.728, 25.3, 24.6.

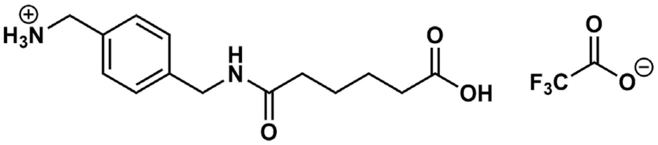

*(4-((5-Carboxypentanamido)methyl)phenyl)methanaminium trifluoroacetate (XA trifluoroacetate).* In a 100 mL RBF, suspended Boc-XA (1.5 g, 4.12 mmol) in DCM (25 mL). Added trifluoroacetic acid (10 mL) and stirred at r.t. for 3 h. The majority of trifluoroacetic acid was removed by concentrating the reaction *in vacuo* and diluting with DCM for 3 cycles. After the third cycle, the concentrated crude was precipitated into diethyl ether to form a white solid. The solid was filtered and dried under vacuum to provide XA trifluoroacetate (1.52 g, 98%). $^1$H NMR (400 MHz, DMSO-d$_6$) δ 12.02 (s, 1H), 8.36 (t, $J$ = 6.1 Hz, 1H), 8.18 (s, 3H), 7.85 – 6.88 (dd, 4H), 4.25 (d, $J$ = 6.0 Hz, 2H), 4.14 – 3.87 (m, 2H), 2.21 (t, $J$ = 6.7 Hz, 2H), 2.14 (t, $J$ = 6.9 Hz, 2H), 1.51 (h, $J$ = 7.1 Hz, 4H). $^{13}$C NMR (101 MHz, DMSO-d$_6$) δ 174.4, 172.0, 157.9, 140.3, 132.3, 128.8, 127.3, 118.6, 64.9, 42.0, 41.6, 35.0, 33.4, 24.8, 24.2, 15.2.

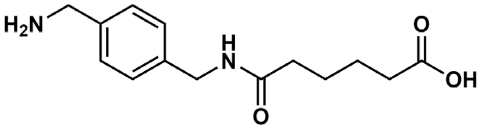

*6-((4-(Aminomethyl)benzyl)amino)-6-oxohexanoic acid (XA).* In a 100 mL RBF, dissolved in XA trifluoroacetate (1.25 g, 3.35 mmol, 1 equiv.) in DI $H_2O$ (20 mL). Heated reactor to 50 °C and added Reillex 402 (1.76 g, 5 equiv). Equipped RBF with reflux condenser and stirred at 50 °C overnight. The reaction was then filtered. The filtrate was concentrated using a rotary evaporator, and the residue was precipitated by addition of EtOH to provide the product as a flaky white solid (0.34 g, 39%). $^1$H NMR (400 MHz, $D_2O$) δ 7.63 – 7.20 (m, 4H), 4.40 (s, 2H), 4.18 (s, 2H), 2.33 (t, $J$ = 7.2 Hz, 2H), 2.17 (t, $J$ = 7.4 Hz, 2H), 1.62 (p, $J$ = 6.8 Hz, 2H), 1.53 (q, $J$ = 7.3 Hz, 2H). $^{13}$C NMR (101 MHz, $D_2O$) δ 183.3, 177.0, 139.2, 131.5, 129.1, 127.8, 42.8, 42.6, 37.1, 35.5, 25.3, 25.2.

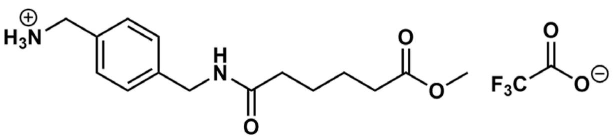

*(4-((6-Methoxy-6-oxohexanamido)methyl)phenyl)methanaminium trifluoroacetate (XA-OMe).* In a 500 mL RBF, Boc-XA-OMe (14 g, 37 mmol) was dissolved in DCM (210 mL). The RBF was chilled to 0 °C in an ice bath and trifluoroacetic acid (70 mL) was added slowly to the reaction. The flask was stirred overnight, warming gradually to room temperature. The majority of

trifluoroacetic acid was removed by concentrating the reaction in vacuo and diluting with DCM for 3 cycles. After the third cycle, the concentrated crude was precipitated into diethyl ether causing a white solid to form. The solid was filtered and dried under vacuum to provide the product in quantitative yield. $^{1}$H NMR (400 MHz, DMSO-d$_6$) δ 8.67 – 7.86 (m, 4H), 7.38 (d, J = 7.8 Hz, 2H), 7.27 (d, J = 7.8 Hz, 2H), 4.26 (d, J = 6.0 Hz, 2H), 4.00 (s, 2H), 3.58 (s, 3H), 2.40 – 2.23 (m, 2H), 2.14 (t, J = 4.3 Hz, 2H), 1.52 (p, J = 3.6 Hz, 4H). $^{13}$C NMR (101 MHz, DMSO-d$_6$) δ 173.7, 172.4, 158.4, 140.7, 132.8, 129.3, 127.8, 51.7, 42.5, 42.1, 35.4, 33.5, 25.2, 24.6.

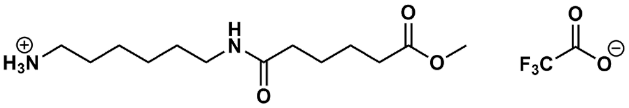


*6-(6-Methoxy-6-oxohexanamido)hexan-1-aminium trifluoroacetate (MA-OMe).* In a 250 mL RBF, Boc-MA-OMe (3.5 g, 9.76 mmol) was dissolved in DCM (50 mL). The RBF was chilled to 0 °C in an ice bath and trifluoroacetic acid (25 mL) was added slowly to the reaction. The flask was stirred overnight, warming gradually to room temperature. The majority of trifluoroacetic acid was removed by concentrating the reaction in vacuo and diluting with DCM for 3 cycles. After the third cycle, the concentrated crude was precipitated into diethyl ether causing a white solid to precipitate. The solid was filtered and dried under vacuum to provide the product (3.22 g, 88%). $^{1}$H NMR (400 MHz, DMSO-d$_6$) δ 8.09 – 7.29 (m, 4H), 3.58 (s, 3H), 3.02 (q, J = 6.6 Hz, 2H), 2.77 (q, J = 6.8 Hz, 2H), 2.29 (d, J = 6.7 Hz, 2H), 2.06 (d, J = 6.8 Hz, 2H), 1.66 – 1.42 (m, 6H), 1.37 (q, J = 7.1 Hz, 2H), 1.32 – 1.14 (m, 4H). $^{13}$C NMR (101 MHz, DMSO-d$^{6}$) δ 173.3, 171.6, 51.2, 38.8, 38.2, 35.0, 33.0, 29.0, 26.9, 25.9, 25.5, 24.8, 24.0.

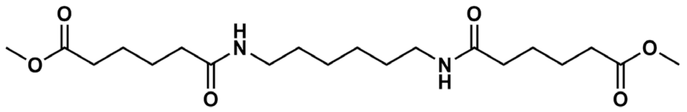


*1,1'-Dimethyl 6,6'-(1,6-hexanediyldiimino)bis[6-oxohexanoate] (AMA-OMe).* A 500 mL round-bottom flask (rbf) equipped with a stir bar was charged with monomethyl adipate (12.13 g, 75.7 mmol, 2.2 equiv.) and DCM (350 mL). To this was added 1-Ethyl-3-(3-dimethylaminopropyl)carbodiimide (14.52 g, 75.7 mmol, 2.2 equiv.) and the rbf was stirred at r.t. for 15 min. The flask was subsequently charged with hexamethylenediamine (4 g, 34.4 mmol, 1 equiv.) and the reaction was stirred at room temperature overnight. The crude reaction mixture was subsequently washed twice with 250 mL 1 M HCl, three times with 250 mL of a saturated solution of sodium bicarbonate, and once with 250 mL brine. The DCM layer was dried over sodium sulfate and concentrated under vacuum to provide AMA-OMe as a flaky white solid (7.55 g, 55%). $^{1}$H NMR (400 MHz, $CDCl_3$) δ 5.76 (s, 2H), 3.24 (q, $J$ = 6.0 Hz, 4H), 2.35 (dt, $J$ = 6.9, 3.4 Hz, 4H), 2.20 (q, $J$ = 5.1 Hz, 4H), 1.66 (h, $J$ = 2.9 Hz, 8H), 1.50 (t, $J$ = 6.9 Hz, 4H), 1.34 (dt, $J$ = 7.0, 4.0 Hz, 4H). $^{13}$C NMR (101 MHz, $CDCl_3$) δ 174.01, 172.78, 51.57, 39.01, 36.21, 33.68, 29.37, 25.93, 25.17, 24.40.

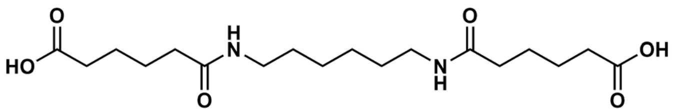

*6,6'-(1,6-Hexanediyldiimino)bis[6-oxohexanoic acid] (AMA).* In a 250 mL rbf, AMA-OMe (7 g, 17.5 mmol) was dissolved in 70 mL THF. The mixture was charged with a solution of 1 M LiOH (70 mL) and stirred at room temperature for 3.5 h. The reaction was acidified to ~pH 4 with 1 M HCl, and the THF was removed under vacuum. The precipitate was filtered and dried under vacuum to provide AMA as a fine white powder (6.51 g, 92%). $^1$H NMR (400 MHz, DMSO) δ 11.99 (s, 2H), 7.73 (t, $J$ = 5.7 Hz, 2H), 3.01 (q, $J$ = 6.5 Hz, 4H), 2.20 (t, $J$ = 6.6 Hz, 4H), 2.12 – 1.90 (m, 4H), 1.48 (tt, $J$ = 7.7, 3.9 Hz, 8H), 1.35 (q, $J$ = 6.8 Hz, 4H), 1.24 (h, $J$ = 4.7 Hz, 4H). $^{13}$C NMR (101 MHz, DMSO) δ 174.27, 171.55, 38.20, 35.00, 33.30, 29.01, 26.01, 24.76, 24.03.

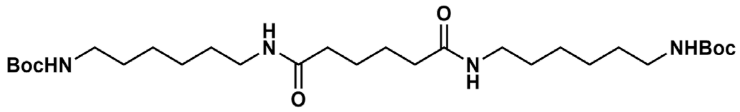

*di-t-butyl ((adipoylbis(azanediyl))bis(hexane-6,1-diyl))dicarbamate(Boc-MAM).* In a 500 mL RBF, EDC·HCl (10.49 g, 54.8 mmol, 2 equiv.) was added to a solution of adipic acid (4 g, 27.4 mmol, 2 equiv) dissolved in DCM (275 mL). *N*-Boc-1,6-diaminohexane (11.84 g, 54.8 mmol, 1 equiv.) was added the RBF, and the reaction was stirred at r.t. overnight. A white precipitate was formed over the course of the reaction which was filtered, rinsed with additional DCM and DI $H_2O$, and dried under vacuum to provide the product (12.4 g, 83.4%). $^1$H NMR (400 MHz, DMSO-$d_6$) δ 7.70 (t, J = 5.6 Hz, 2H), 6.75 (t, J = 5.7 Hz, 2H), 2.99 (q, J = 6.6 Hz, 4H), 2.88 (q, J = 6.6 Hz, 4H), 2.12 – 1.94 (m, 4H), 1.43 (p, J = 3.5 Hz, 4H), 1.36 (s, 25H), 1.21 (p, J = 3.6 Hz, 8H). $^{13}$C NMR (101 MHz, DMSO-$d_6$) δ 171.7, 155.6, 77.3, 38.3, 35.3, 29.4, 29.1, 28.3, 26.1, 26.0, 25.1.

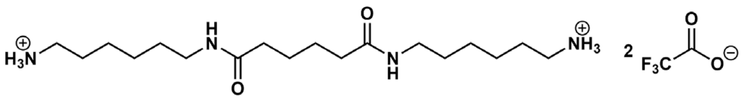

*6,6'-(adipoylbis(azanediyl))bis(hexan-1-aminium) trifluoroacetate (MAM trifluoroacetate).* In a 250 mL RBF, trifluoroacetic acid (25 mL) was added to a solution of Boc-MAM (5 g, 9.21 mmol) dissolved in DCM (75 mL). The majority of trifluoroacetic acid was removed by concentrating the reaction *in vacuo* and diluting with DCM for 3 cycles. After the third cycle, the concentrated crude was precipitated into diethyl ether. The resulting residue was dissolved in MeOH, transferred to a scintillation vial, and concentrated via rotary evaporation to provide the product as a highly viscous, colorless liquid (5.26 g, 80.0%). $^1$H NMR (400 MHz, DMSO-$d_6$) δ

8.32 – 7.40 (m, 8H), 3.01 (q, J = 6.6 Hz, 4H), 2.87 – 2.68 (m, 4H), 2.03 (td, J = 5.5, 2.4 Hz, 4H), 1.51 (p, J = 7.4 Hz, 4H), 1.44 (p, J = 3.6 Hz, 4H), 1.37 (p, J = 7.1 Hz, 4H), 1.26 (dp, J = 11.3, 6.4 Hz, 8H). $^{13}$C NMR (101 MHz, DMSO-d$_6$) δ 171.8, 158.3, 99.7, 48.6, 38.8, 38.2, 35.3, 29.0, 26.9, 25.9, 25.5, 25.1.

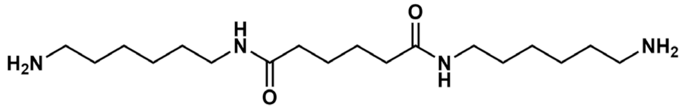


*N*[1],*N*[6]-bis(6-aminohexyl)adipamide (MAM). In a 500 mL RBF, dissolved MSM trifluoroacetate (6.5 g, 11.4 mmol) in MeOH (350 mL). Added Amberlyst A-21 ion exchange resin (7.42 g, 3 equiv.) and stirred at r.t. overnight. The reaction mixture was filtered and concentrated via rotary evaporation to provide a residue that was further purified via precipitation in ethyl ether. The product was obtained as a flaky white solid (3.57 g, 91.5%). $^{1}$H NMR (400 MHz, DMSO-d$_6$) δ 7.74 (t, J = 5.7 Hz, 2H), 3.00 (q, J = 6.6 Hz, 4H), 2.59 (t, J = 7.1 Hz, 4H), 2.02 (d, J = 6.4 Hz, 4H), 1.71 – 1.32 (m, 12H), 1.25 (tq, J = 12.7, 7.6 Hz, 9H). $^{13}$C NMR (101 MHz, DMSO-d$_6$) δ 171.7, 40.5, 38.3, 35.3, 30.9, 29.1, 26.2, 25.9, 25.1.

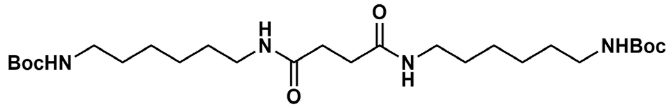


Di-*t*-butyl ((succinylbis(azanediyl))bis(hexane-6,1-diyl))dicarbamate (Boc-MSM). In a 500 mL RBF, EDC·HCl (9.74 g, 50.8 mmol, 2 equiv.) was added to a solution of succinic acid (3 g, 25.4 mmol, 1 equiv) dissolved in DCM (250 mL). *N*-Boc-1,6-diaminohexane (10.99 g, 50.8 mmol, 2 equiv.) was added the RBF, and the reaction was stirred at r.t. overnight. A white precipitate was formed over the course of the reaction which was filtered, rinsed with additional DCM and DI $H_2O$, and dried under vacuum to provide the product (7.71 g, 58.9%). $^{1}$H NMR (400 MHz, DMSO-d$_6$) δ 7.70 (t, J = 5.6 Hz, 1H), 6.75 (t, J = 5.7 Hz, 1H), 2.99 (q, J = 6.6 Hz, 2H), 2.88 (q, J = 6.6 Hz, 2H), 2.01 (d, J = 6.3 Hz, 2H), 1.60 – 1.41 (m, 2H), 1.36 (s, 12H), 1.21 (p, J = 3.7 Hz, 5H). $^{13}$C NMR (101 MHz, DMSO-d$_6$) δ 171.7, 155.6, 77.3, 38.3, 35.3, 29.4, 29.1, 28.3, 26.1, 26.0, 25.1.

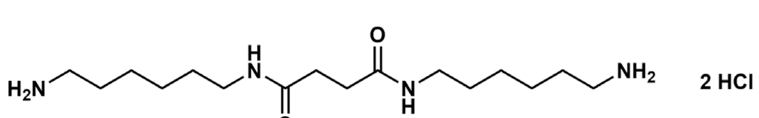


6,6'-(succinylbis(azanediyl))bis(hexan-1-aminium) hydrochloride (MSM HCl). In a 1000 mL RBF, Boc-MSM (7.7 g, 15.0 mmol, 1 equiv) is dissolved in 800 mL MeOH. The flask is cooled to 0 °C in an ice bath and subsequently charged with oxalyl chloride (3.85 mL, 45.0 mmol, 3 equiv.). The RBF was warmed to r.t. and stirred overnight. The following day, the RBF was concentrated via rotary evaporation and the residue purified by triturating with EtOH to provide the product as a white solid (4.93 g, 85.1%). $^{1}$H NMR (400 MHz, DMSO-d$_6$) δ 8.23 (s, 5H), 8.09 (t, J = 5.7 Hz, 3H), 3.23 (q, J = 6.5 Hz, 4H), 2.96 (h, J = 5.9 Hz, 4H), 2.72 (s, 5H), 1.76 (p, J = 7.1

Hz, 4H), 1.59 (p, J = 7.1 Hz, 4H), 1.54 – 1.21 (m, 8H). $^{13}$C NMR (101 MHz, DMSO-$d_6$) δ 171.7, 39.1, 38.8, 31.4, 29.4, 27.3, 26.3, 26.0.

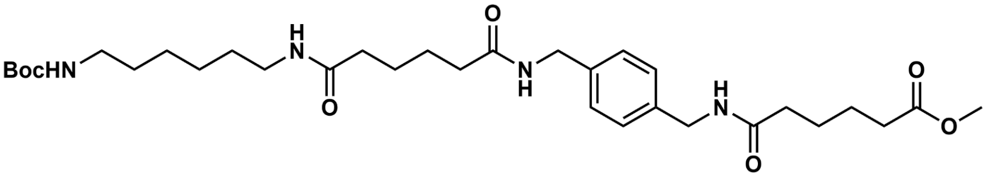

*Methyl 6-((4-(19,19-dimethyl-3,8,17-trioxo-18-oxa-2,9,16-triazaicosyl)benzyl)amino)-6-oxohexanoate (Boc-MAXA-OME)*. In a 1000 ml RBF, dissolved Boc-MA (10 g, 29 mmol, 1 equiv.) in DMF (200 mL). Charged RBF with EDC·HCl (6.12 g, 31.9 mmol, 1.1 equiv.). In a separate beaker, dissolved XA-OMe (11.39 g, 29 mmol, 1 equiv.) in DMF (200 mL) and neutralized trifluoroacetate with $Et_3N$ (6.07 mL, 43.5 mmol, 1.5 equiv.). Poured XA-OMe solution into RBF and stirred reaction at r.t. overnight, resulting in the formation of a solid white precipitate over the course of the reaction. The precipitate was filtered, rinsed with $H_2O$, and dried under vacuum to provide the product as a white powder (8.82 g, 50.2%). $^1$H NMR (400 MHz, DMSO-$d_6$) δ 8.26 (q, $J$ = 6.5 Hz, 2H), 7.71 (t, $J$ = 5.6 Hz, 1H), 7.17 (s, 4H), 6.75 (t, $J$ = 5.3 Hz, 1H), 4.21 (d, $J$ = 5.9 Hz, 4H), 3.58 (s, 3H), 3.00 (q, $J$ = 6.6 Hz, 2H), 2.88 (q, $J$ = 6.6 Hz, 2H), 2.31 (d, $J$ = 6.9 Hz, 2H), 2.21 – 1.94 (m, 6H), 1.62 – 1.42 (m, 8H), 1.36 (s, 13H), 1.29 – 1.04 (m, 4H). $^{13}$C NMR (101 MHz, DMSO-$d_6$) δ 173.7, 172.4, 172.2, 172.2, 156.0, 138.6, 127.6, 77.8, 51.7, 42.2, 38.8, 35.7, 35.7, 35.4, 33.5, 29.9, 29.6, 28.7, 26.6, 26.5, 25.6, 25.5, 25.2, 24.6.

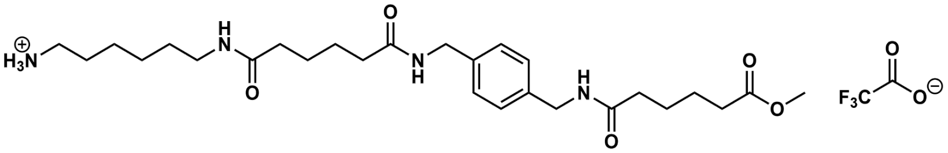

*6-(6-((4-((6-Methoxy-6-oxohexanamido)methyl)benzyl)amino)-6-oxohexanamido)hexan-1-aminium trifluoroacetate(MAXA-OMe)*. In a 500 mL RBF, added trifluoroacetic acid (70 mL) to a solution of MAXA-OMe in DCM (210 mL) and stirred reaction at r.t. overnight. The majority of trifluoroacetic acid was removed by concentrating the reaction *in vacuo* and diluting with DCM for 3 cycles. After the third cycle, the concentrated crude was precipitated into diethyl ether to form a white solid. The solid was filtered and dried under vacuum to provide MAXA-OMe in quantitative yield. $^1$H NMR (400 MHz, DMSO-$d_6$) δ 8.28 (q, J = 6.3 Hz, 4H), 7.88 – 7.46 (m, 8H), 7.17 (s, 8H), 4.21 (d, J = 5.8 Hz, 8H), 3.58 (s, 5H), 3.02 (q, J = 6.5 Hz, 4H), 2.76 (q, J = 6.8 Hz, 4H), 2.31 (d, J = 7.0 Hz, 3H), 2.18 – 2.07 (m, 8H), 2.05 (d, J = 7.3 Hz, 4H), 1.68 – 1.42 (m, 19H), 1.37 (p, J = 7.1 Hz, 5H), 1.33 – 1.13 (m, 8H). $^{13}$C NMR (101 MHz, DMSO-$d_6$) δ 173.3, 171.9, 171.8, 138.1, 127.1, 51.2, 41.7, 38.8, 38.2, 35.3, 35.2, 34.9, 33.0, 29.0, 26.9, 25.9, 25.4, 25.1, 25.0, 24.7, 24.1.

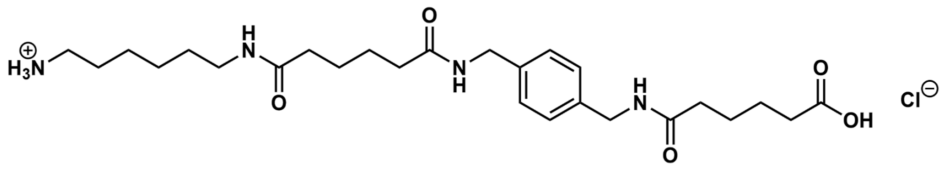

*6-((4-((6-((6-Aminohexyl)amino)-6-oxohexanamido)methyl)benzyl)amino)-6-oxohexanoic acid hydrochloride (MAXA HCl)*. In a 500 mL RBF, partially dissolved MAXA-OMe (8 g, 12.9 mmol) in MeOH (80 mL). Added 1 M LiOH (40 mL) to reaction, equipped reactor with reflux condenser, and heated to 60 °C overnight. Acidified reaction with 1 M HCl to pH 4 and removed MeOH *in vacuo*. Filtered resulting white solid and dried under vacuum to provide the product as a low melting white solid. $^{1}$H NMR (400 MHz, DMSO) δ 8.30 (q, $J$ = 5.1 Hz, 2H), 7.78 (t, $J$ = 5.6 Hz, 1H), 7.17 (s, 4H), 4.22 (d, $J$ = 5.9 Hz, 4H), 3.02 (q, $J$ = 6.6 Hz, 2H), 2.75 (t, $J$ = 7.5 Hz, 2H), 2.21 (t, $J$ = 6.8 Hz, 2H), 2.13 (m, 4H), 2.04 (d, $J$ = 6.0 Hz, 2H), 1.61 – 1.43 (m, 10H), 1.38 (t, $J$ = 6.8 Hz, 2H), 1.33 – 1.14 (m, 4H). $^{13}$C NMR (101 MHz, DMSO-$d_6$) δ 174.4, 172.0, 171.9, 171.8, 138.1, 127.2, 127.1, 41.7, 38.7, 38.2, 35.2, 35.2, 35.0, 33.4, 29.0, 26.9, 25.9, 25.5, 25.1, 25.0, 24.9, 24.2.

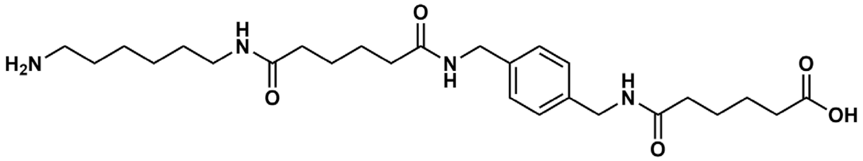

*6-((4-((6-((6-Aminohexyl)amino)-6-oxohexanamido)methyl)benzyl)amino)-6-oxohexanoic acid (MAXA)*. In a 500 mL RBF, suspended MAXA HCl (7 g, 13.3 mmol, 1 equiv.) in $H_2O$ (140 mL) and heated to 95 °C to dissolve. Added Reillex 402 (7 g, 5 equiv.) to solution, equipped RBF with reflux condenser, and stirred at 95 °C overnight. Filtered solution while hot. Removed $H_2O$ *in vacuo*, and suspended the resulting white solid residue in MeOH. The suspension was filtered to provide the product as a white solid (4.40 g, 67.5%). NMR analysis of MAXA was found to be spectroscopically identical to MAXA HCl.

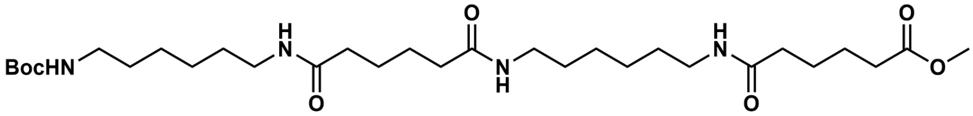

*Methyl 2,2-dimethyl-4,13,18,27-tetraoxo-3-oxa-5,12,19,26-tetraazadotriacontan-32-oate (Boc-MAMA-OMe)*. In a 250 mL RBF, dissolved Boc-MA (2.50 g, 11.6 mmol, 1 equiv.) in DMF (50 mL). Charged RBF with EDC·HCl (2.14 g, 12.77 mmol, 1.1 equiv.). In a separate beaker, dissolved MA-OMe (2.70 g, 11.61 mmol, 1 equiv.) in DMF (50 mL) and neutralized trifluoroacetate salt with $Et_3N$ (1.52 mL, 17.42 mmol, 1.5 equiv.). Poured MA-OMe solution into RBF and stirred reaction at r.t. overnight. The flask was chilled at -20 °C for 5 days, causing a white precipitate to form. The precipitate was filtered, rinsed with $H_2O$, and dried under vacuum to provide the product as a white powder (0.68 g, 16%). $^{1}$H NMR (400 MHz, DMSO-$d_6$) δ 7.72 (dt, J = 11.3, 5.6 Hz, 3H), 6.75 (s, 1H), 3.57 (s, 3H), 3.00 (q, J = 6.5 Hz, 6H), 2.30 (d, J = 6.6 Hz,

2H), 2.02 (t, J = 6.3 Hz, 6H), 1.46 (dq, J = 18.0, 3.7 Hz, 8H), 1.35 (d, J = 10.3 Hz, 17H), 1.22 (dt, J = 8.0, 3.8 Hz, 9H). $^{13}$C NMR (101 MHz, DMSO-$d_6$) δ 173.2, 171.7, 171.6, 155.6, 77.3, 51.2, 38.3, 35.3, 35.0, 33.0, 29.4, 29.1, 28.3, 26.1, 26.0, 25.1, 24.8, 24.0.

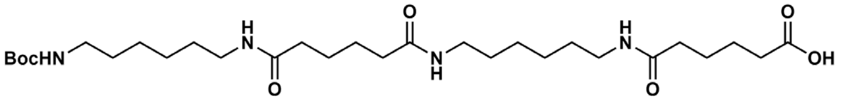


2,2-dimethyl-4,13,18,27-tetraoxo-3-oxa-5,12,19,26-tetraazadotriacontan-32-oic acid (Boc-MAMA). In a 50 mL RBF, Boc-MAMA-OMe (0.5 g, 0.85 mmol) was partially dissolved in MeOH (5 mL). The RBF was charged with 1 M LiOH (5 mL) and stirred for 3.5 h at r.t. The reaction was subsequently acified to pH ~4 with 1 M HCl. MeOH was removed via rotary evaporation, and reaction was subsequently filtered to provide the product as crystalline white powder (0.45 g, 92%).

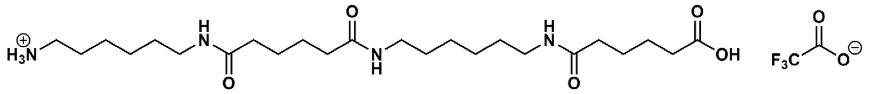


6-(6-((6-(5-carboxypentanamido)hexyl)amino)-6-oxohexanamido)hexan-1-aminium trifluoroacetate (MAMA trifluoroacetate). In a 100 mL RBF, Boc-MAMA (0.4 g, 0.7 mmol) was suspended in 10 mL DCM. The RBF was charged with trifluoroacetic acid (5 mL) and stirred at r.t. for 2 h. The majority of trifluoroacetic acid was removed by concentrating the reaction in vacuo and diluting with DCM for 3 cycles. After the third cycle, the concentrated crude was precipitated into diethyl ether causing a white solid to form. The solid was filtered and dried under vacuum to provide the product (0.24 g, 58%). $^{1}$H NMR (400 MHz, DMSO) δ 11.96 (b, 1 H), 8.31 – 7.03 (m, 5H), 3.01 (p, $J$ = 6.1 Hz, 5H), 2.77 (q, $J$ = 6.7 Hz, 2H), 2.27 – 2.10 (m, 2H), 2.03 (h, $J$ = 3.6 Hz, 5H), 1.66 – 1.41 (m, 9H), 1.36 (q, $J$ = 6.7 Hz, 6H), 1.30 – 0.98 (m, 7H). $^{13}$C NMR (101 MHz, DMSO) δ 174.4, 171.8, 171.8, 171.7, 158.2, 157.9, 38.8, 38.3, 38.2, 35.3, 35.1, 33.4, 29.1, 29.0, 26.9, 26.1, 25.9, 25.4, 25.1, 24.9, 24.1.

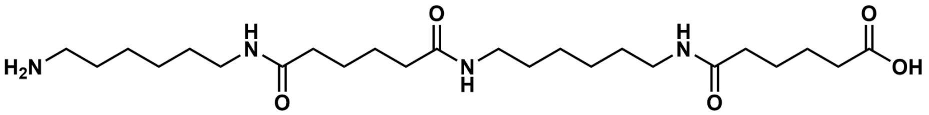


*6-((6-(6-((6-aminohexyl)amino)-6-oxohexanamido)hexyl)amino)-6-oxohexanoic acid (MAMA).* A 20 mL scintillation vial was charged with MAMA trifluoroacetate (200 mg, 0.34 mmol) and DI $H_2O$ (5 mL). Once the substrate had fully dissolved, Reillex 402 (0.18 g, 5 equiv.) was added and vial was capped and stirred at r.t. overnight. The reaction was subsequently filtered, and the solution was concentrated via rotary evaporation. The residue was precipitated in EtOH and filtered to provide the product as a white solid (12 mg, 7.5%).

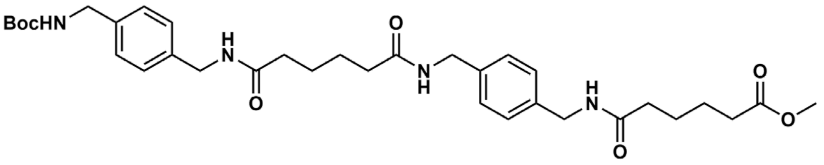

*Methyl 6-((4-((6-((4-(((t-butoxycarbonyl)amino)methyl)benzyl)amino)-6-oxohexanamido) methyl) benzyl)amino)-6-oxohexanoate (Boc-XAXA-OMe).* Boc-XA (4 g, 11.0 mmol, 1 equiv.) was dissolved in DMF (90 mL) in a 250 mL RBF. The RBF was charged with EDC·HCl (2.32 g, 12.1 mmol, 1.1 equiv.). In a separate beaker, XA-OMe (4.31 g, 11.0 mmol, 1 equiv.) was dissolved in DMF (90 mL) and neutralized trifluoroacetate salt with $Et_3N$ (2.29 mL, 16.65 mmol, 1.5 equiv.). Poured XA-OMe solution into RBF and stirred reaction at r.t. overnight, resulting in the formation of a solid white precipitate over the course of the reaction. The precipitate was filtered, rinsed with $H_2O$, and dried under vacuum to provide the product as a white powder (2.49 g, 36%). $^1$H NMR (400 MHz, DMSO-$d_6$) δ 8.26 (q, J = 6.1 Hz, 2H), 7.17 (s, 6H), 4.21 (d, J = 5.9 Hz, 4H), 4.08 (d, J = 6.2 Hz, 2H), 3.62 – 3.53 (m, 3H), 2.31 (d, J = 6.7 Hz, 2H), 2.11 (d, J = 7.3 Hz, 5H), 1.63 – 1.44 (m, 6H), 1.38 (s, 7H). $^{13}$C NMR (101 MHz, DMSO-$d_6$) δ 173.3, 171.9, 171.8, 138.6, 138.2, 138.0 127.4, 127.1, 127.1, 126.9, 77.7, 51.2, 44.5, 43.1, 41.7, 35.2, 35.1, 34.9, 33.0, 28.2, 25.1, 24.9, 24.7, 24.2, 24.1.

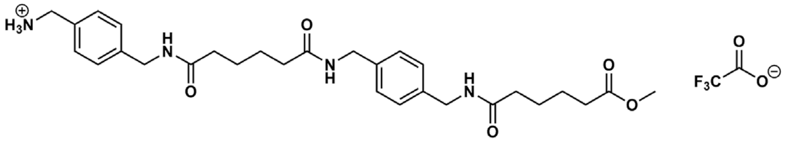

*(4-((6-((4-((6-Methoxy-6-oxohexanamido)methyl)benzyl)amino)-6-oxohexanamido)methyl) phenyl)methanaminium trifluoroacetate (XAXA-OMe).* In a 250 mL rbf, suspended Boc-XAXA-OMe (2.00 g, 3.20 mmol) in DCM (30 mL). Added trifluoroacetic acid (10 mL) and stirred at r.t. for 3 h. The majority of trifluoroacetic acid was removed by concentrating the reaction *in vacuo* and diluting with DCM for 3 cycles. After the third cycle, the concentrated crude was precipitated into diethyl ether causing a white solid to form. The solid was filtered and dried under vacuum to provide XAXA-OMe (1.77 g, 86.8%). $^1$H NMR (400 MHz, DMSO-$d_6$) δ 8.31 (dt, J = 27.2, 6.2 Hz, 3H), 8.16 (s, 3H), 7.33 (dd, J = 42.3, 7.7 Hz, 5H), 7.17 (s, 3H), 4.26 (d, J = 5.9 Hz, 2H), 4.21 (d, J = 5.8 Hz, 3H), 4.00 (q, J = 5.8 Hz, 3H), 3.59 (s, 3H), 2.30 (t, J = 6.4 Hz, 2H), 2.12 (t, J = 6.5 Hz, 6H), 1.51 (s, 8H). $^{13}$C NMR (101 MHz, DMSO-$d_6$) δ 173.3, 172.0, 171.9, 171.8, 140.3, 138.1, 132.3, 128.8, 127.3, 127.1, 51.2, 42.0, 41.7, 41.6, 35.2, 34.9, 33.0, 25.0, 24.7, 24.1.

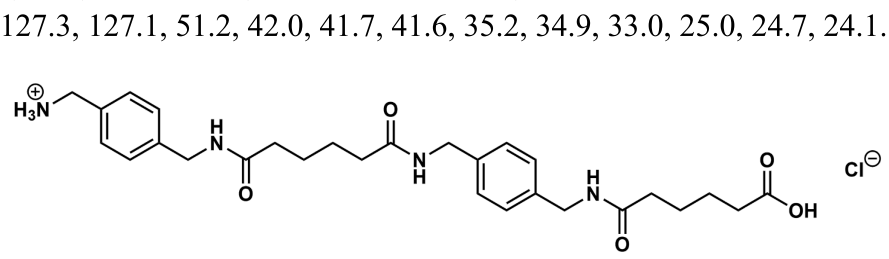

(4-((6-((4-((5-carboxypentanamido)methyl)benzyl)amino)-6-oxohexanamido)methyl) phenyl)methanaminium (XAXA hydrochloride). In a 100 mL RBF, Added 1 M LiOH (15 mL) to a solution of XAXA-OMe (1.5 g, 2.35 mmol) in MeOH (15 mL). The RBF was equipped with a reflux condenser, heated to 50 °C, and stirred overnight. The reaction was subsequently acidified to pH ~4 with 1 M HCl. Methanol was removed via rotary evaporation, and the reaction was filtered to provide the product as a white solid (0.79 g, 62%).

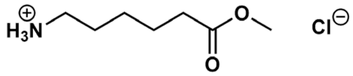

*Methyl 6-aminohexanoate hydrochloride.* A 250 mL RBF was charged with 6-aminohexanoic acid (5 g, 38.1 mmol, 1 equiv.) and MeOH (125 mL). The RBF was cooled to 0 °C in an ice bath and equipped with a dropwise addition funnel containing thionyl chloride (4.8 mL, 66.7 mmol, 1.75 eq uiv.). Thionyl chloride was added to the RBF dropwise over the course of 1 h, and subsequently stirred at r.t. overnight. The RBF was concentrated by rotary evaporation and subsequently dried in a vacuum overnight to provide the product as a white solid (4.82 g, 70%). Characterization data was found to match reported literature.[2]

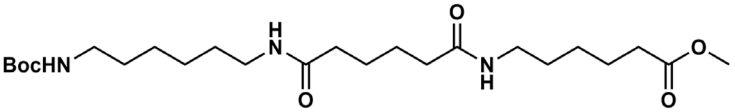

*Methyl 6-(6-((6-N-Boc-aminohexyl)amino)-6-oxohexanamido)hexanoate (Boc-MAH-OMe).* In a 20 mL scintillation vial, boc-MA (0.5 g, 1.3 mmol, 1 equiv) was dissolved in DMF (3.5 mL) with EDC·HCl (0.31 g, 1.4 mmol, 1.1 equiv). In a separate vial, methyl 6-aminohexanoate hydrochloride (0.29 g, 1.6 mmol, 1.1 equiv) was dissolved in DMF (3.5 mL) and $Et_3N$ (0.23 g, 1.5 mmol, 1.1 equiv). The two solutions were combined and stirred at r.t. overnight. The solution was dissolved in 50 mL DCM and extracted with 3×40 mL saturated aqueous $NH_4Cl$ solution, 3×40 mL 1 M HCl solution, 3×40 mL saturated $NaHCO_3$ solution, and 40 mL brine. The organic layer was dried over $MgSO_4$, filtered, and dried using a rotary evaporator to provide the product as a white solid (0.27 g, 40%). $^1$H NMR (400 MHz, $CDCl_3$) δ 6.37 – 5.58 (m, 2H), 4.58 (s, 1H), 3.66 (s, 3H), 3.24 (p, *J* = 6.3 Hz, 5H), 3.10 (d, *J* = 7.7 Hz, 3H), 2.31 (t, *J* = 7.4 Hz, 3H), 2.20 (q, *J* = 4.9 Hz, 5H), 1.64 (q, *J* = 5.8 Hz, 7H), 1.57 – 1.39 (m, 17H), 1.34 (qd, *J* = 8.2, 4.6 Hz, 8H). $^{13}$C NMR (101 MHz, $CDCl_3$) δ 174.2, 173.1, 79.2, 51.7, 39.4, 36.2, 34.0, 30.1, 29.5, 29.3, 28.6, 26.5, 25.1, 24.6.

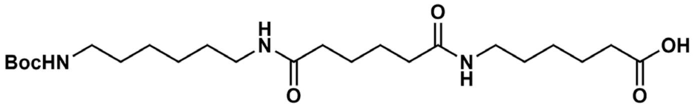

*6-(6-((6-N-Boc-aminohexyl)amino)-6-oxohexanamido)hexanoic acid (Boc-MAH).* In a 20 mL scintillation vial, partially dissolved Boc-MAH-OMe (0.2 g, 0.42 mmol) in THF (2 mL) and added an equal volume of 1 M LiOH. The mixture was stirred at r.t. for 3.5 h resulting in the full dissolution of the starting material. The reaction was subsequently acidified to pH ~4 by addition of 1M HCl. THF was removed by rotary evaporation. The resulting white suspension was centrifuged, and the supernatant was discarded. The precipitate was dried under vacuum to provide the product as a white powder (0.15 g, 78%). $^{1}$H NMR (400 MHz, DMSO-$d_6$) δ 11.97 (s, 1H), 7.71 (q, J = 5.2 Hz, 2H), 6.75 (s, 1H), 2.99 (q, J = 6.6 Hz, 4H), 2.88 (q, J = 6.6 Hz, 2H), 2.18 (t, J = 7.3 Hz, 2H), 2.01 (d, J = 6.3 Hz, 4H), 1.64 – 1.40 (m, 6H), 1.36 (s, 15H), 1.29 – 1.03 (m, 7H). $^{13}$C NMR (101 MHz, DMSO-$d_6$) δ 174.9, 172.2, 156.0, 77.8, 38.8, 38.7, 35.8, 34.1, 29.9, 29.6, 29.4, 28.7, 26.6, 26.4, 25.6, 24.7.

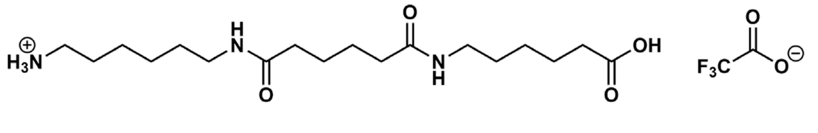

*6-(6-((5-carboxypentyl)amino)-6-oxohexanamido)hexan-1-aminium trifluoroacetate (MAH trifluoroacetate).* In a 20 mL scintillation vial, Boc-MAH (0.15 g, 0.33 mmol) was suspended in DCM (2 mL) and the vial was charged with trifluoroacetic acid (1 mL). The reaction was stirred at r.t. for 2 h. The majority of trifluoroacetic acid was removed by concentrating the reaction *in vacuo* and diluting with DCM for 3 cycles. After the third cycle, the concentrated crude was precipitated into diethyl ether. The ether was decanted and the residue was dried under vacuum to provide the product as an oil which slowly crystallized over the course of ~48 h. $^{1}$H NMR (400 MHz, DMSO-$d_6$) δ 11.99 (s, 1H), 8.46 – 6.77 (m, 5H), 3.00 (p, $J$ = 6.4 Hz, 4H), 2.76 (q, $J$ = 7.7 Hz, 2H), 2.18 (t, $J$ = 7.4 Hz, 2H), 2.03 (s, 4H), 1.48 (dd, $J$ = 26.8, 9.8 Hz, 8H), 1.37 (t, $J$ = 7.2 Hz, 4H), 1.26 (dq, $J$ = 15.4, 8.2 Hz, 6H). $^{13}$C NMR (101 MHz, DMSO-$d_6$) δ 174.4, 171.8, 171.8, 38.8, 38.2, 38.2, 35.3, 33.6, 29.0, 28.9, 26.9, 26.0, 25.9, 25.4, 25.1, 24.2.

6-(6-((6-aminohexyl)amino)-6-oxohexanamido)hexanoic acid (MAH). Dissolved MAH trifluoroacetate (50 mg, 0.11 mmol, 1 equiv) in DI water (5 mL) in a 20 mL scintillation vial and added Reillex 402 (56 mg, 5 equiv.). Stirred at r.t. overnight. The reaction was filtered, and the filtrate was concentrated via rotary evaporation. The residue was precipitated in EtOH to provide the product as a white solid (18.3 mg, 48.3%). $^{1}$H NMR (400 MHz, $D_2O$) δ 3.18 (t, J = 6.8 Hz, 4H), 2.99 (t, J = 7.6 Hz, 2H), 2.50 – 2.00 (m, 6H), 1.82 – 1.45 (m, 12H), 1.45 – 0.96 (m, 6H). $^{13}$C NMR (101 MHz, $D_2O$) δ 181.2, 176.4, 176.3, 39.4, 39.1, 39.0, 35.5, 35.4, 28.0, 28.0, 26.6, 25.7, 25.4, 25.1, 24.9, 24.6.

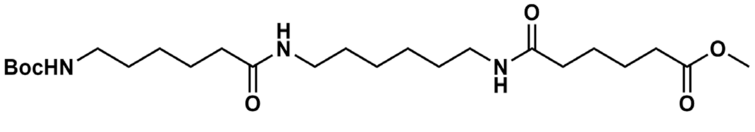

*Methyl 6-((6-(6-N-boc-aminohexanamido)hexyl)amino)-6-oxohexanoate (Boc-HMA-OMe).* Boc-HMA-OMe was synthesized following the procedure described for Boc-MAH-OMe using MA-OMe (0.5 g, 1.3 mmol, 1 equiv) and *N*-boc-aminocaproic acid (0.34 g, 1.4 mmol, 1.1 equiv) as the starting materials. Product was collected as a white solid (0.21 g, 33%). $^1$H NMR (400 MHz, $CDCl_3$) δ 5.84 (s, 1H), 5.74 (s, 1H), 4.60 (s, 1H), 3.66 (s, 3H), 3.23 (q, $J$ = 6.5 Hz, 4H), 3.09 (s, 2H), 2.34 (d, $J$ = 6.7 Hz, 2H), 2.26 – 2.06 (m, 4H), 1.92 (s, 2H), 1.64 (h, $J$ = 6.6 Hz, 6H), 1.50 (q, $J$ = 7.1 Hz, 6H), 1.43 (s, 9H), 1.39 – 1.11 (m, 7H). $^{13}$C NMR (101 MHz, $CDCl_3$) δ 174.1, 173.1, 172.8, 156.2, 51.7, 39.1, 39.1, 36.7, 36.4, 33.8, 29.9, 29.6, 28.6, 26.5, 26.1, 25.5, 25.3, 24.6.

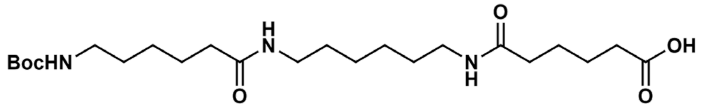


6-((6-(6-*N*-Boc-aminohexanamido)hexyl)amino)-6-oxohexanoic acid (Boc-HMA). Boc-HMA was prepared following the method described for Boc-MAH by dissolving boc-HMA-OMe (0.20 g, 0.42 mmol) in THF (2 mL) and 1 M LiOH (2 mL). Product was isolated as a white powder (0.16 g, 80%). $^1$H NMR (400 MHz, DMSO-$d_6$) δ 11.97 (s, 1H), 8.23 – 7.29 (m, 2H), 6.75 (s, 1H), 3.32 (s, 3H), 2.99 (q, $J$ = 6.5 Hz, 4H), 2.88 (q, $J$ = 6.9 Hz, 2H), 2.18 (t, $J$ = 7.3 Hz, 2H), 2.02 (s, 4H), 1.68 – 1.41 (m, 6H), 1.36 (s, 14H), 1.22 (d, $J$ = 7.4 Hz, 7H). $^{13}$C NMR (101 MHz, DMSO-$d_6$) δ 174.4, 171.7, 155.6, 77.3, 38.3, 38.2, 35.3, 33.6, 29.4, 29.1, 28.9, 28.3, 26.1, 26.0, 25.1, 24.2.

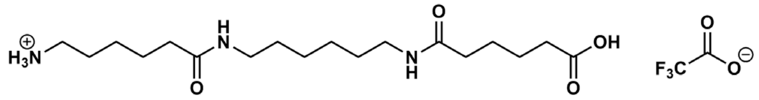


*6-((6-(5-Carboxypentanamido)hexyl)amino)-6-oxohexan-1-aminium trifluoroacetate (HMA trifluoroacetate).* HMA trifluoroacetate was synthesized following the procedure described for MAH trifluoroacetate by dissolving Boc-HMA (1.5 g, 0.33 mmol) in DCM (2 mL) and trifluoroacetic acid (1 mL). Product was isolated as an oil that slowly crystallized over the course of ~48 h. $^1$H NMR (400 MHz, DMSO-$d_6$) δ 11.98 (s, 1H), 8.66 – 6.88 (m, 5H), 3.00 (q, J = 6.5 Hz, 4H), 2.86 – 2.62 (m, 2H), 2.19 (t, J = 6.6 Hz, 2H), 2.04 (t, J = 7.4 Hz, 4H), 1.48 (dp, J = 11.1, 3.9 Hz, 8H), 1.35 (q, J = 6.8 Hz, 4H), 1.31 – 1.09 (m, 6H). $^{13}$C NMR (101 MHz, DMSO-$d_6$) δ 174.4, 171.7, 171.6, 157.8, 38.7, 38.3, 38.3, 35.1, 33.4, 29.1, 26.8, 26.1, 25.5, 24.9, 24.8, 24.1.

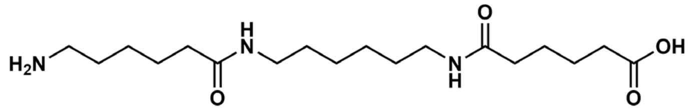


6-((6-(6-Aminohexanamido)hexyl)amino)-6-oxohexanoic acid (HMA). In a 20 mL scintillation vial, dissolved HMA trifluoroacetate (50 mg, 0.11 mmol, 1 equiv) in DI water (10 mL). Added Reillex 402 (56 mg, 5 equiv.). Stirred at r.t. overnight. The reaction was subsequently filtered and the filtrate was collected. Water was removed via rotary evaporation, the residue was precipitated in ethanol to provide the product as a white powder (11.7 mg, 30.9 %). $^1$H NMR (400 MHz, $D_2O$) δ 3.09 (t, $J$ = 6.8 Hz, 4H), 2.90 (t, $J$ = 7.6 Hz, 2H), 2.16 (t, $J$ = 7.0 Hz, 6H), 1.77 – 1.47 (m, 8H), 1.47 – 1.35 (m, 4H), 1.35 – 1.06 (m, 6H). $^{13}$C NMR (101 MHz, $D_2O$) δ 182.5, 176.6, 176.5, 39.2, 39.2, 39.1, 36.5, 35.6, 35.5, 28.1, 28.1, 26.4, 25.5, 25.3, 25.0, 24.9, 24.9.

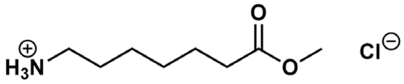

*Methyl 7-aminoheptanoate hydrochloride.* In a 250 mL RBF, $SOCl_2$ (7.50 mL, 103.3 mmol, 3 equiv.) was added dropwise to a stirred solution 7-aminoheptanoic acid (5 g, 34.4 mmol, 1 equiv.) in MeOH (50 mL) at 0 °C. The RBF was warmed to room temperature and stirred for an additional 3 h. The reaction mixture was subsequently concentrated via rotary evaporation and precipitated into tetrahydrofuran which, upon filtration, provided the product as a white solid (5.73 g, 91.7%). Characterization data was found to match reported literature.[3]

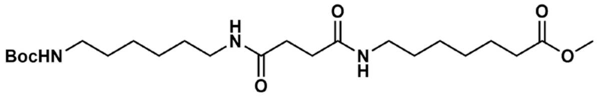

*Methyl 7-(4-((6-N-boc-aminohexyl)amino)-4-oxobutanamido)heptanoate (Boc-MSP-OMe).* In a 20 mL scintillation vial, dissolved Boc-MS (1.00 g, 3.2 mmol, 1 equiv.) in DMF (5 mL) and added EDC HCl (0.67 g, 3.5 mmol, 1.1 equiv.). In a separate vial dissolved methy heptanoate hydrochloride (0.68 g, 3.5 mmol, 1.1 equiv.) in DMF (3.5 mL) and neutralized with $Et_3N$ (0.7 mL, 4.7 mmol, 1.5 equiv.). Combined solutions and stirred at r.t. overnight. The solution was dissolved in 100 mL DCM and extracted with 3×100 mL saturated aqueous $NH_4Cl$ solution, 3×100 mL 1 M HCl solution, 3×100 mL saturated $NaHCO_3$ solution, and 100 mL brine. The organic layer was dried over $Na_2SO_4$, filtered, and dried using a rotary evaporator to provide the product as a white solid (1.07 g, 73.8%). $^1$H NMR (400 MHz, DMSO-$d_6$) δ 7.98 (t, $J$ = 5.6 Hz, 2H), 6.99 (t, $J$ = 5.8 Hz, 1H), 3.81 (s, 3H), 3.23 (q, $J$ = 6.5 Hz, 4H), 3.12 (q, $J$ = 6.6 Hz, 2H), 2.51 (d, $J$ = 8.5 Hz, 6H), 1.73 (q, $J$ = 7.3 Hz, 2H), 1.60 (s, 14H), 1.46 (dp, $J$ = 11.6, 3.8 Hz, 8H). $^{13}$C NMR (101 MHz, DMSO-$d_6$) δ 173.4, 171.1, 155.6, 77.3, 51.2, 38.4, 33.2, 31.0, 29.4, 29.1, 28.9, 28.3, 28.2, 26.1, 26.0, 24.4.

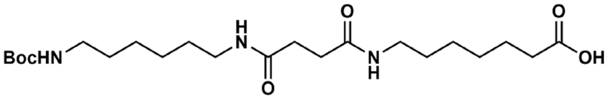

*7-(4-((6-N-boc-aminohexyl)amino)-4-oxobutanamido)heptanoic acid (Boc-MSP).* In a 100 mL RBF, 1 M LiOH (9 mL) was added to stirred solution of Boc-MSP-OMe (0.9 g, 1.97 mmol) in THF (9 mL). The reaction was stirred at r.t. for 4 h and subsequently acified to pH ~4 by addition of 1 M HCl. THF was removed via rotary evaporation and the reaction was filtered to provide the product as a white solid (0.69 g, 79%). $^1$H NMR (400 MHz, DMSO-$d_6$) δ 11.96 (s, 1H), 7.75 (t, $J$ = 5.7 Hz, 2H), 6.75 (t, $J$ = 5.8 Hz, 1H), 2.99 (q, $J$ = 6.6 Hz, 4H), 2.88 (q, $J$ = 6.7 Hz, 2H), 2.26 (s, 4H), 2.18 (t, $J$ = 7.4 Hz, 2H), 1.46 (q, $J$ = 7.2 Hz, 3H), 1.36 (s, 15H), 1.29 – 0.99 (m, 9H). $^{13}$C NMR (101 MHz, DMSO-$d_6$) δ 174.5, 171.1, 155.6, 77.3, 38.4, 33.6, 31.0, 29.4, 29.1, 29.0, 28.3, 26.1, 26.0, 24.4.

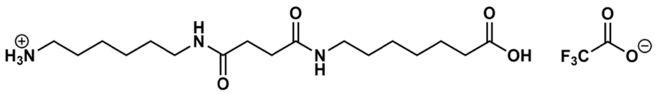

*6-(4-((6-carboxyhexyl)amino)-4-oxobutanamido)hexan-1-aminium trifluoroacetate (MSP trifluoroacetate).* MSP trifluoroacetate was synthesized following the procedure described for MAH trifluoroacetate by dissolving Boc-MSP (0.5 g, 1.12 mmol) in DCM (7.5 mL) and trifluoroacetic acid (2.5 mL). Product was isolated as a white solid (0.47 g, 90%). $^{1}$H NMR (400 MHz, $D_2O$) δ 3.16 (t, *J* = 6.8 Hz, 4H), 2.99 (t, *J* = 7.6 Hz, 2H), 2.50 (s, 4H), 2.39 (t, *J* = 7.4 Hz, 2H), 1.62 (dq, *J* = 21.8, 7.2 Hz, 4H), 1.54 – 1.43 (m, 4H), 1.43 – 1.12 (m, 9H). $^{13}$C NMR (101 MHz, $D_2O$) δ 179.2, 174.4, 174.4, 114.9, 39.4, 39.2, 39.1, 33.7, 31.5, 31.5, 28.1, 28.0, 27.8, 26.6, 25.6, 25.4, 25.2, 24.2.

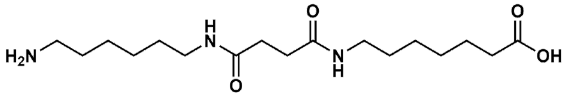

7-(4-((6-aminohexyl)amino)-4-oxobutanamido)heptanoic acid (MSP). A 20 mL scintillation vial was charged with MSP trifluoroacetate (0.4 g, 0.87 mmol) DI $H_2O$ (10 mL) and heated to 80 °C to dissolve. Reillex 402 (1 g, 11 equiv) was added to the vial, and the reaction was stirred at 80 °C overnight. The reaction was filtered, and the filtrate was concentrated via rotary evaporation and subsequently triturated with EtOH to provide the product as a white solid (0.083 g, 28%). $^{1}$H NMR (400 MHz, $D_2O$) δ 3.16 (d, *J* = 7.0 Hz, 4H), 2.99 (t, *J* = 7.6 Hz, 2H), 2.51 (s, 4H), 2.23 (t, *J* = 7.4 Hz, 2H), 1.66 (p, *J* = 7.3 Hz, 2H), 1.53 (dq, *J* = 20.3, 6.8 Hz, 6H), 1.44 – 1.17 (m, 8H). $^{13}$C NMR (101 MHz, $D_2O$) δ 182.7, 174.4, 174.3, 39.4, 39.3, 39.1, 36.5, 31.6, 31.6, 28.2, 28.1, 28.0, 26.6, 25.7, 25.4, 25.3, 25.2.

6-[[6-[[6-[(6-Aminohexyl)amino]-1,6-dioxohexyl]amino]hexyl]amino]-6-oxohexanoic acid (MAMA) was synthesized as previously described. [4]

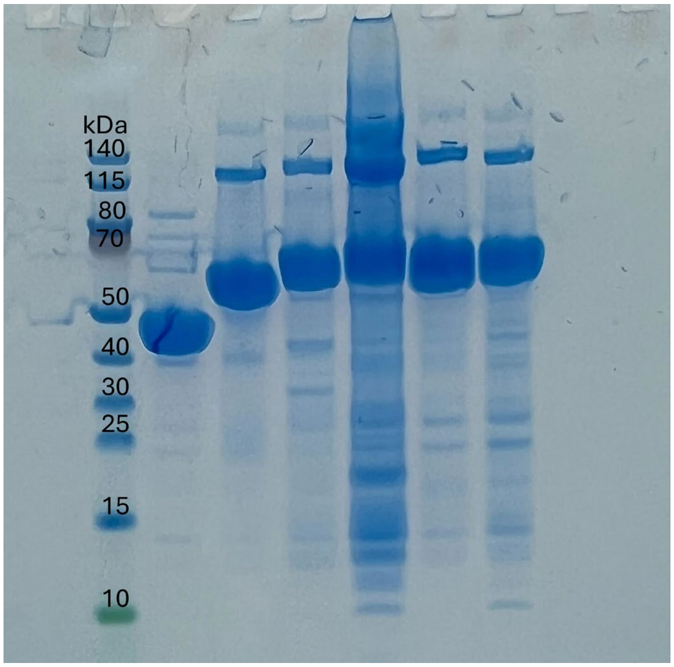


**Figure S1: SDS-PAGE analysis of recombinant N-terminal His-tagged proteins purified from *E.coli* BL21(DE3).** Novex Tris-Glycine gels (4%-12%, Invitrogen) were used. DdaG expected size: 45.58 kDa; SfaB expected size: 65.06 kDa; AsbA expected size: 69.91 kDa; AsbB expected size: 72.01 kDa; AcsA expected size: 70.7 kDa; DesD expected size: 66.6 kDa.

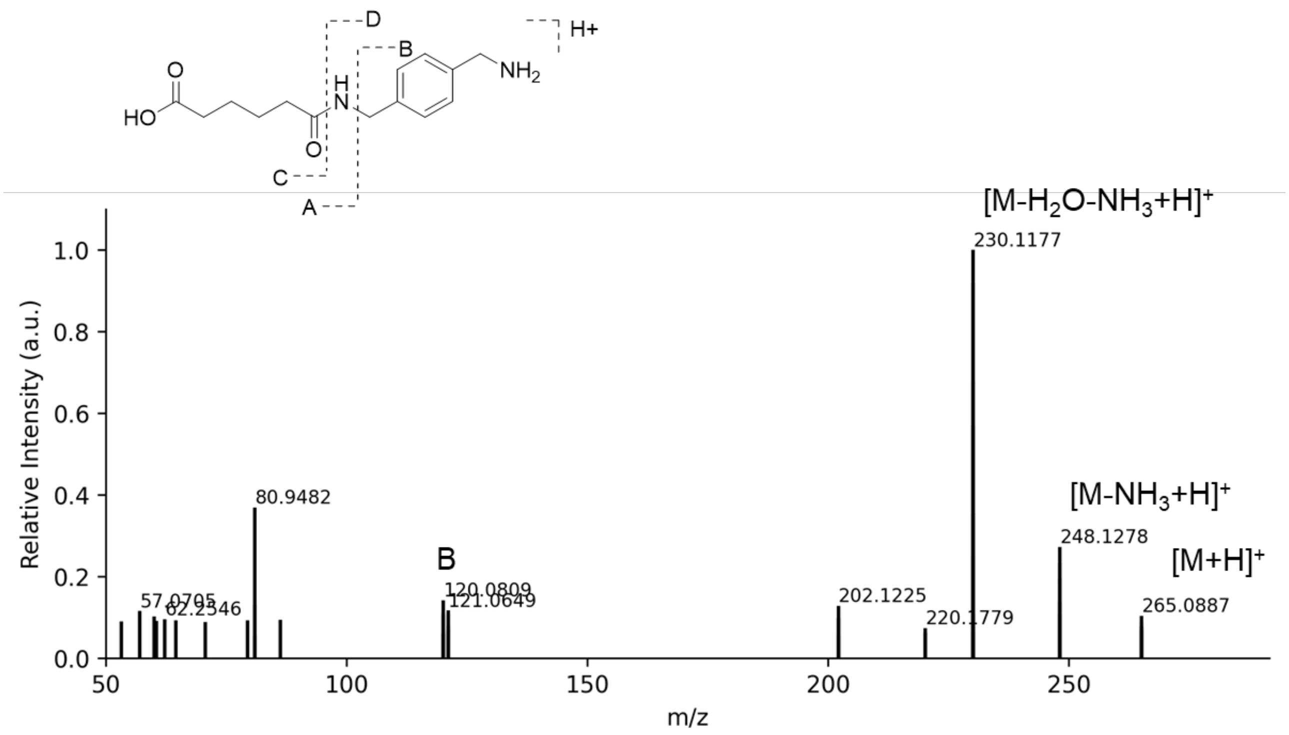


**Figure S2: I.DOT/OPSI-MS$^2$ of XA.** A collision energy of 35 eV was used.

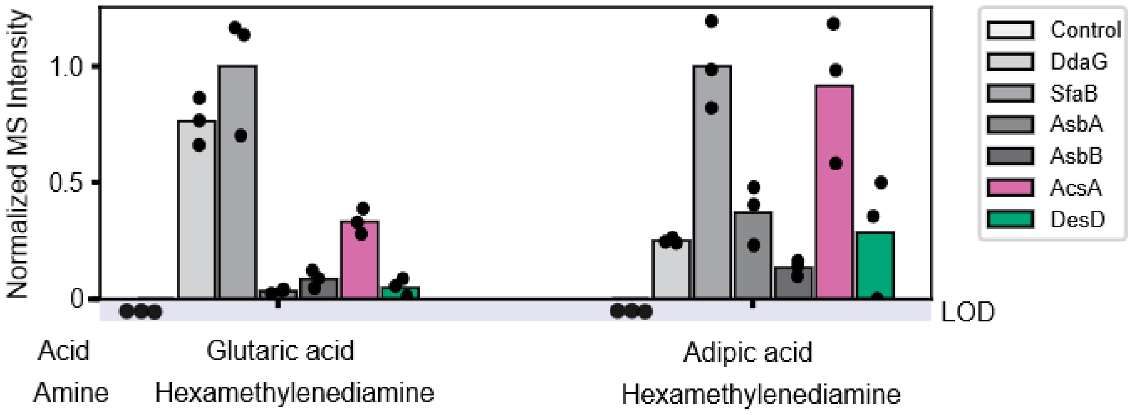


**Figure S3**. **Comparison of NIS synthetases and previously identified amide synthetases for synthesis of ω-amino acid diads**. *In vitro* biochemical assays were conducted by incubating different diacids (i.e., glutaric acid and adipic acid) with hexamethylenediamine in the presence of enzymes (i.e., DdaG, SfaB, AsbA, AsbB, AcsA and DesD) or no-enzyme control. The MS signal intensity was normalized to SfaB. Products were assayed using OPSI-MS. All replicates shown are biological replicates. Sample size is *N*=3. LOD: limit of detection.

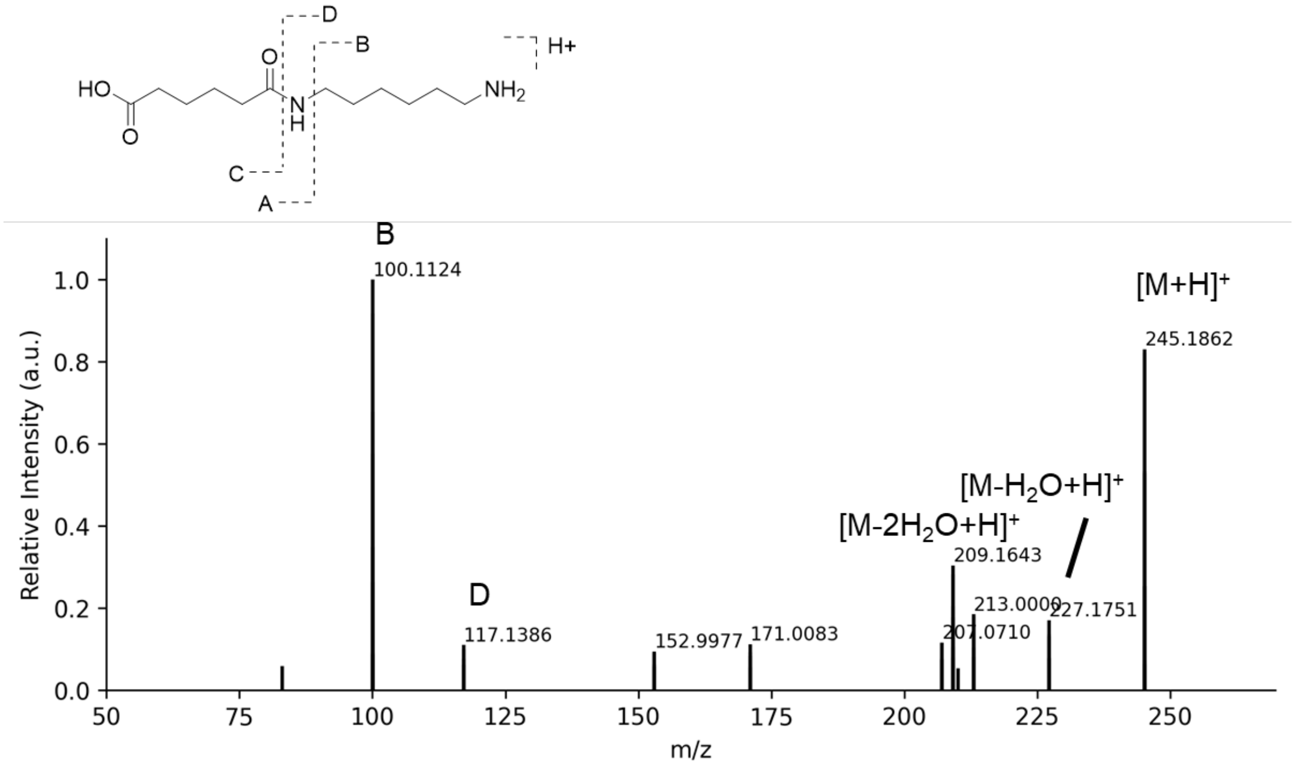


**Figure S4: I.DOT/OPSI-MS² of MA.** A collision energy of 35 eV was used.

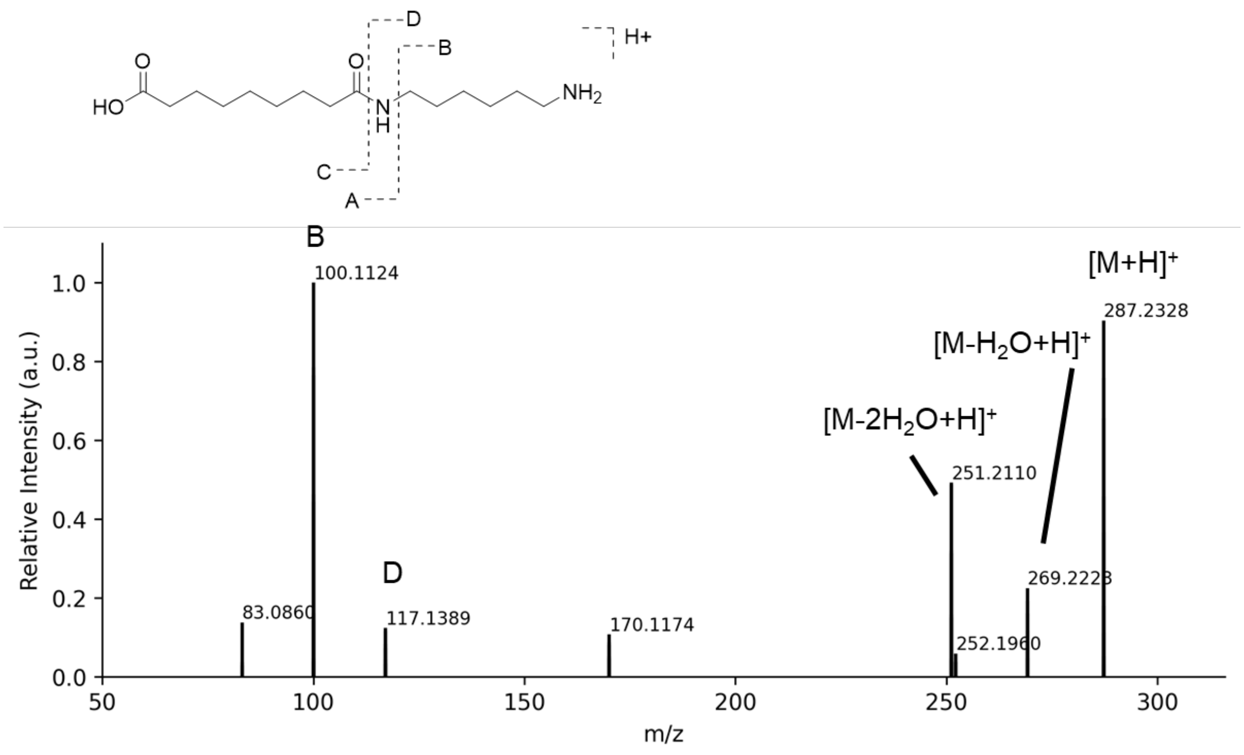


**Figure S5: I.DOT/OPSI-MS² of MZ.** A collision energy of 35 eV was used.

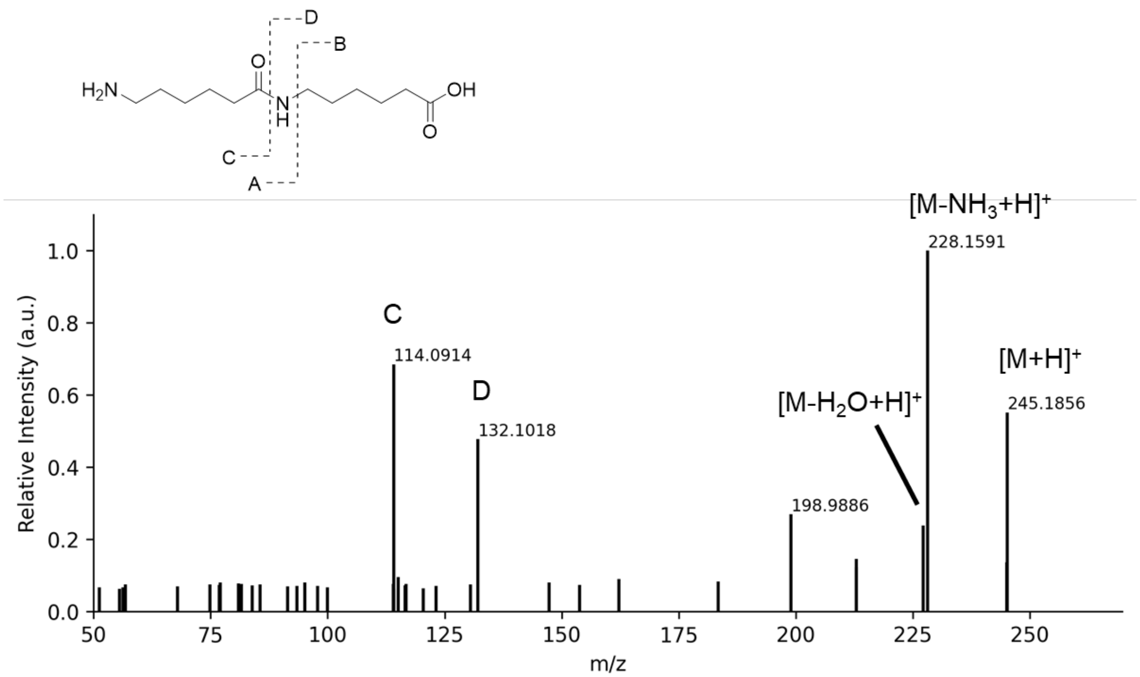


**Figure S6: I.DOT/OPSI-MS$^2$ of HH.** A collision energy of 35 eV was used.

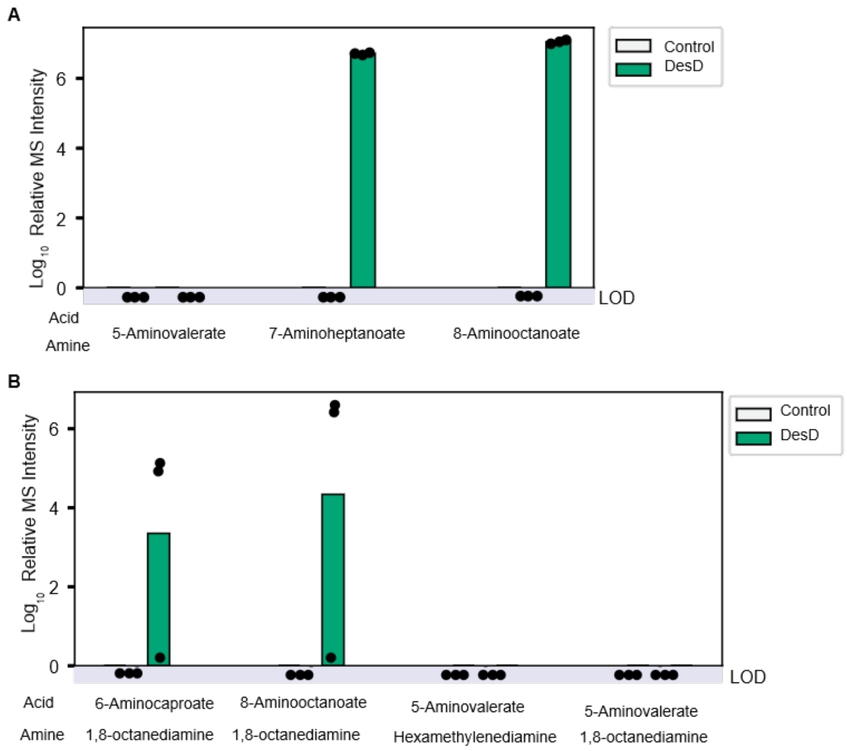


**Figure S7**. **DesD can synthesize diverse diads**. A) *In vitro* biochemical assays were conducted by incubating single ω-amino acids (i.e., 5-aminovalerate, 7-aminohepanoate, or 8-aminooctanoate) in the presence of DesD or no-enzyme control. Synthesis of the associated ω-amino acid diad was measured by OPSI-MS and is reported as the $\log_{10}$ signal intensity. B) *In vitro* biochemical assays were conducted by incubating different ω-amino acids (i.e., 5-aminovalerate, 6-aminocaporate, or 8-aminooctanoate) with diamines (i.e., hexamethylene diamine or 1,8-octanediamine) in the presence of DesD or no-enzyme control. Diamine diad products were assayed using OPSI-MS and is reported as the $\log_{10}$ signal intensity. All replicates shown are biological replicates. Sample size is $N$=3. LOD: limit of detection.

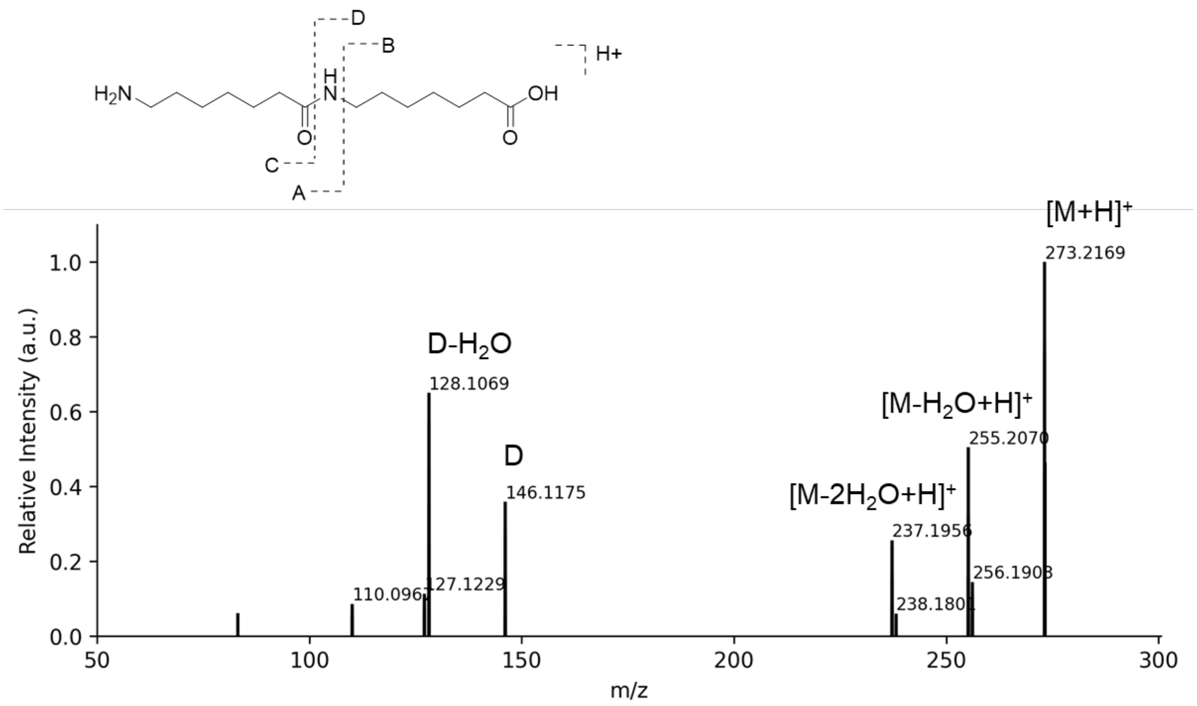


**Figure S8: I.DOT/OPSI-MS$^2$ of the dimerization product of 7-aminoheptanoic acid.** A collision energy of 35 eV was used.

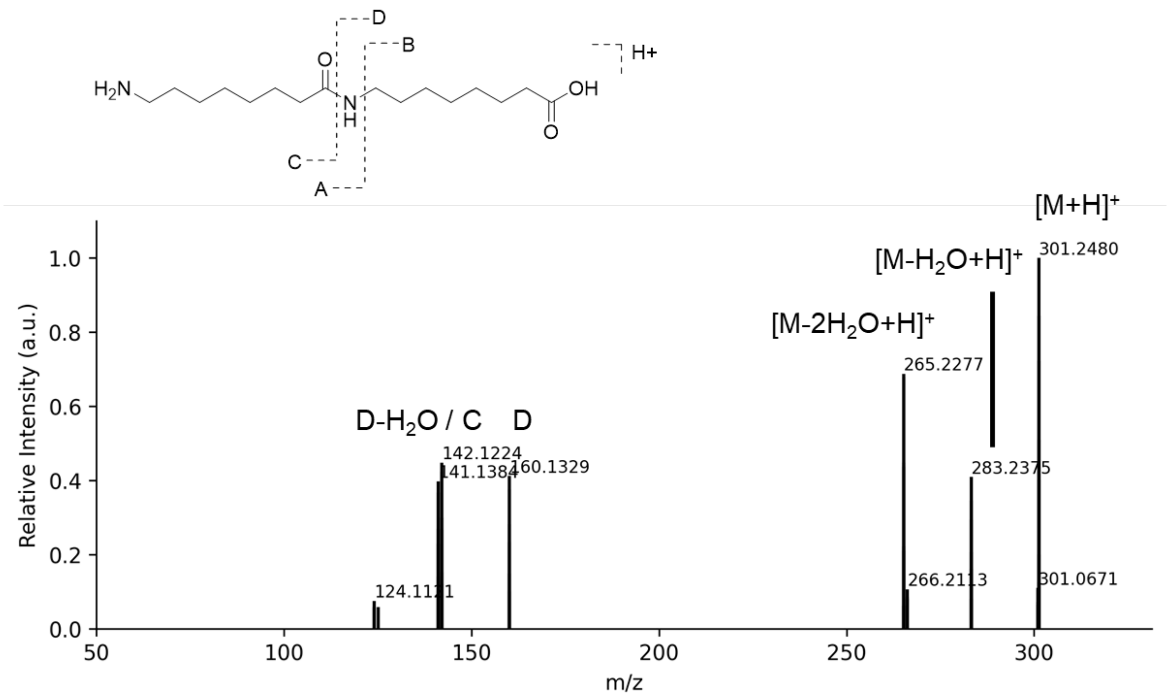


**Figure S9: I.DOT/OPSI-$MS^2$ of the dimerization product of 8-aminooctanoic acid.** A collision energy of 35 eV was used.

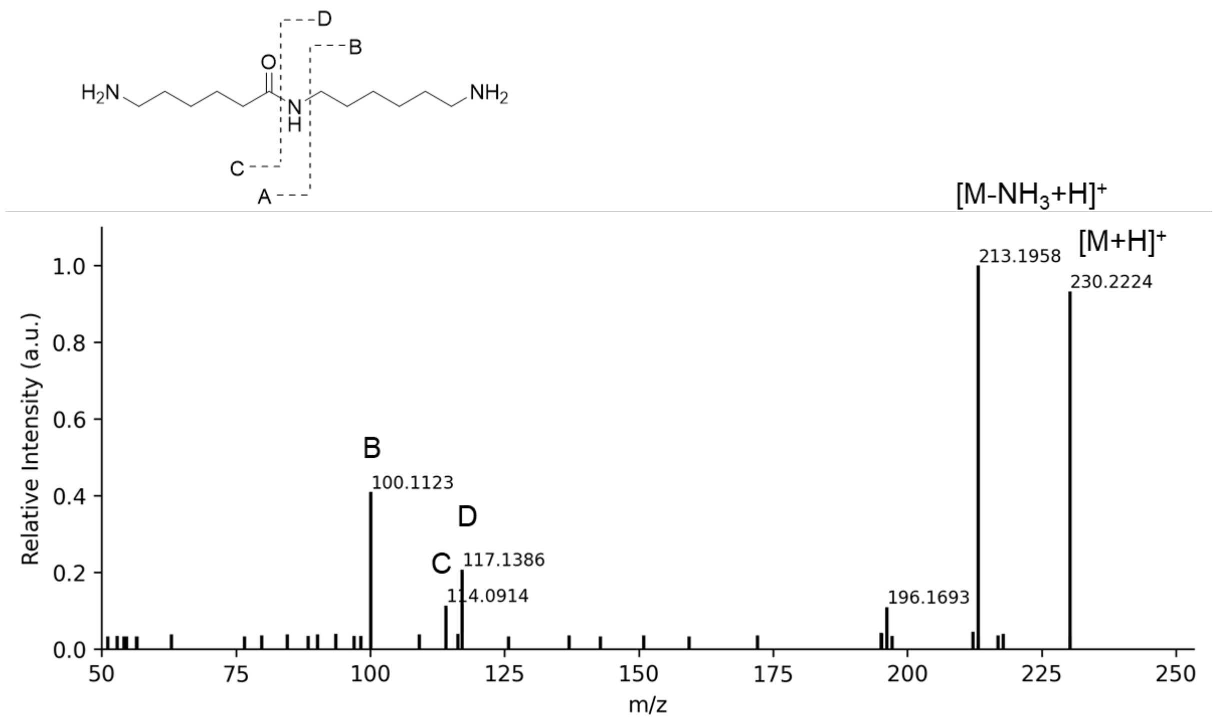


**Figure S10: I.DOT/OPSI-MS$^2$ of MH.** A collision energy of 35 eV was used.

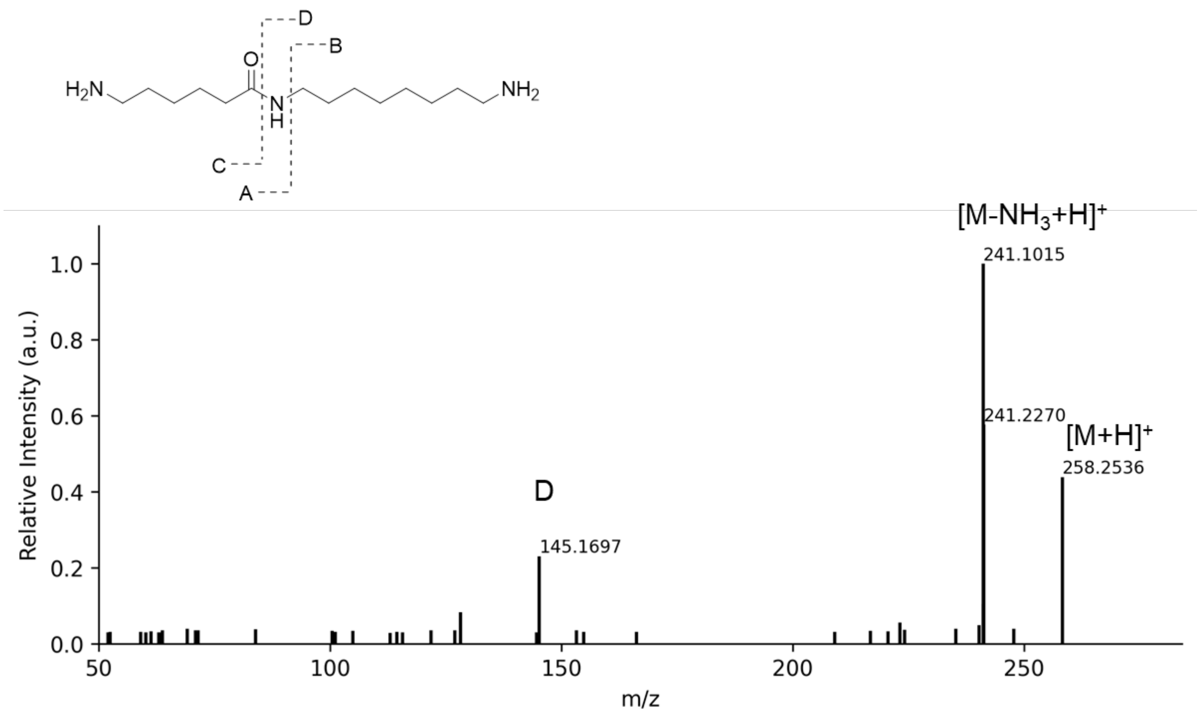


**Figure S11: I.DOT/OPSI-MS$^2$ of ligation product of M and 1,8-octanediamine.** A collision energy of 35 eV was used.

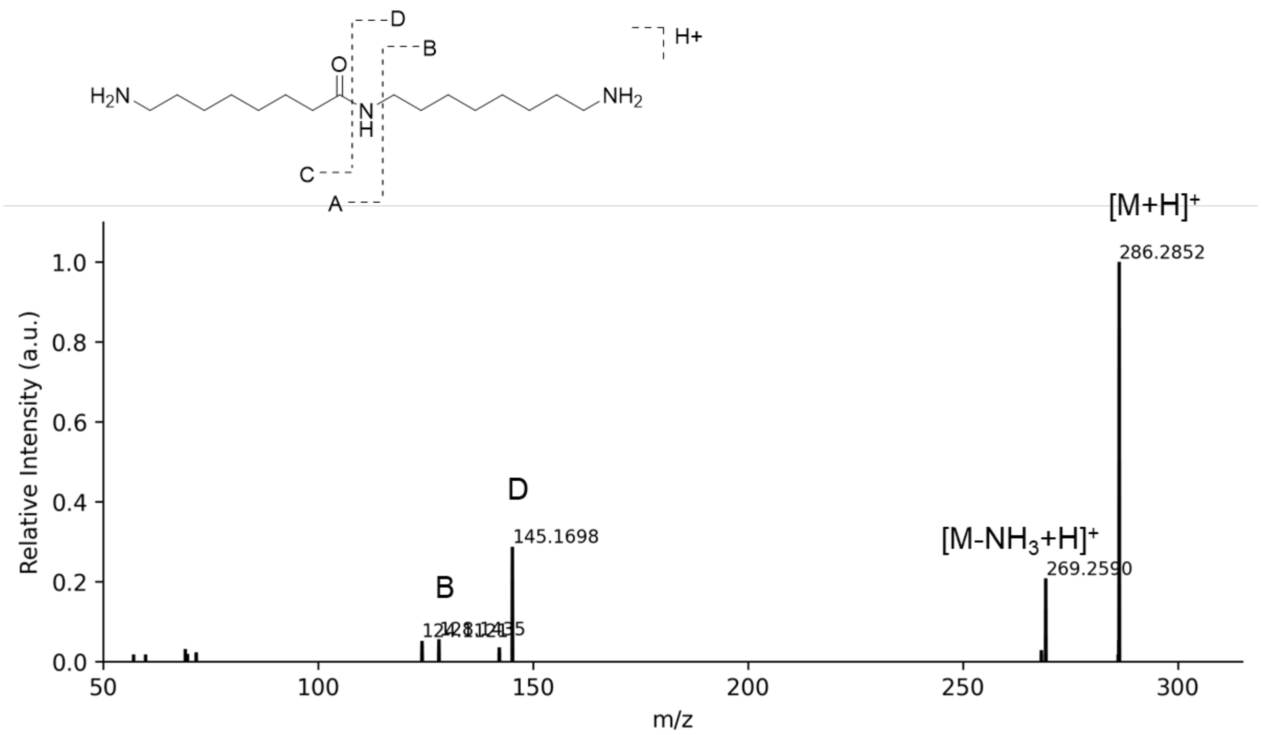


**Figure S12: I.DOT/OPSI-MS$^2$ of the ligation product of 8-aminooctanoic acid and 1,8-octanediamine.** A collision energy of 35 eV was used.

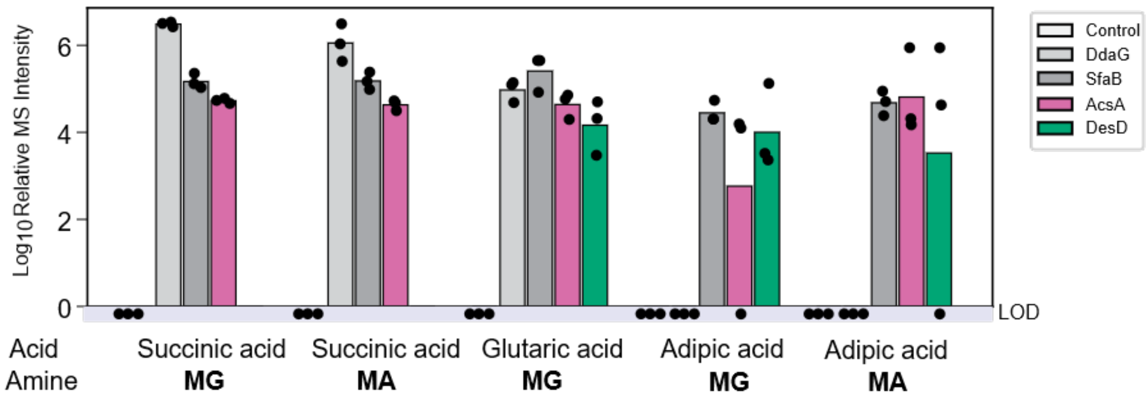


**Figure S13**. **Amide synthetases can synthesize diverse diacid triads**. *In vitro* biochemical assays were conducted by incubating different diacids (i.e., succinic acid, glutaric acid or adipic acid) with ω-amino acid diads (i.e., **MG** and **MA**) in the presence of enzymes (i.e., DdaG, SfaB, AcsA or DesD) or no-enzyme control. Symmetrical and asymmetrical diacid triad products were assayed using OPSI-MS. All replicates shown are biological replicates. Sample size is $N$=3. LOD: limit of detection.

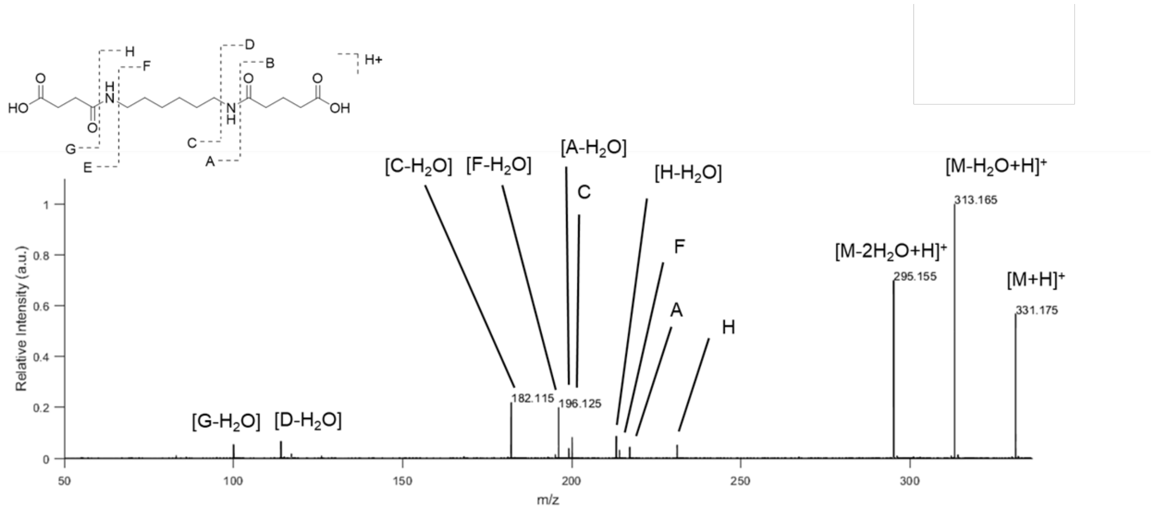


**Figure S14: I.DOT/OPSI-MS$^2$ of SMG.**

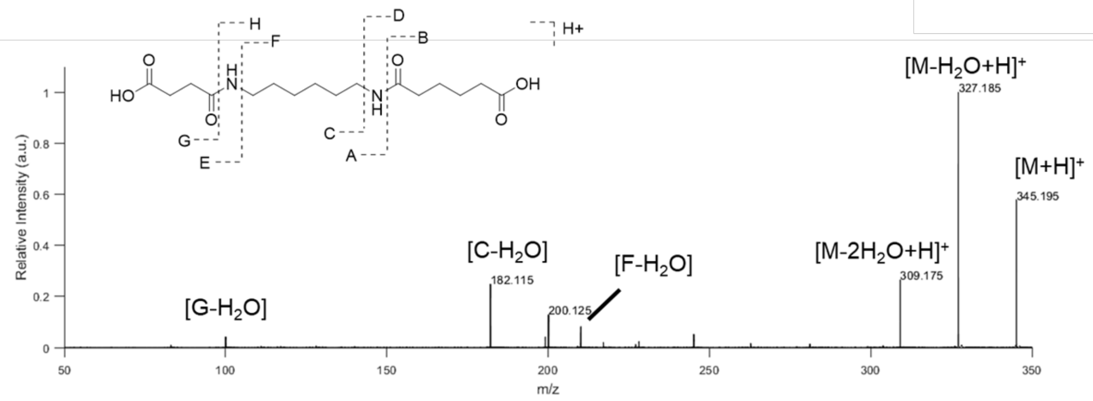


**Figure S15: I.DOT/OPSI-MS[2] of SMA.**

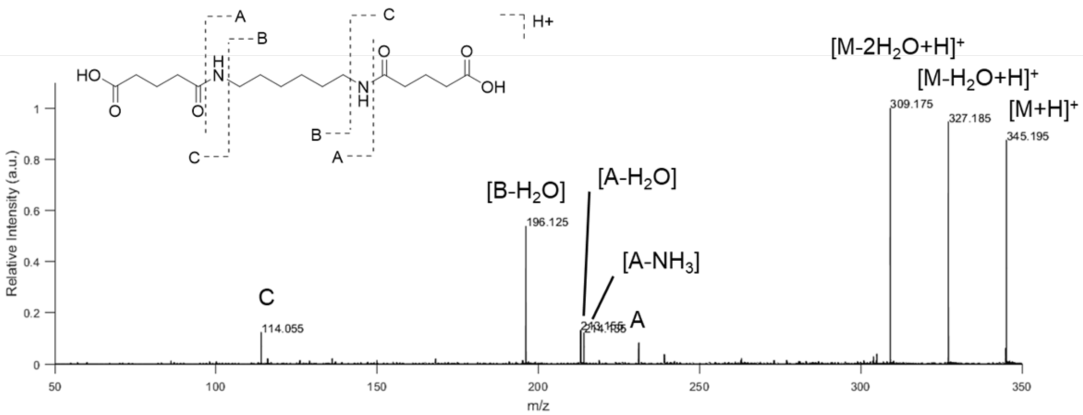


**Figure S16: I.DOT/OPSI-MS$^2$ of GMG.**

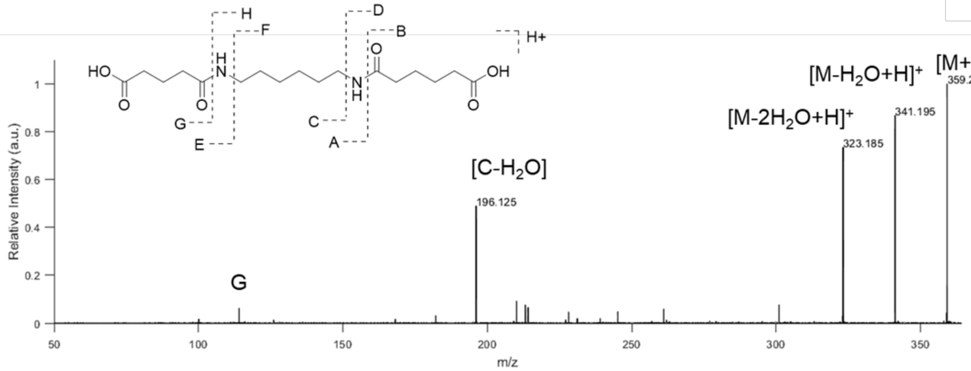


**Figure S17: I.DOT/OPSI-MS[2] of GMA.**

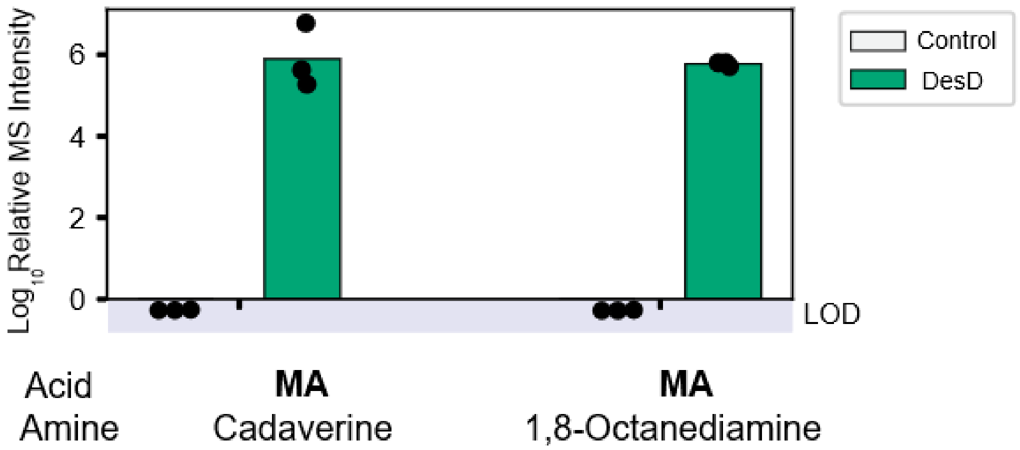


**Figure S18**. **Amide synthetases can synthesize diamine triads**. *In vitro* biochemical assays were conducted by incubating **MA** with diamines (cadaverine or 1,8-octanediamine) in the presence of DesD or no-enzyme control. Diamine triad products were assayed using OPSI-MS. All replicates shown are biological replicates. Sample size is *N*=3. LOD: limit of detection.

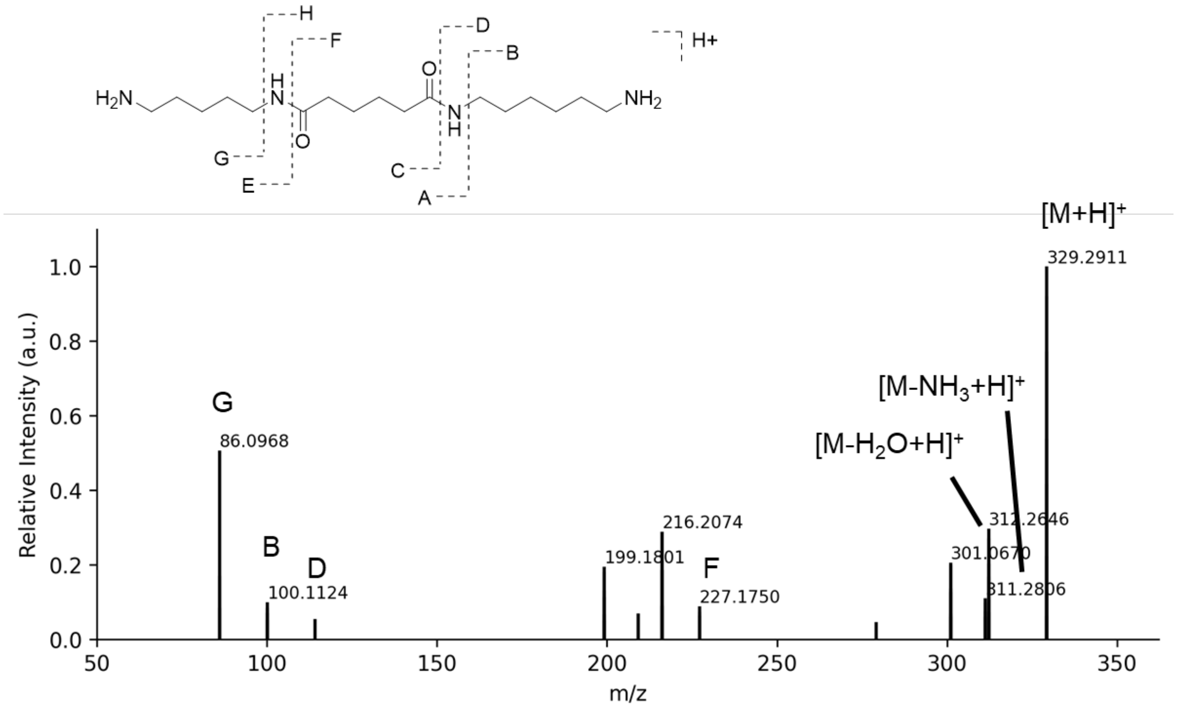


**Figure S19: I.DOT/OPSI-MS$^2$ of MAC.** A collision energy of 35 eV was used.

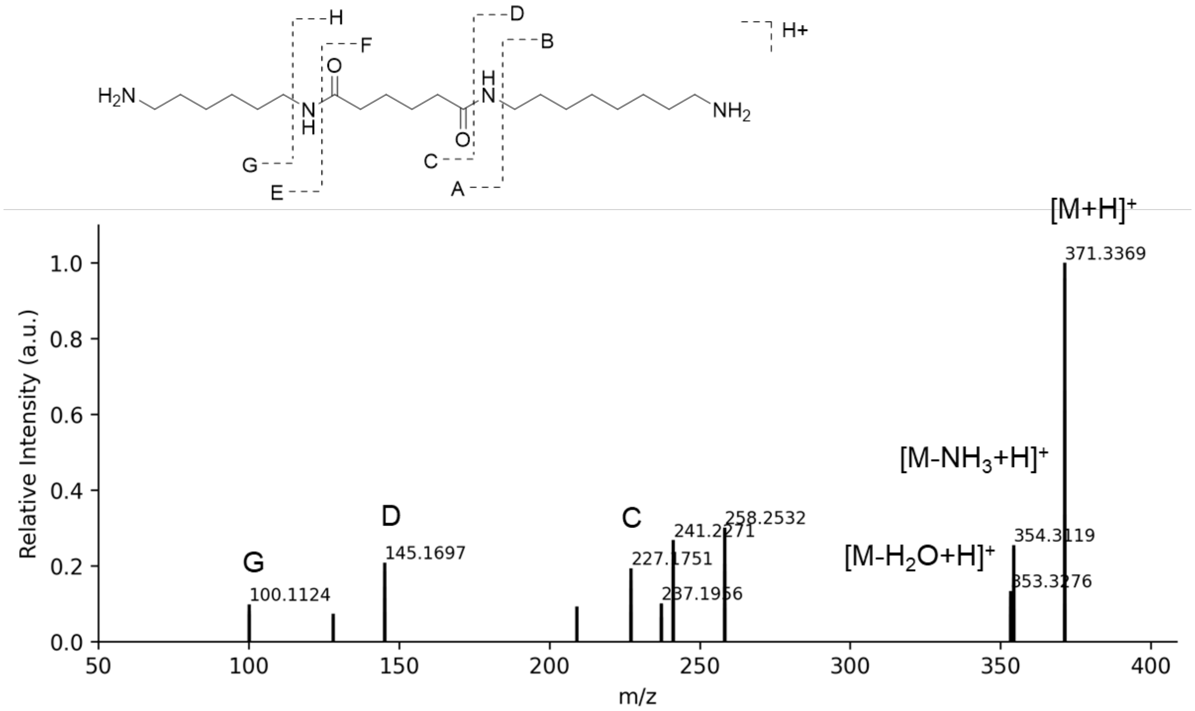


**Figure S20: I.DOT/OPSI-MS[2] of the ligation product of MA and 1,8-octanediamine.** A collision energy of 35 eV was used.

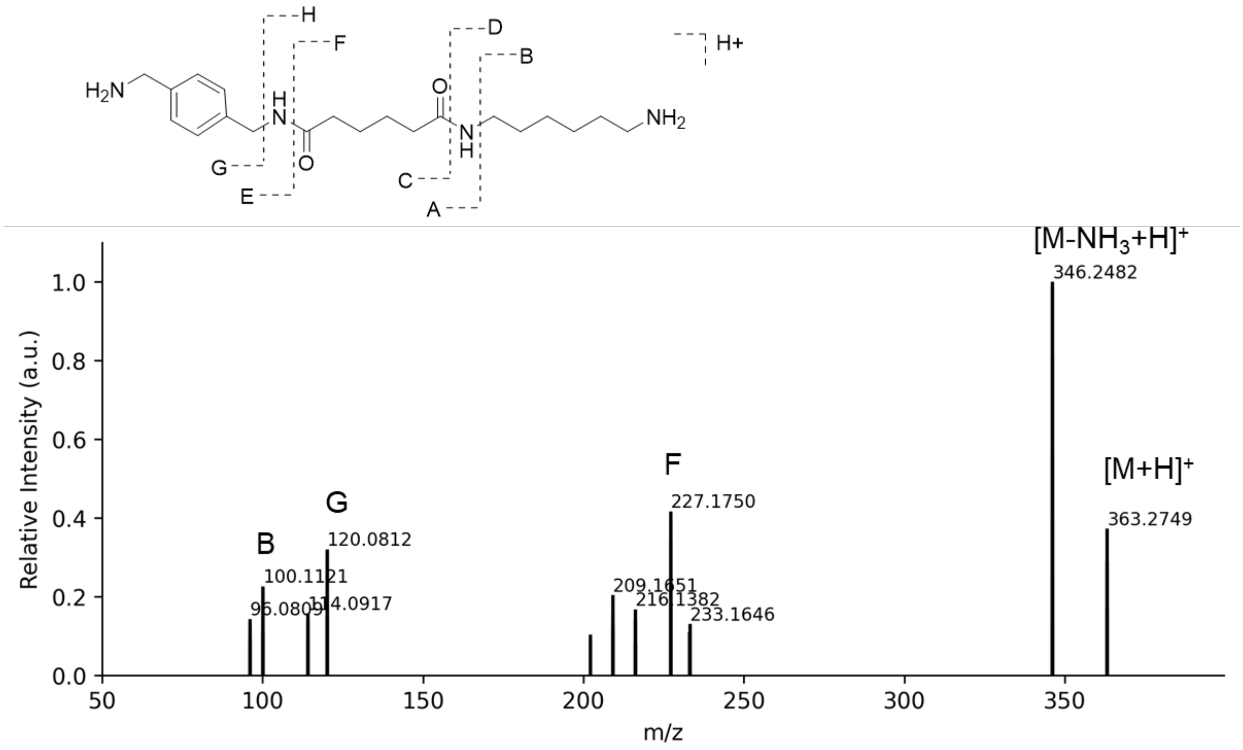


**Figure S21: I.DOT/OPSI-MS² of MAX.** A collision energy of 35 eV was used.

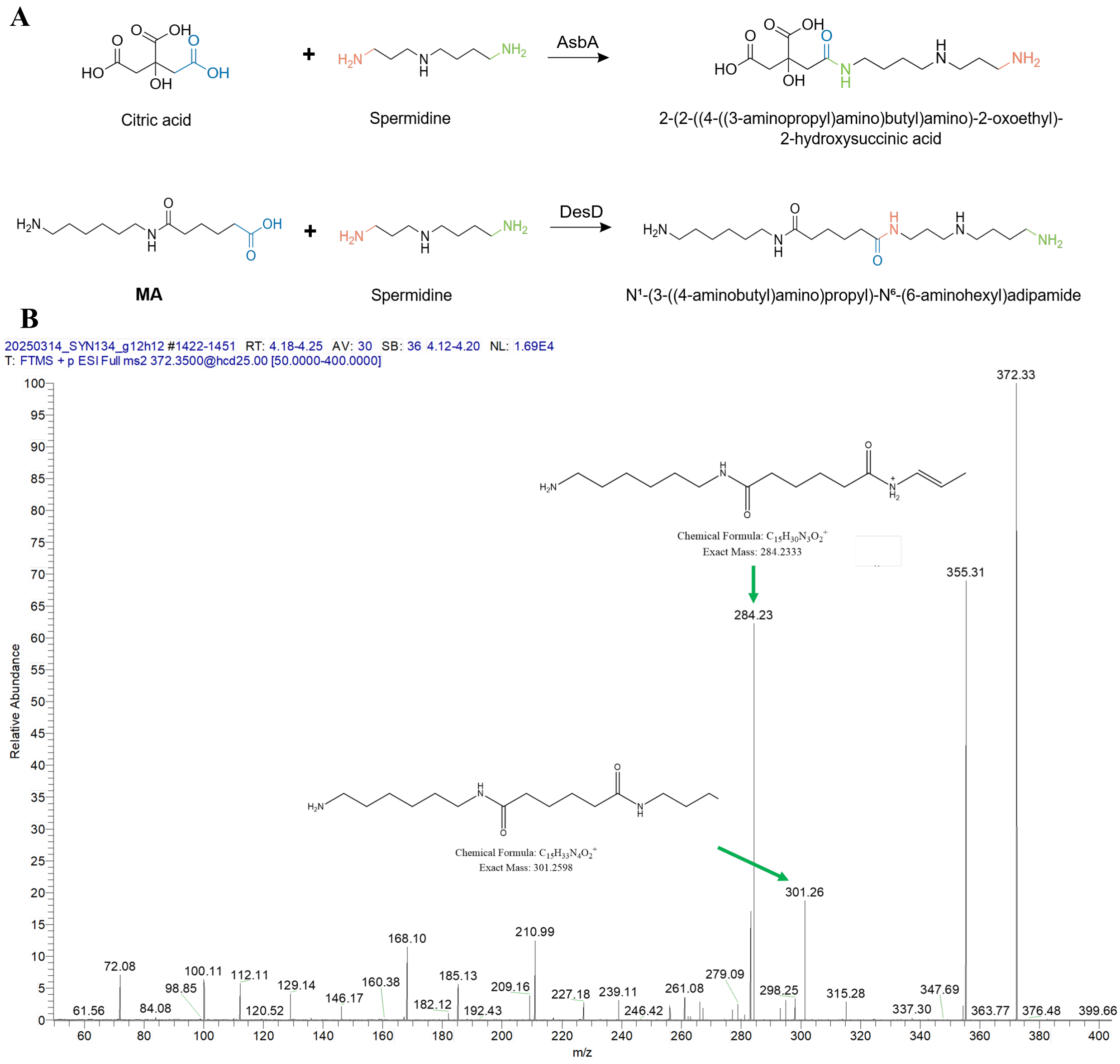


**Figure S22**. **Different regioselectivity of DesD and AsbA.** A) In the reactions with spermidine, DesD preferentially catalyzes amide bond formation at the amine near the secondary amine, whereas AsbA favors ligation at the distal primary amine. B) MS/MS spectrum of the ligation product formed between spermidine and the **MA** diad using DesD.

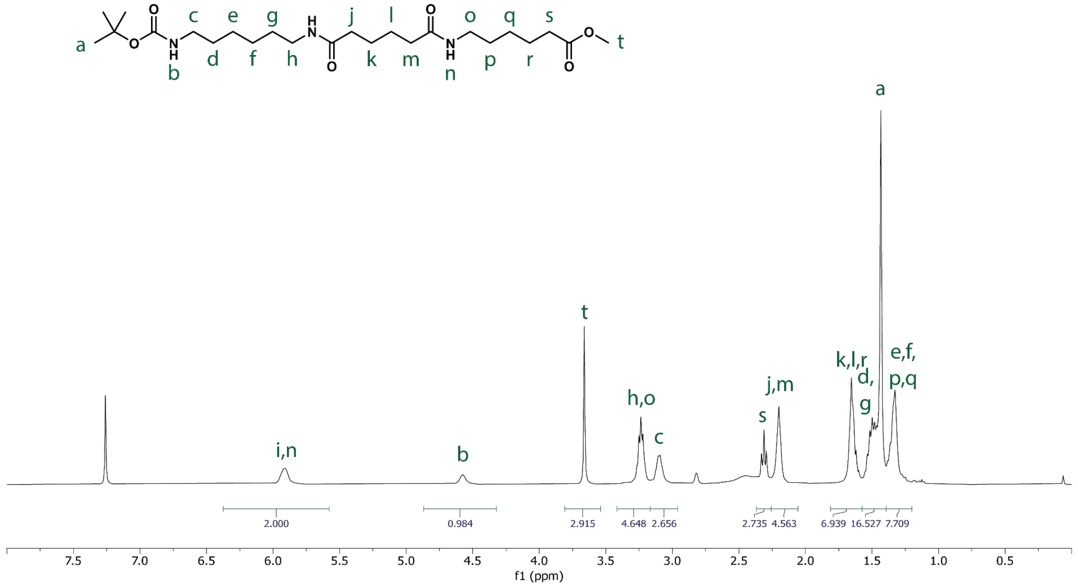


**Figure S23: $^{1}$H NMR spectrum of Boc-MAH-OMe in $CDCl_3$.**

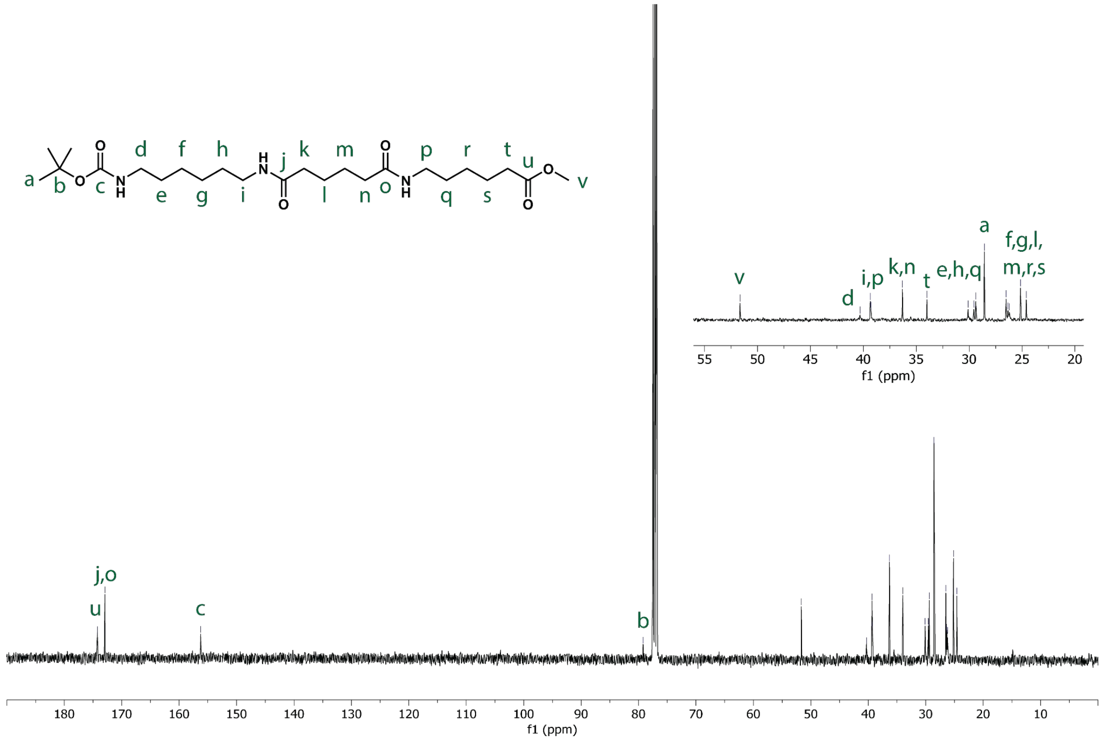


**Figure S24: $^{13}$C NMR spectrum of Boc-MAH-OMe in $CDCl_3$.**

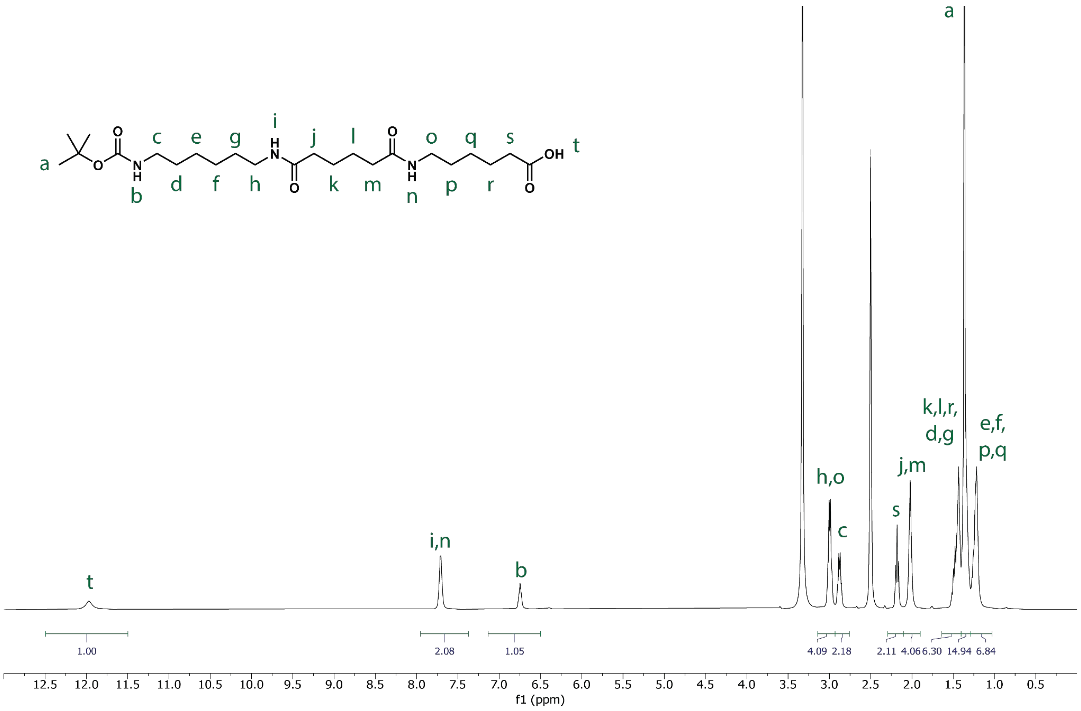


**Figure S25: $^{1}H$ NMR spectrum of Boc-MAH in DMSO-$d_6$.**

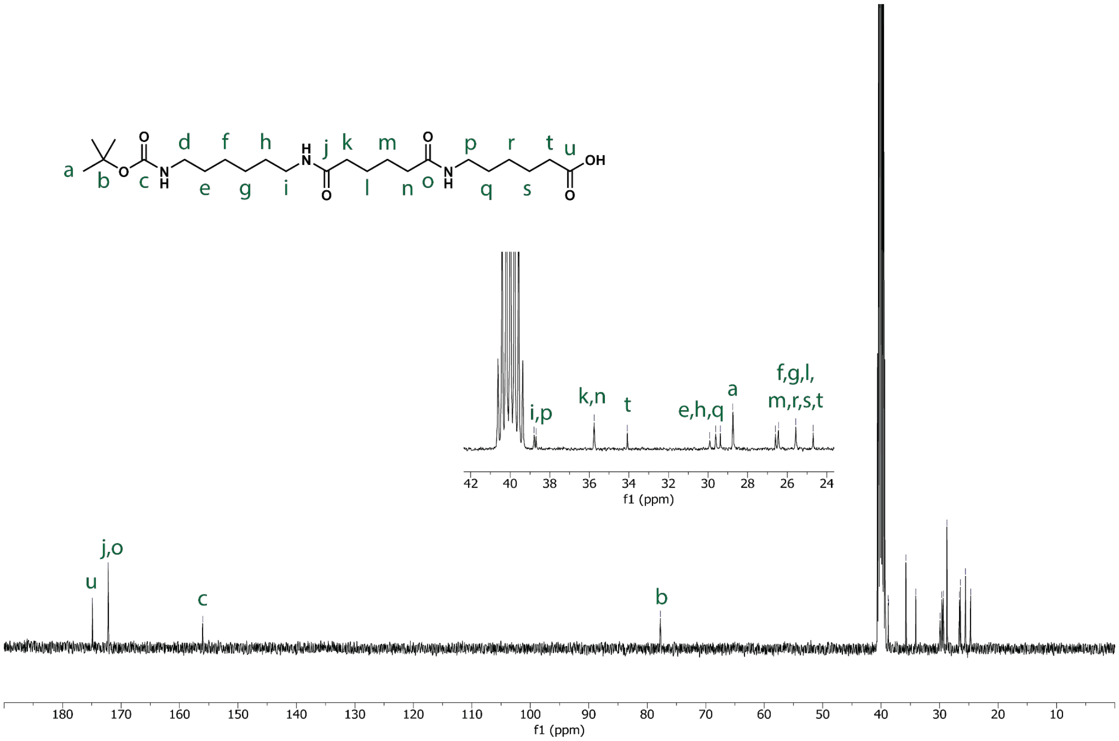


**Figure S26: $^{13}C$ NMR spectrum of Boc-MAH in DMSO-$d_6$.**

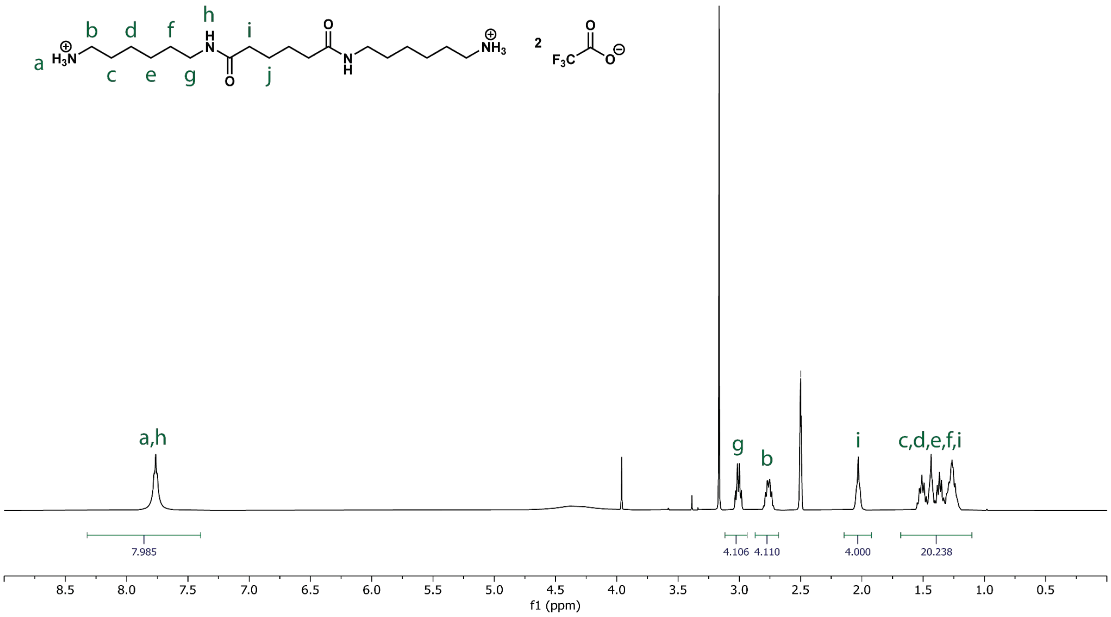


**Figure S27: $^{1}$H NMR spectrum of MAH trifluoroacetate in DMSO-d$_6$.**

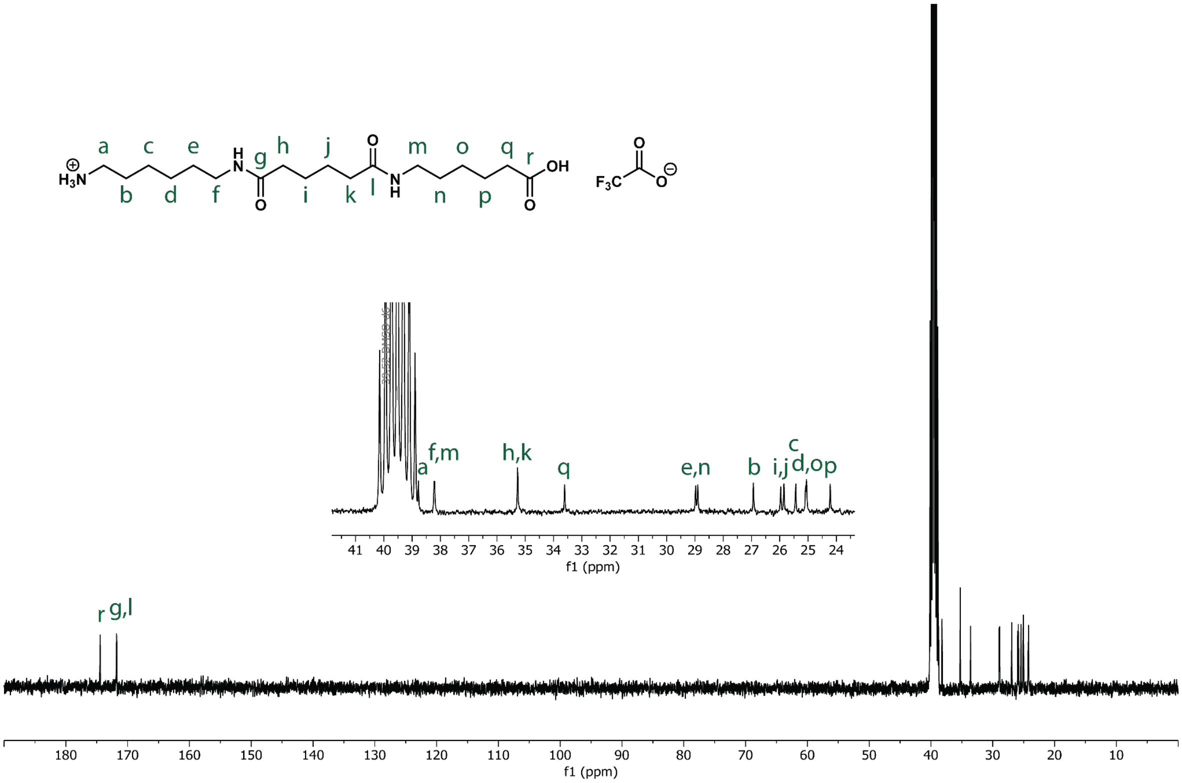


**Figure S28: $^{13}$C NMR spectrum of MAH trifluoroacetate in DMSO-d$_6$.**

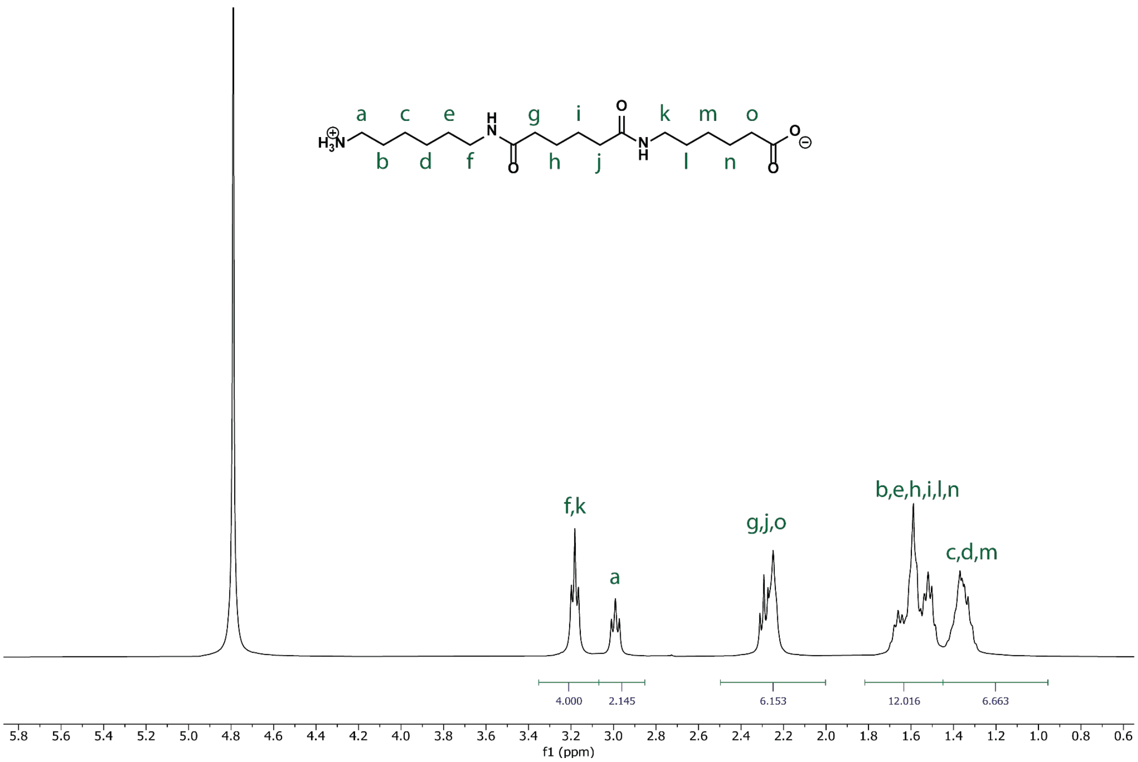


**Figure S29: $^{1}$H NMR spectrum of MAH in $D_2O$.**

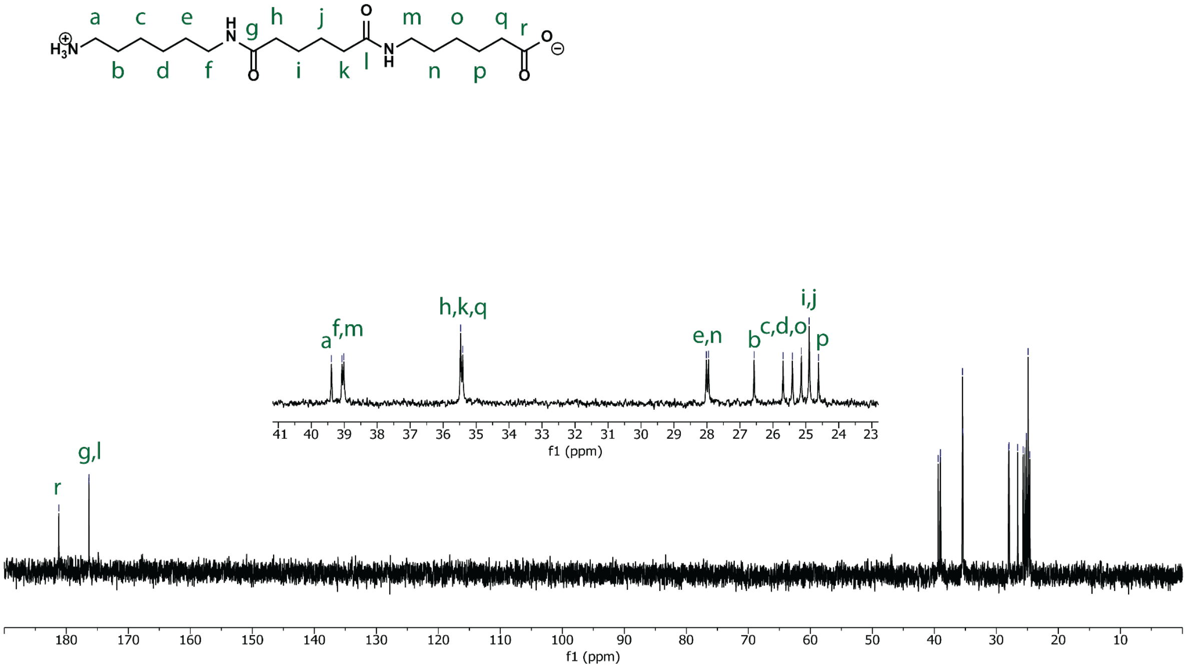


**Figure S30: $^{13}$C NMR spectrum of MAH in $D_2O$.**

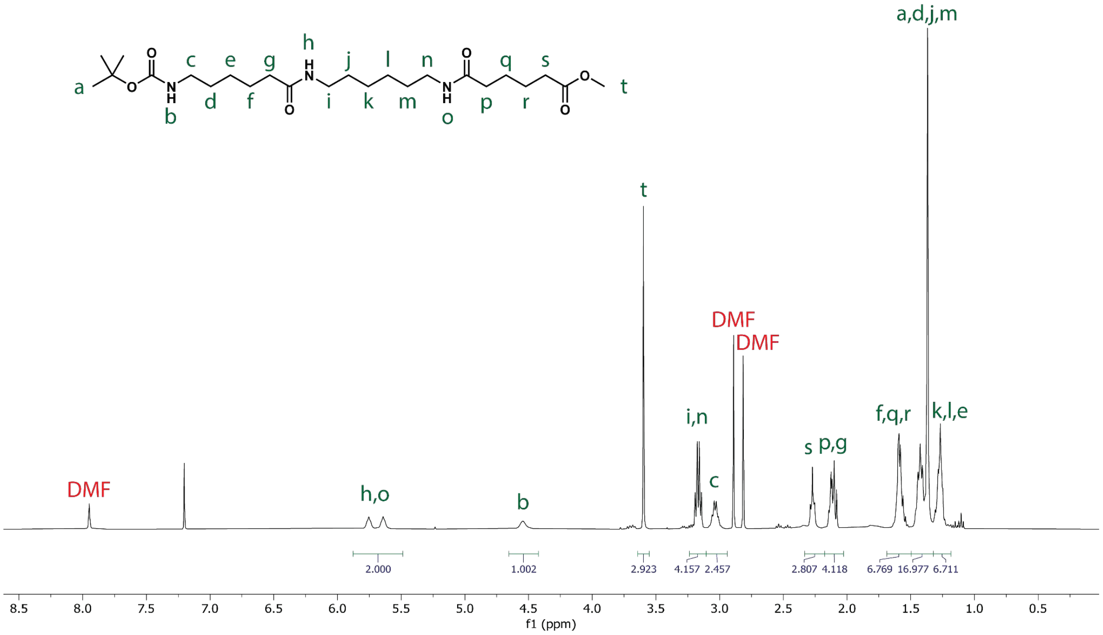


**Figure S31: $^1$H NMR spectrum of Boc-HMA-OMe in CDCl$_3$.**

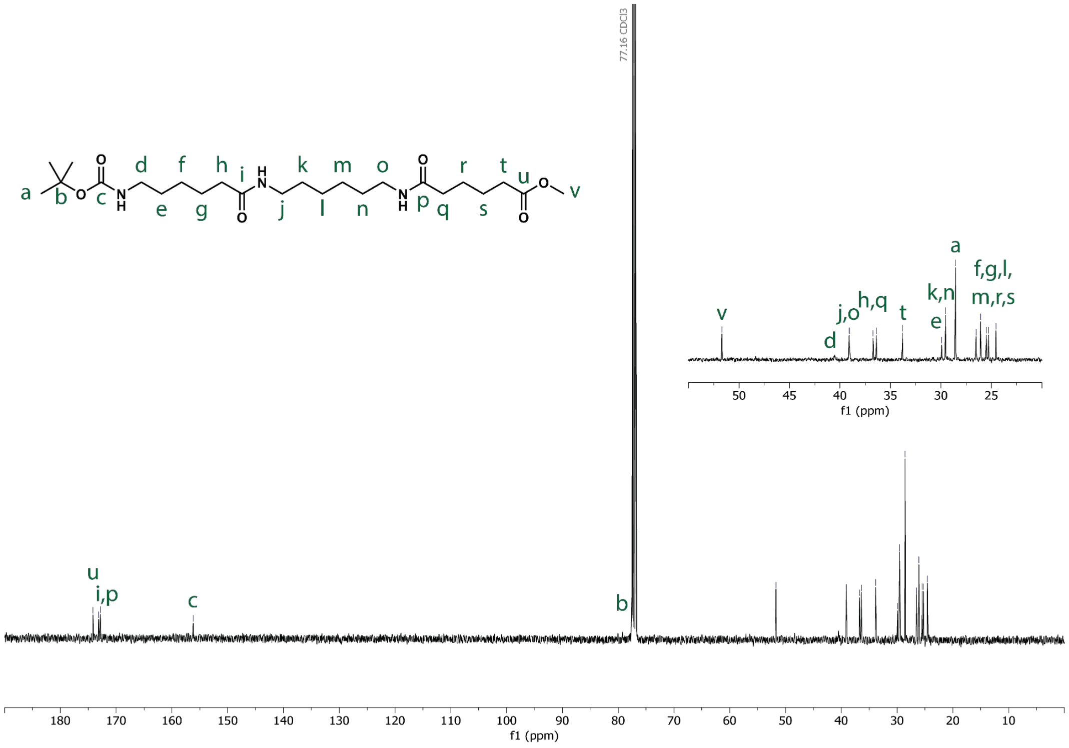


**Figure S32: $^{13}$C NMR spectrum of Boc-HMA-OMe in CDCl$_3$.**

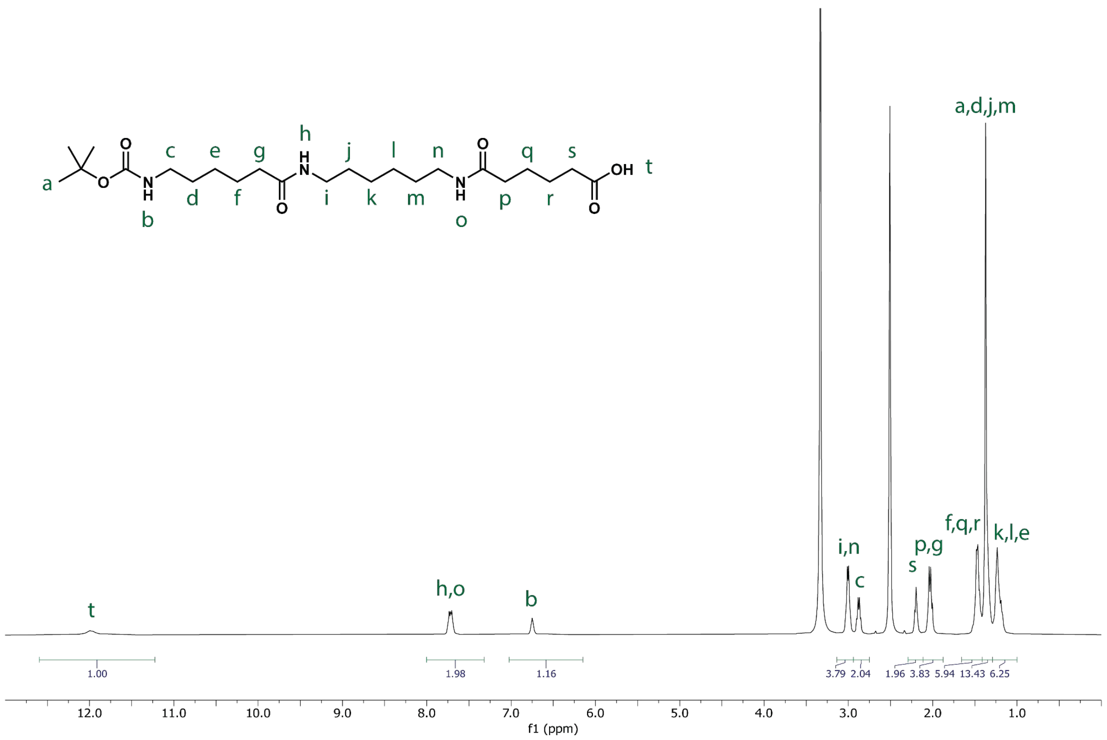


**Figure S33: $^{1}$H NMR spectrum of Boc-HMA in DMSO-d$_6$.**

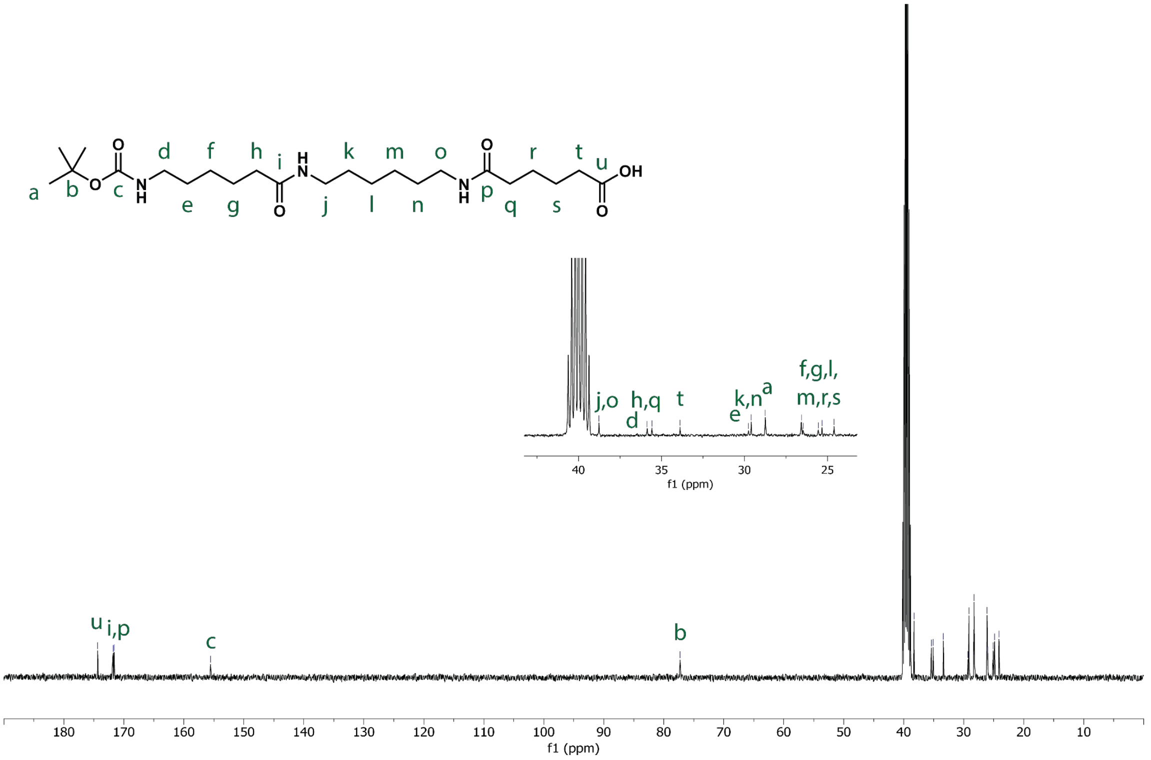


**Figure S34: $^{13}$C NMR spectrum of Boc-HMA in DMSO-d$_6$.**

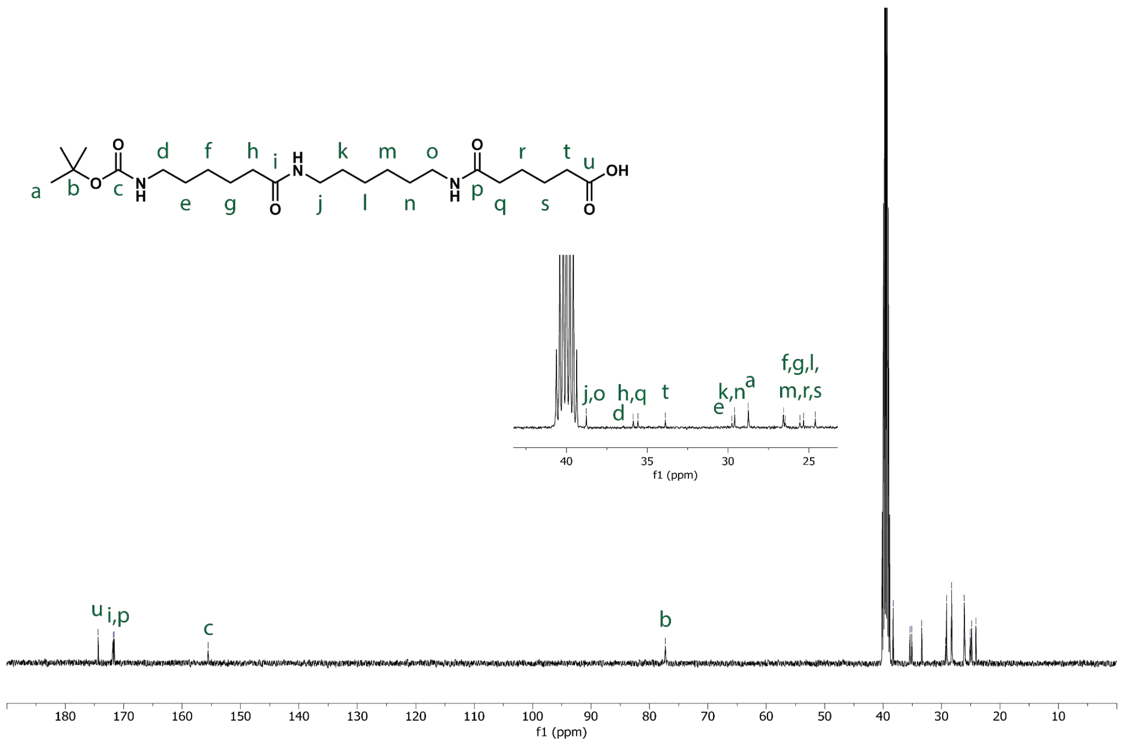


**Figure S35: $^{13}$C NMR spectrum of Boc-HMA in DMSO-d$_6$.**

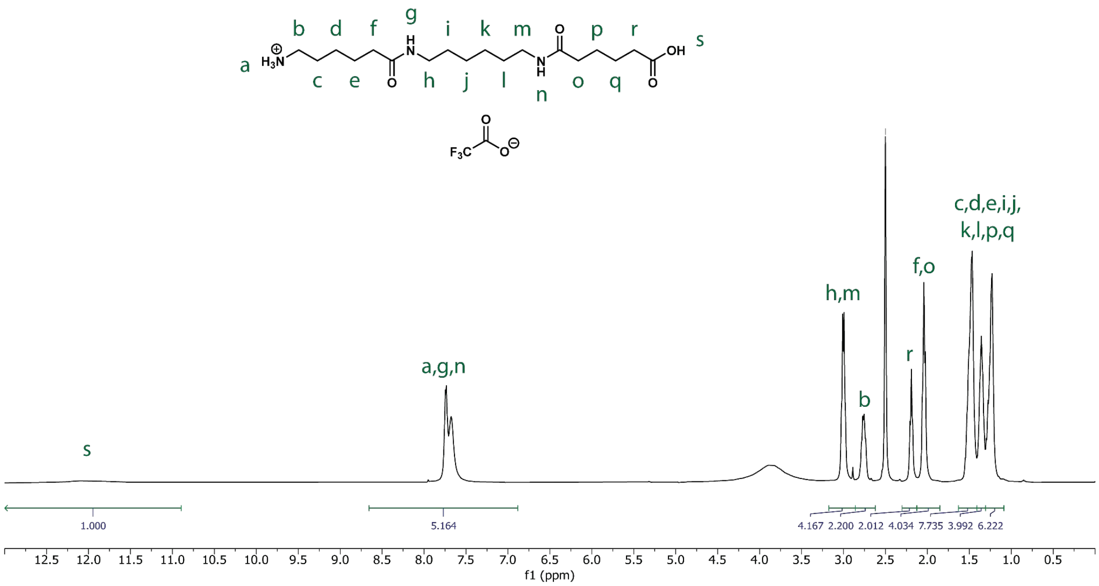


**Figure S36: $^{1}$H NMR spectrum of HMA trifluoroacetate in DMSO-d$_6$.**

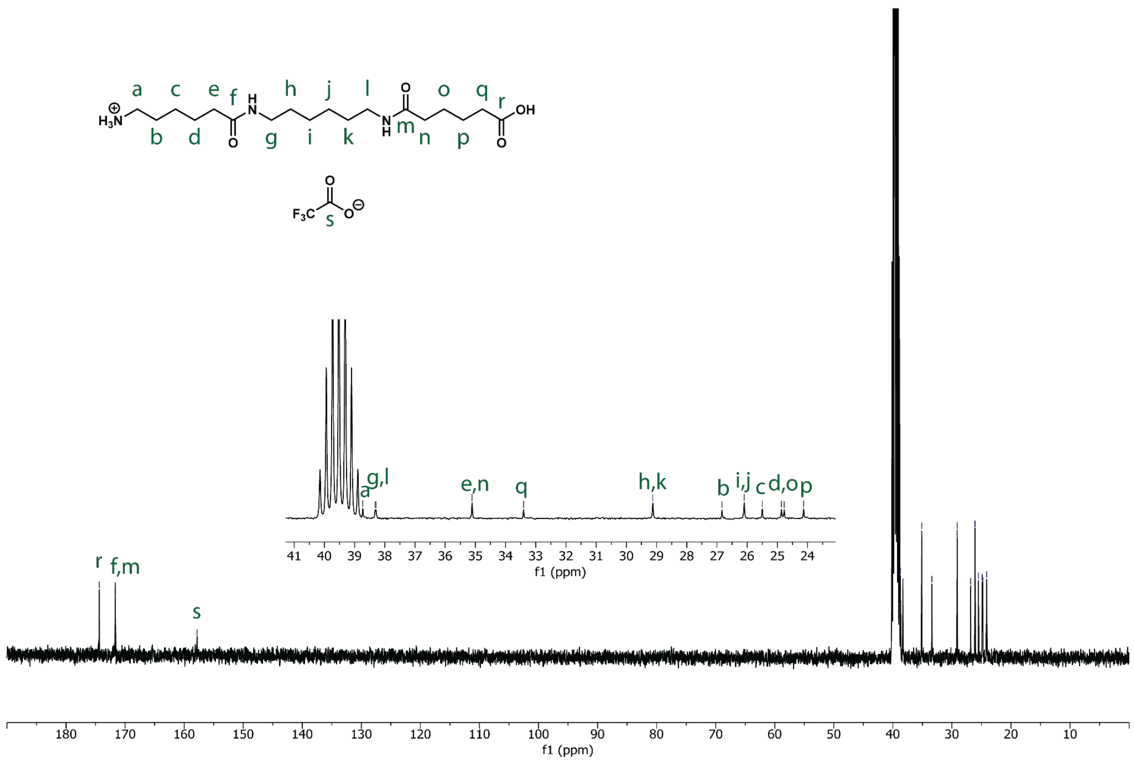


**Figure S37: [13]C NMR spectrum of HMA trifluoroacetate in DMSO-d$_6$.**

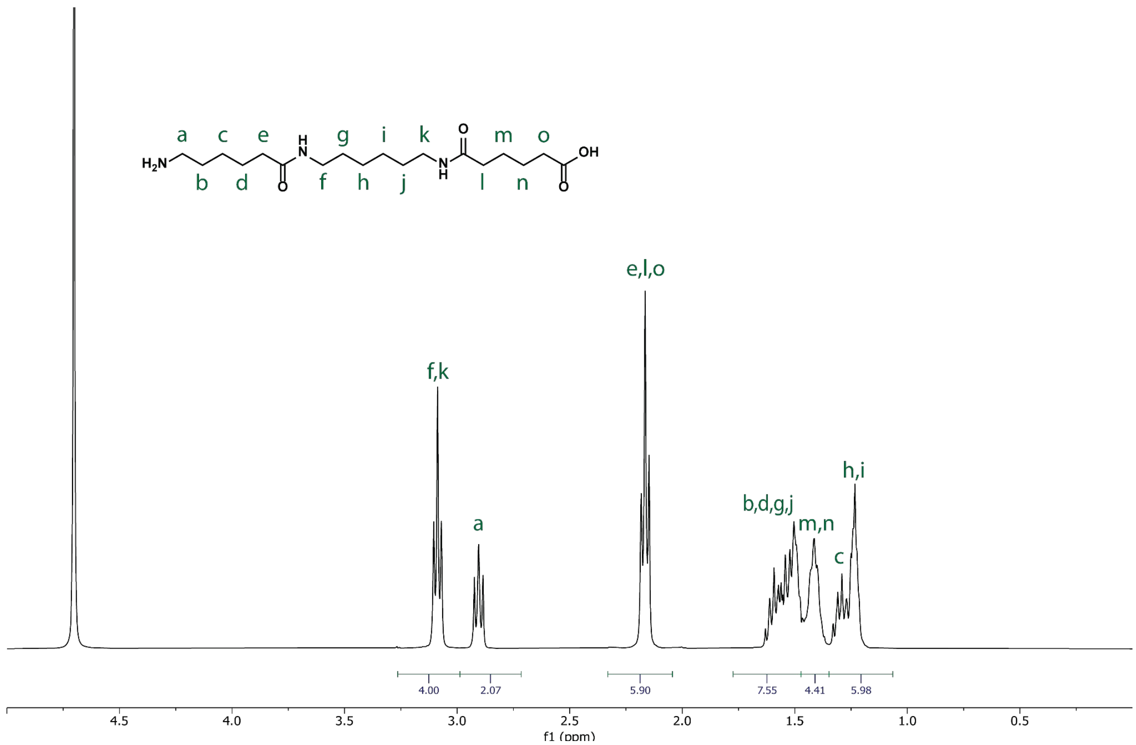


**Figure S38: [1]H NMR spectrum of HMA in D$_2$O.**

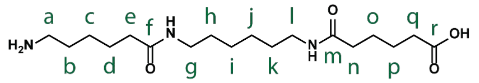

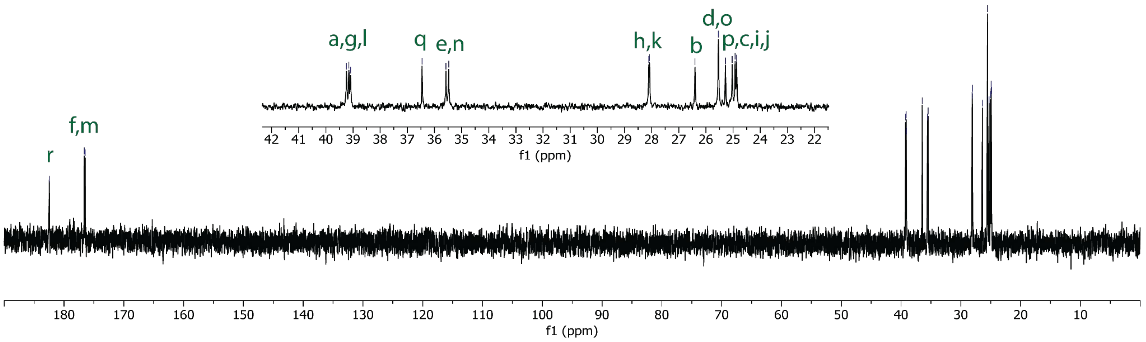


**Figure S39: $^{13}$C NMR spectrum of HMA in $D_2O$.**

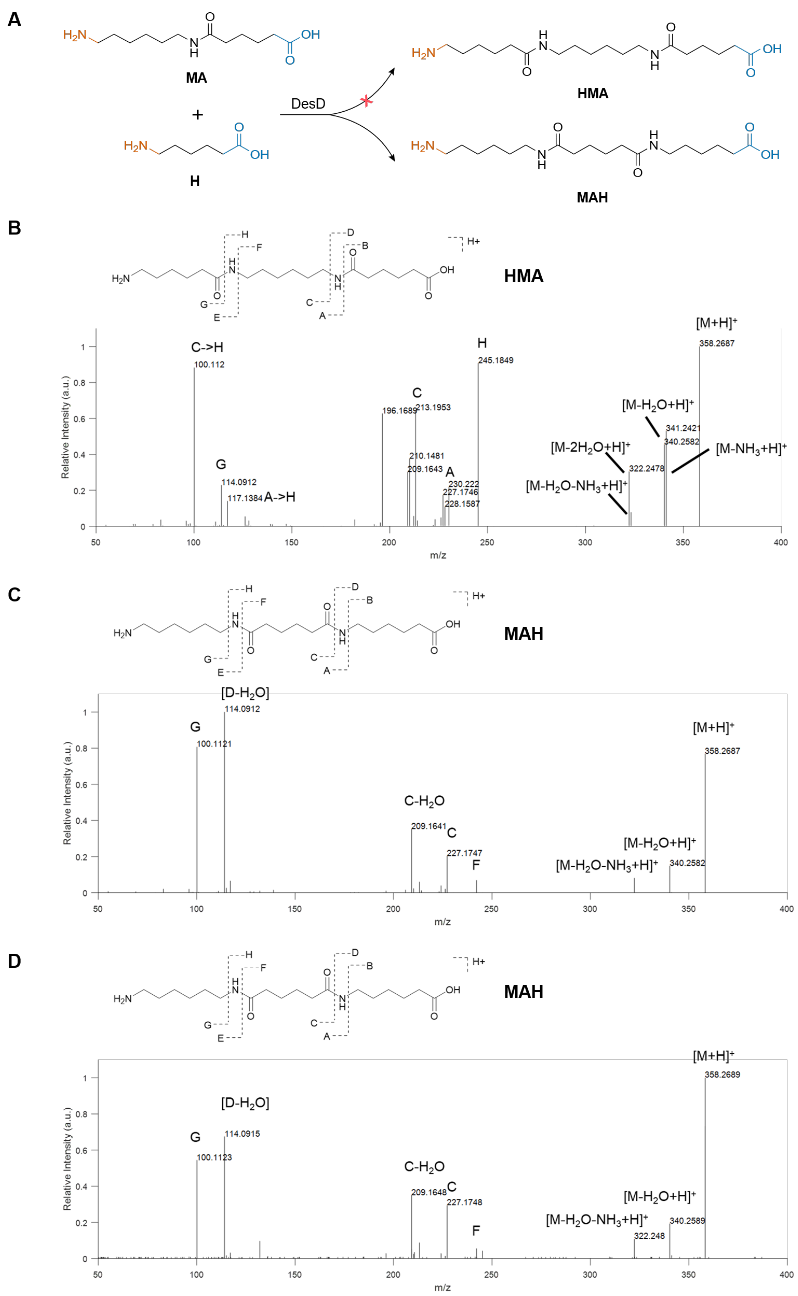


**Figure S40**. **DesD can synthesize ω-amino triads with regioselectivity.** A) In the reactions of **MA** diad and 6-aminocaproic acid **H**, DesD ligates the carboxylic group of **MA** and the amine group of **H** to form **MAH**. B) MS/MS spectrum of the ligation product **MAH**.

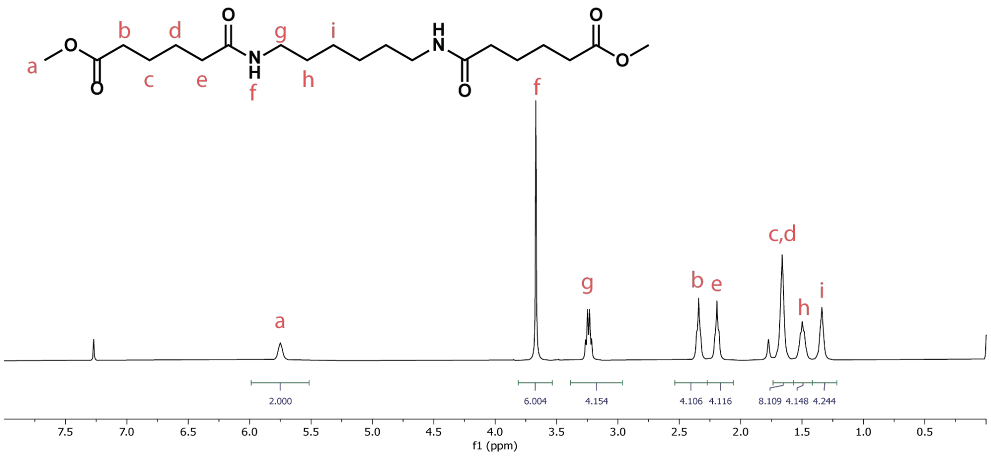


**Figure S41: $^{1}$H NMR spectrum of AMA-OMe in $CDCl_3$.**

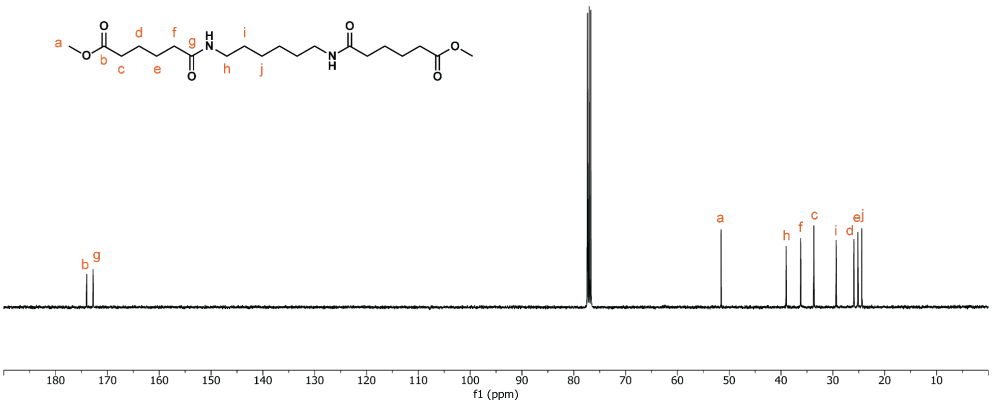


**Figure S42: $^{1}$H NMR spectrum of AMA-OMe in $CDCl_3$.**

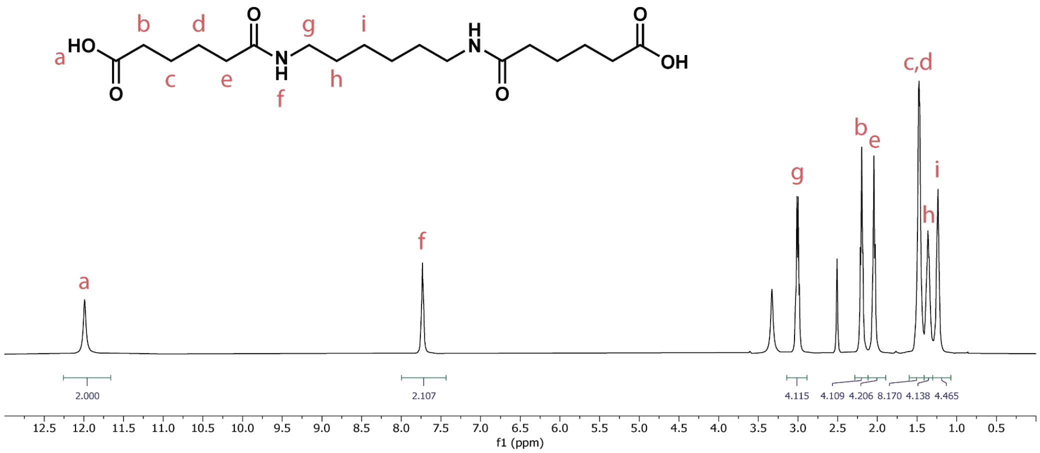


**Figure S43: ¹H NMR spectrum of AMA in DMSO-d₆.**

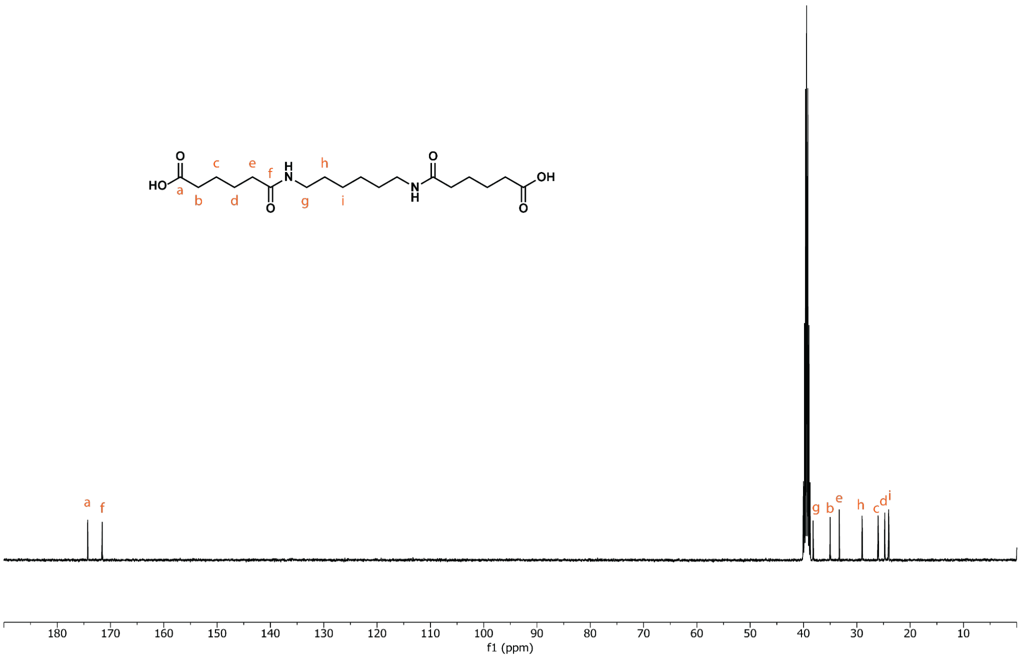


**Figure S44: ¹³C NMR spectrum of AMA in DMSO-d₆.**

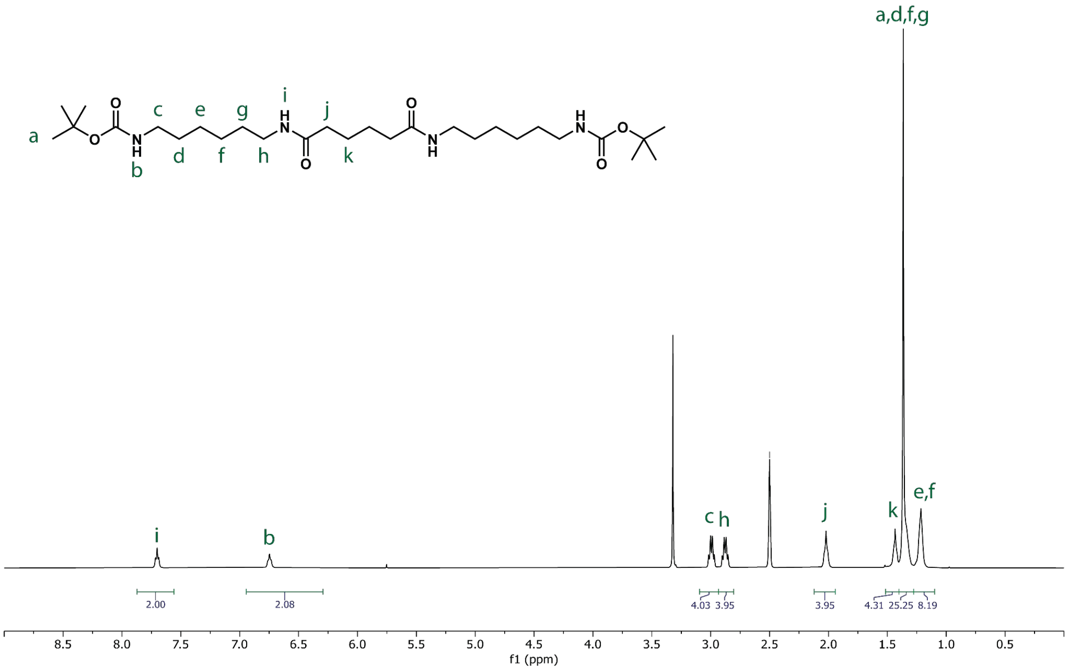


**Figure S45: $^1$H NMR spectrum of Boc-MAM in DMSO-d$_6$.**

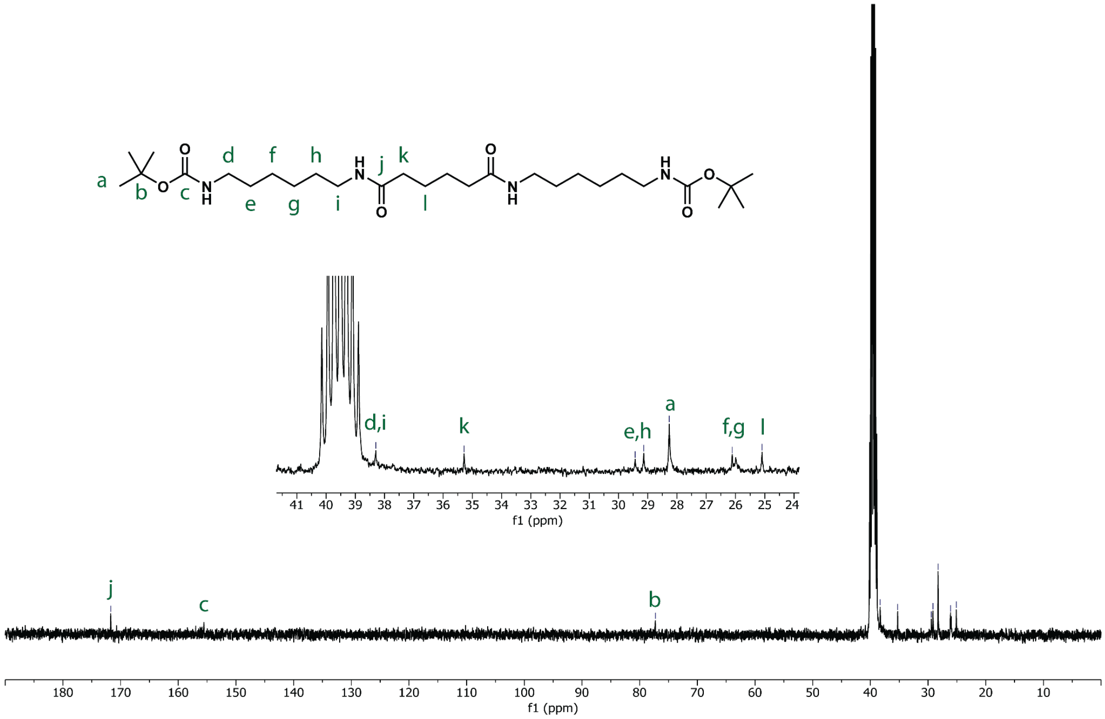


**Figure S46: $^{13}$C NMR spectrum of Boc-MAM in DMSO-d$_6$.**

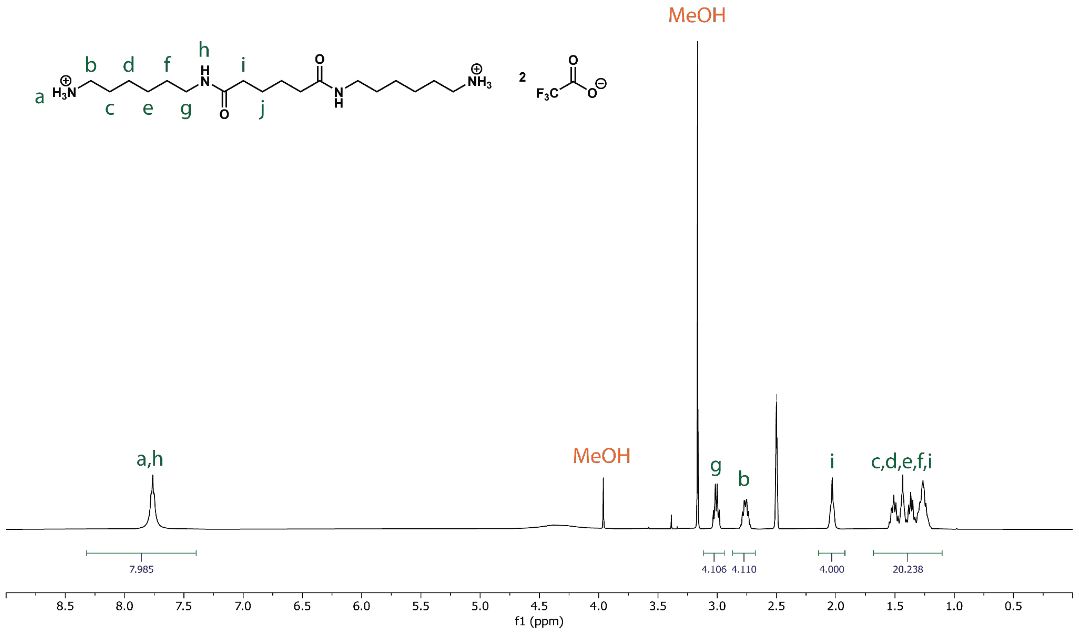


**Figure S47: $^{1}H$ NMR spectrum of MAM trifluoroacetate in DMSO-d$_6$.**

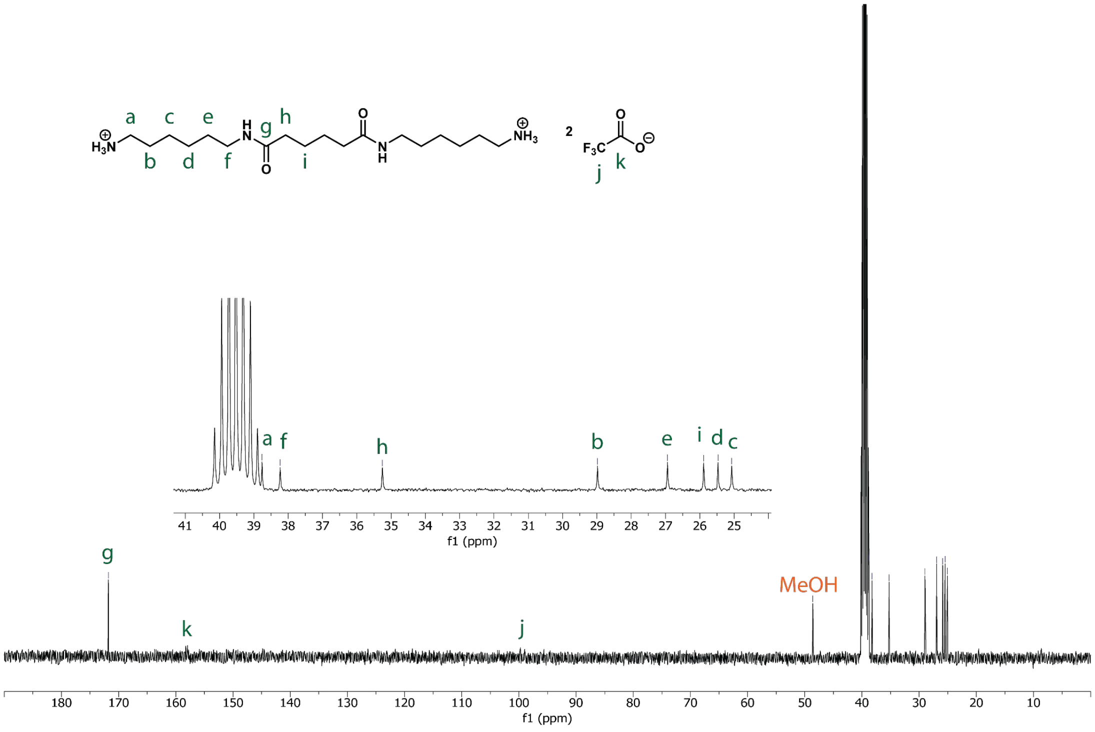


**Figure S48: $^{13}C$ NMR spectrum of MAM trifluoroacetate in DMSO-d$_6$.**

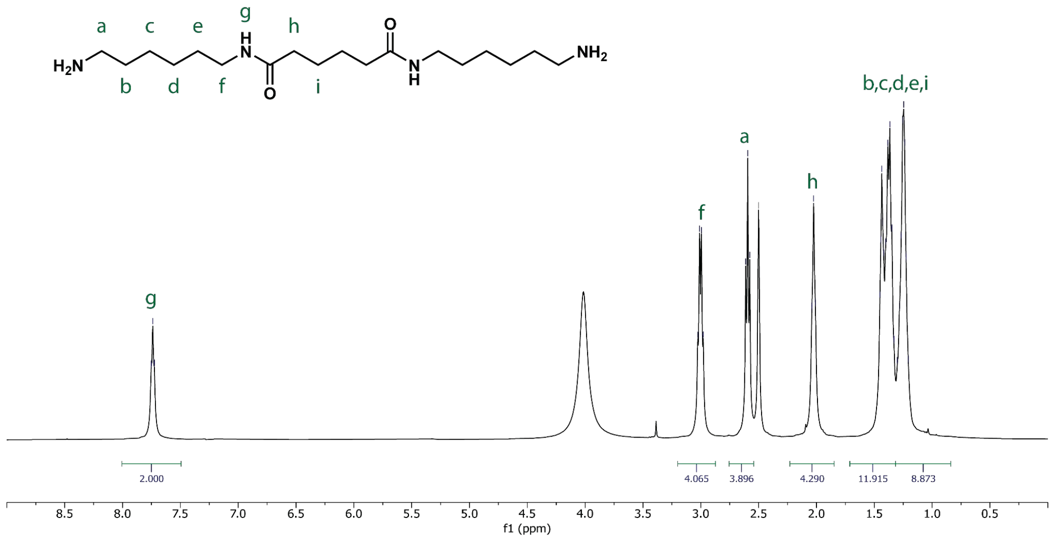


Figure S49: $^{1}$H NMR spectrum of MAM in $D_2O$.

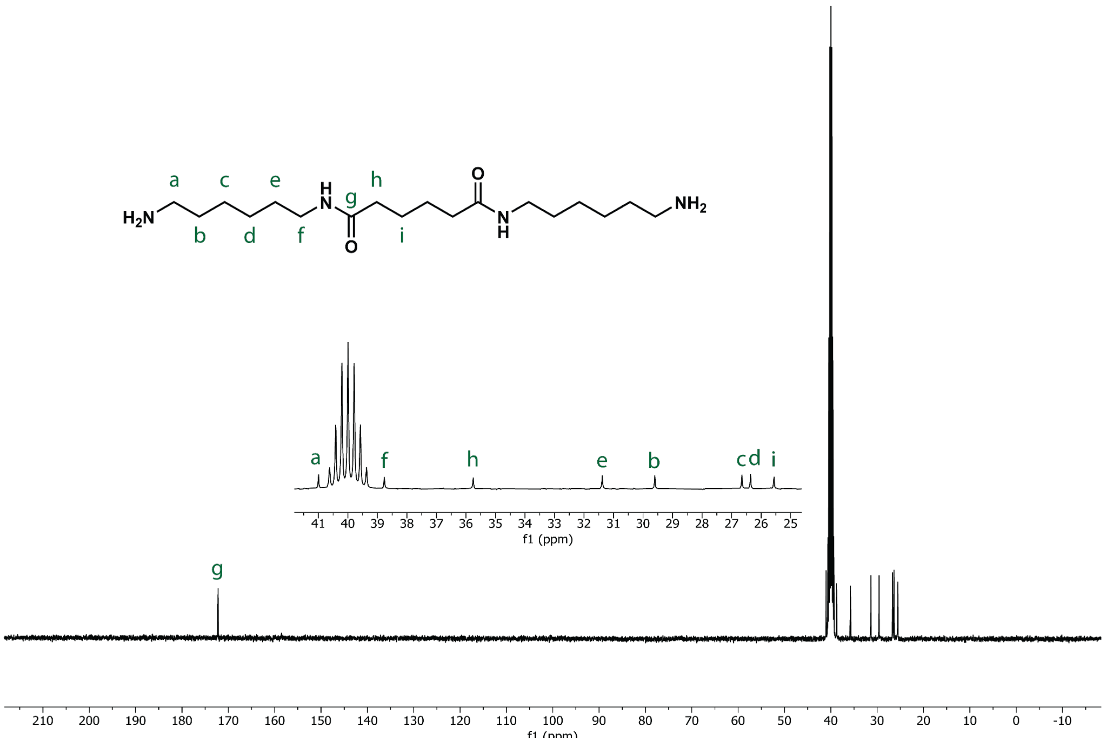


Figure S50: $^{13}$C NMR spectrum of MAM in DMSO-$d_6$.

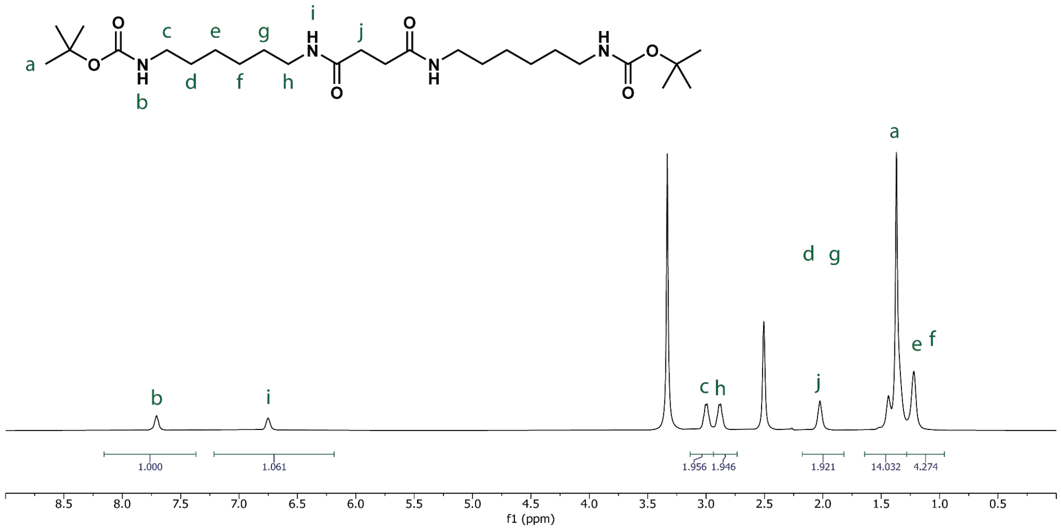


**Figure S51: [1]H NMR spectrum of Boc-MSM in DMSO-d6.**

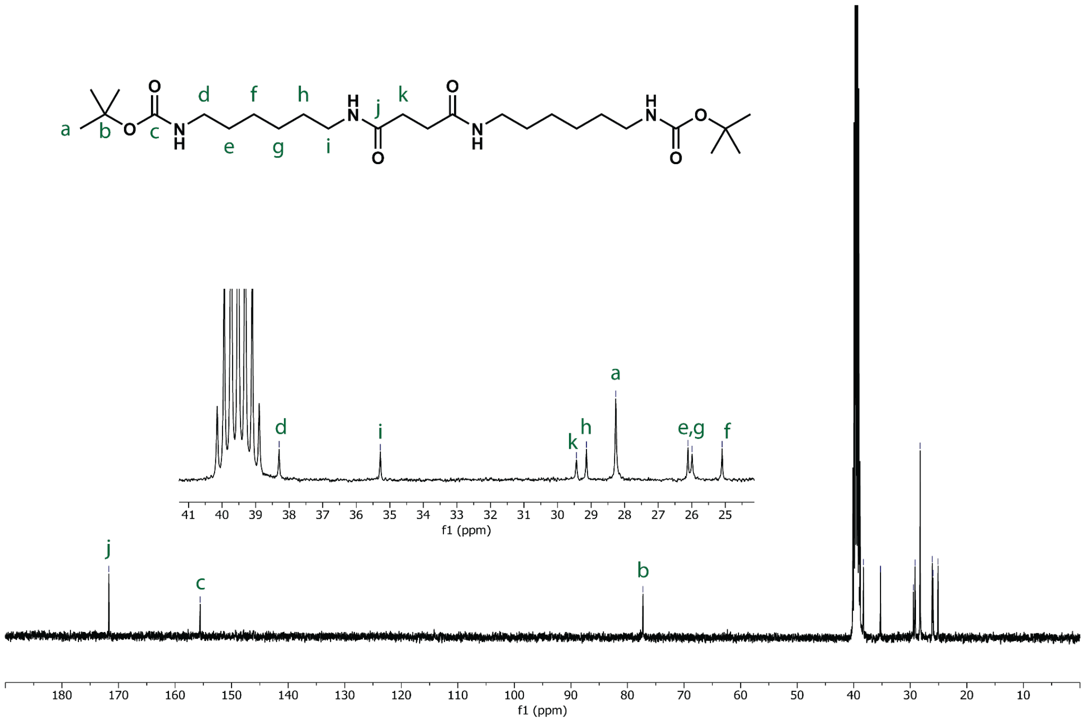


**Figure S52: [13]C NMR spectrum of Boc-MSM in DMSO-d6.**

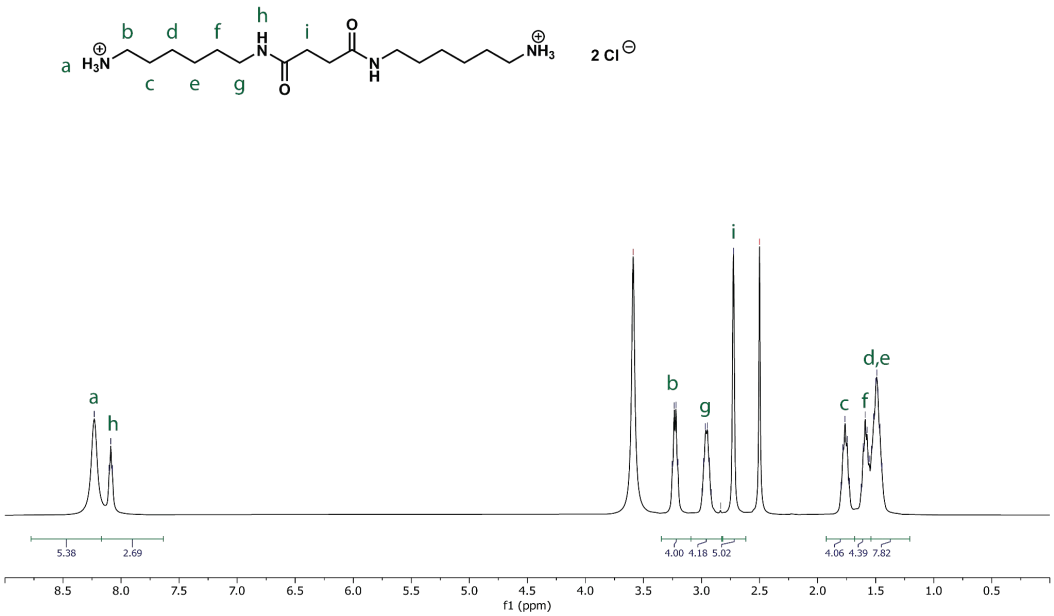


**Figure S53: $^{1}$H NMR spectrum of MSM HCl in DMSO-d$_6$.**

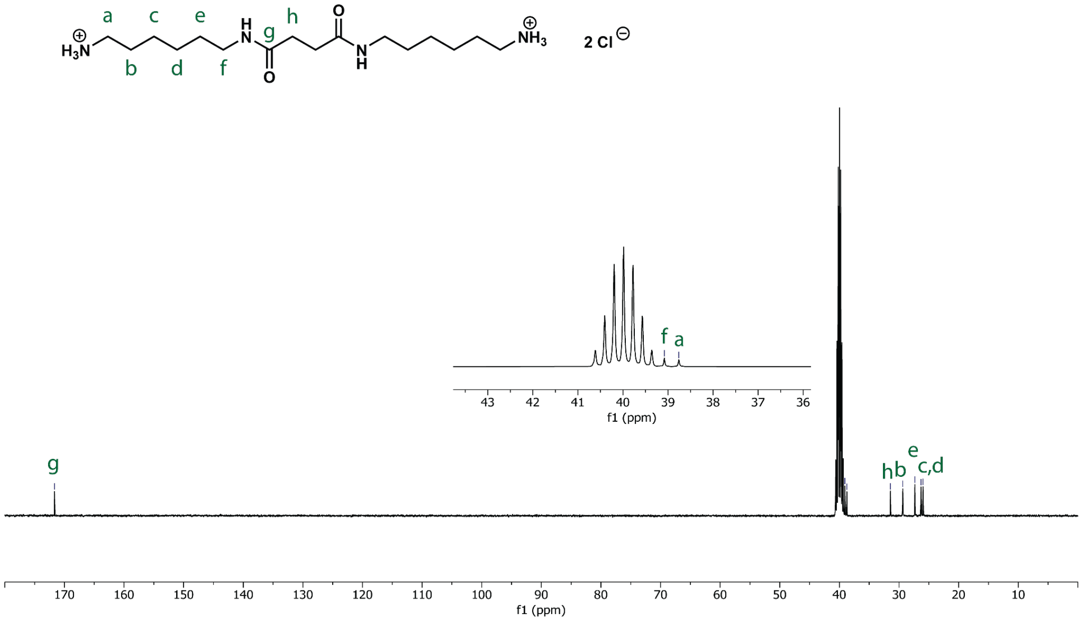


**Figure S54: $^{13}$C NMR spectrum of MSM HCl in DMSO-d$_6$.**

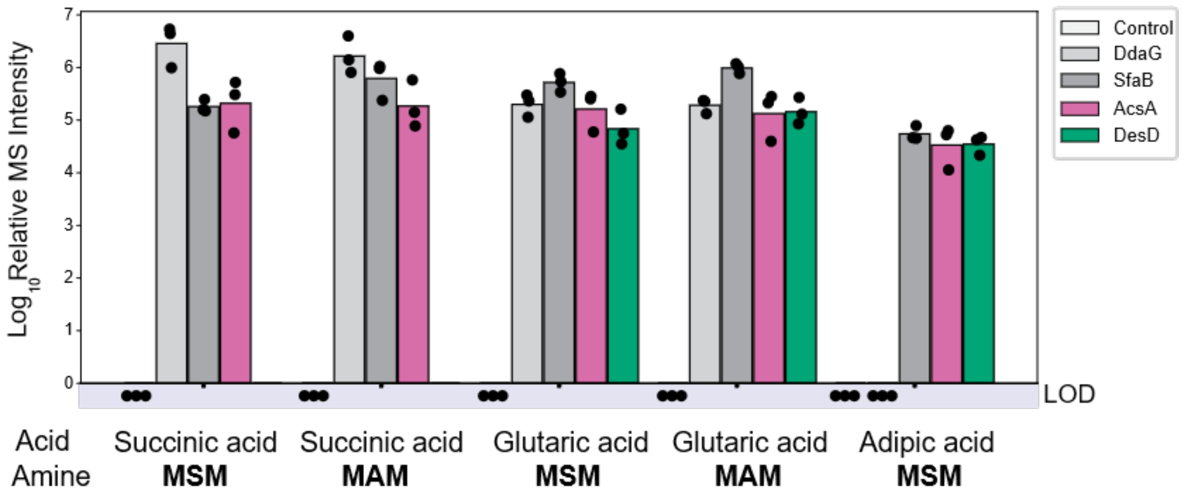


**Figure S55**. **Amide synthetases can synthesize ω-amino acid tetrads**. *In vitro* biochemical assays were conducted by incubating diamine triads **MSM** or **MAM** with diacids (i.e., succinic acid, glutaric acid or adipic acid) in the presence of enzymes (i.e., DdaG, SfaB, AsbA, AsbB, AcsA and DesD) or no-enzyme control. Products were assayed using OPSI-MS. All replicates shown are biological replicates. Sample size is $N$=3. LOD: limit of detection.

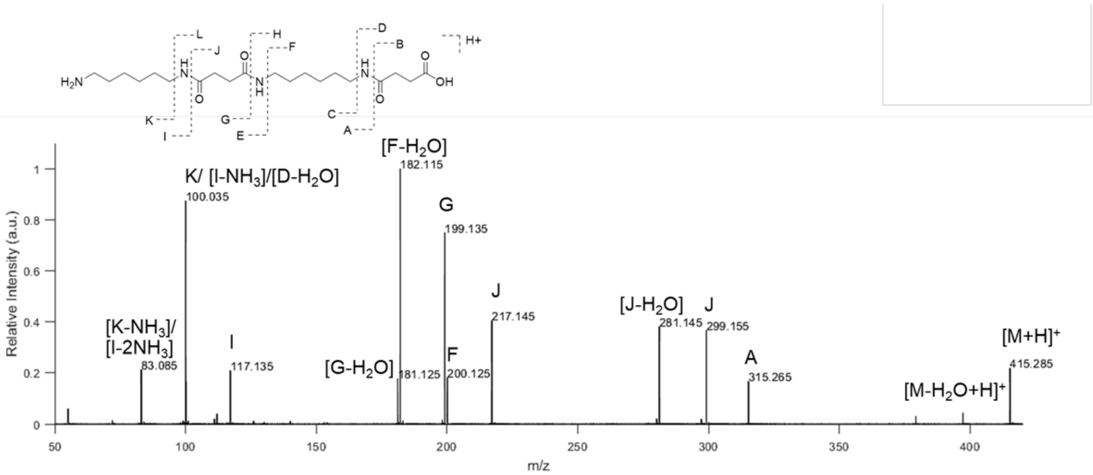


**Figure S56: I.DOT/OPSI-MS$^2$ of MSMS.** A collision energy of 40 eV was used.

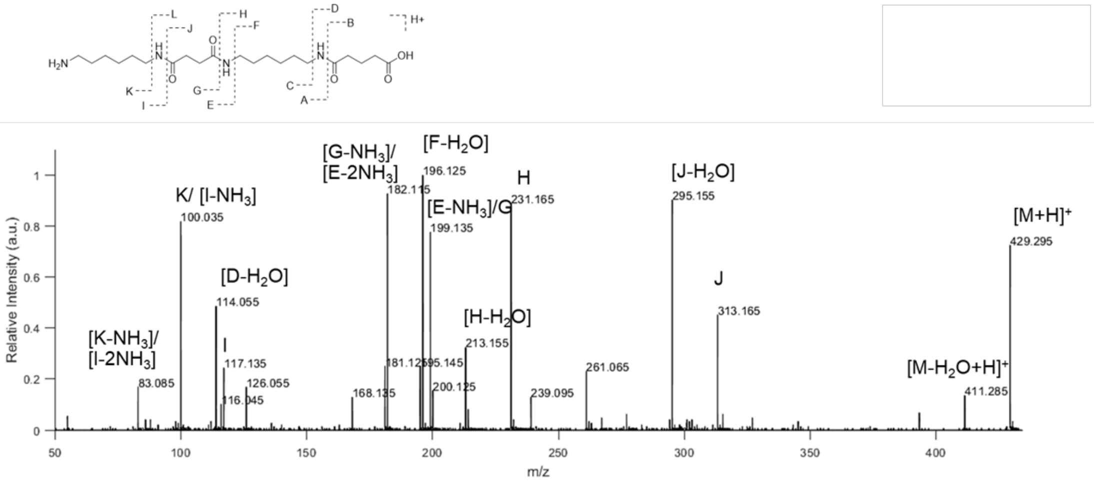


**Figure S57: I.DOT/OPSI-MS$^2$ of GMSM.** A collision energy of 40 eV was used.

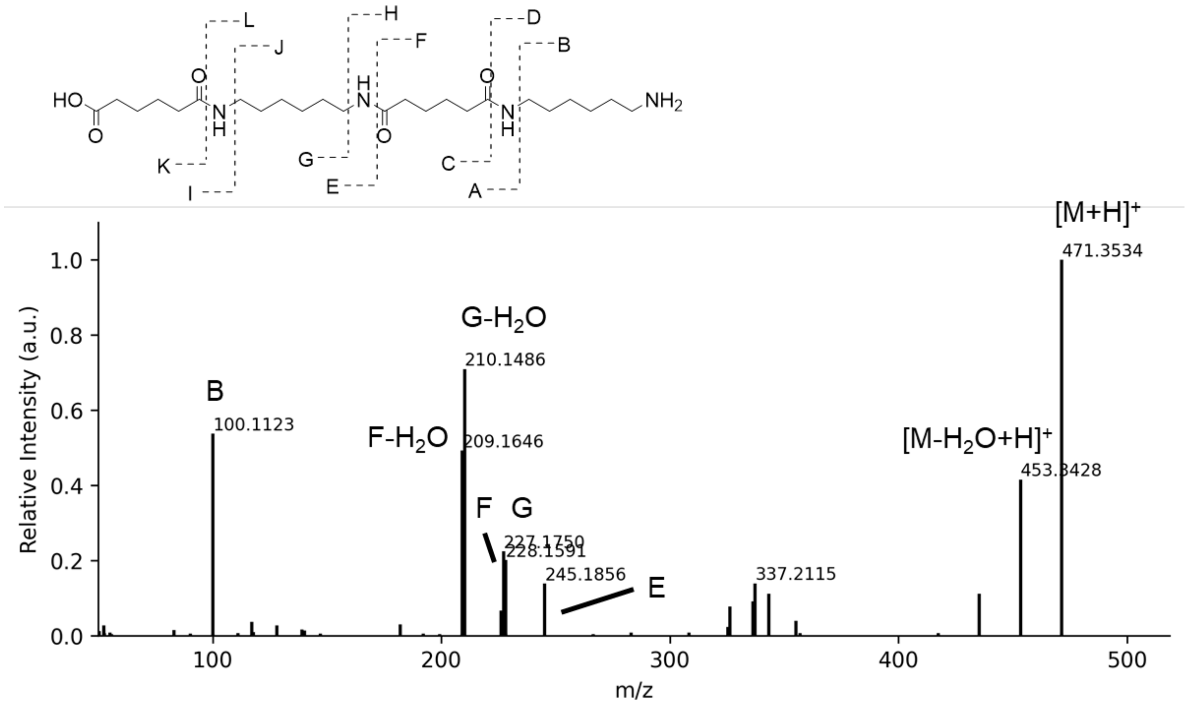


**Figure S58: I.DOT/OPSI-MS$^2$ of MAMA.** A collision energy of 35 eV was used.**MAM** and **A** were used as the substrates.

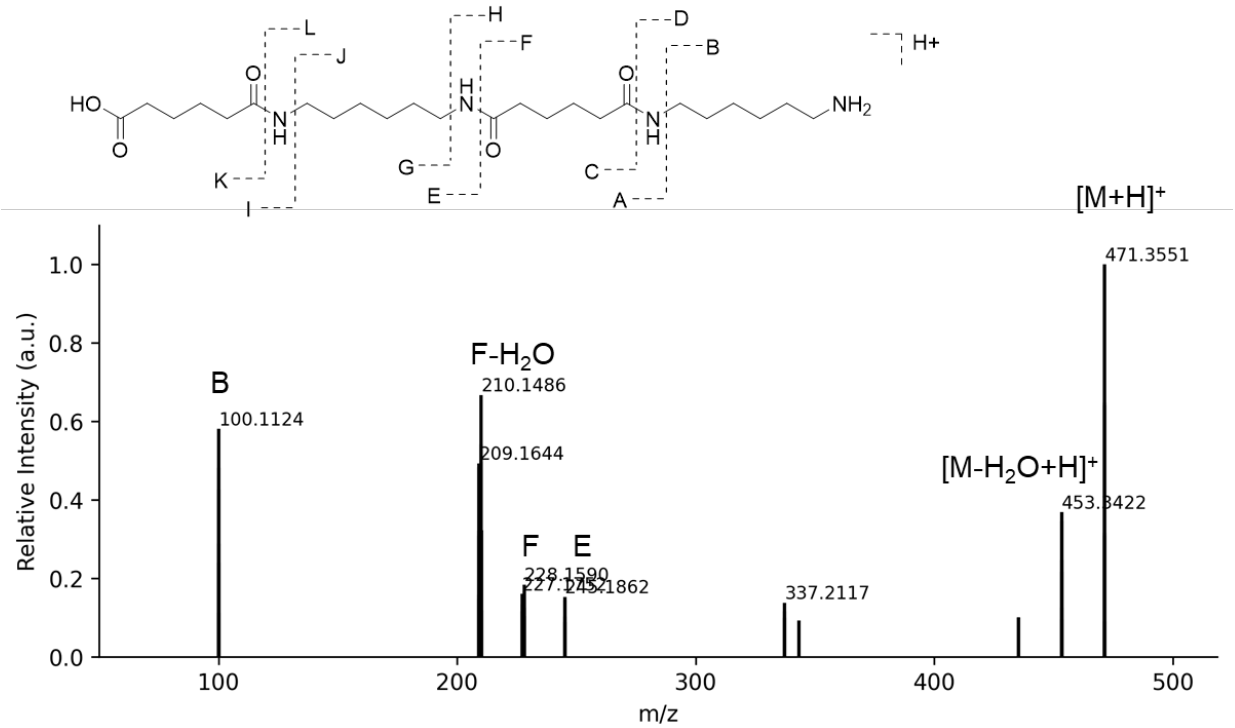


**Figure S59: I.DOT/OPSI-MS[2] of MAMA.** A collision energy of 35 eV was used. **AMA** and **M** were used as the substrates.

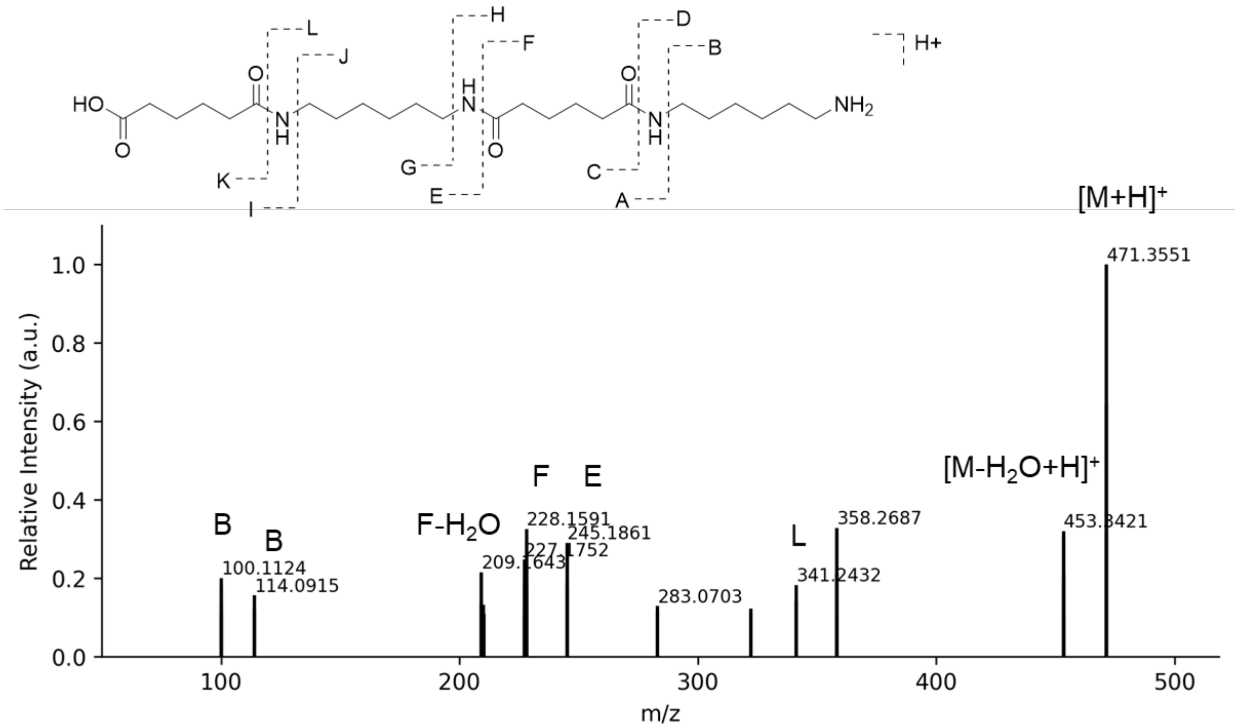


**Figure S60: I.DOT/OPSI-MS$^2$ of MAMA.** A collision energy of 35 eV was used. **MA** was used as the substrate.

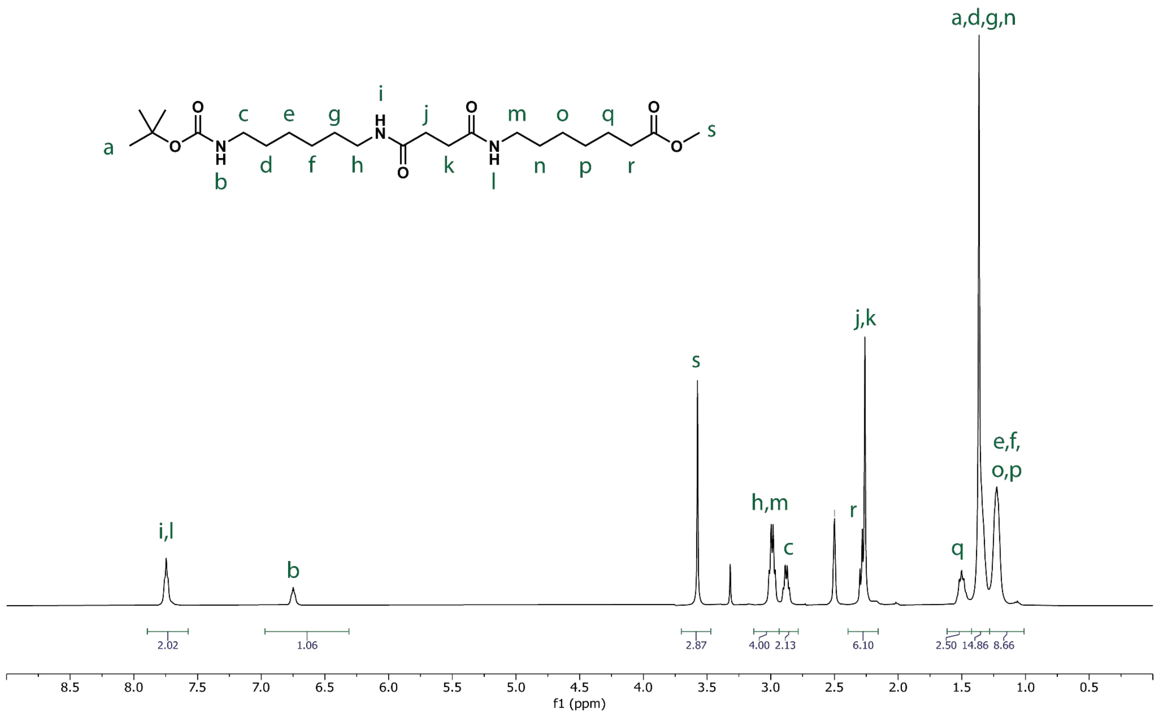


**Figure S61: $^{1}$H NMR spectrum of Boc-MSP-OMe in DMSO-d$_6$.**

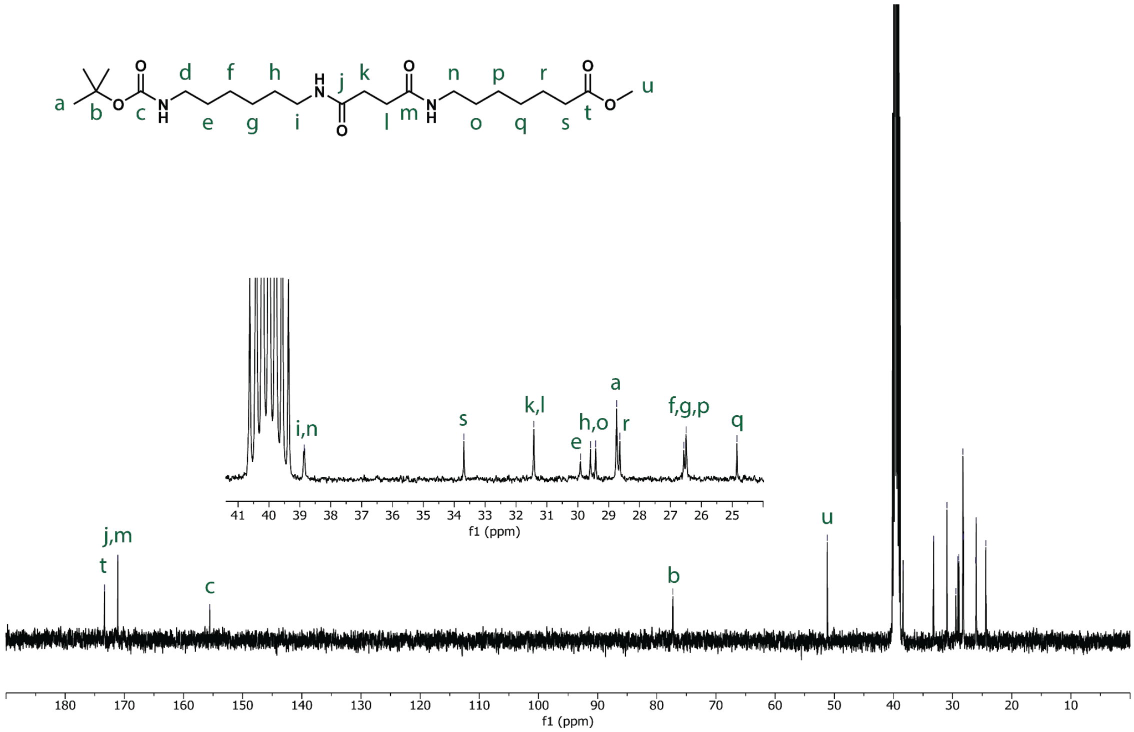


**Figure S62: $^{13}$C NMR spectrum of Boc-MSP-OMe in DMSO-d$_6$.**

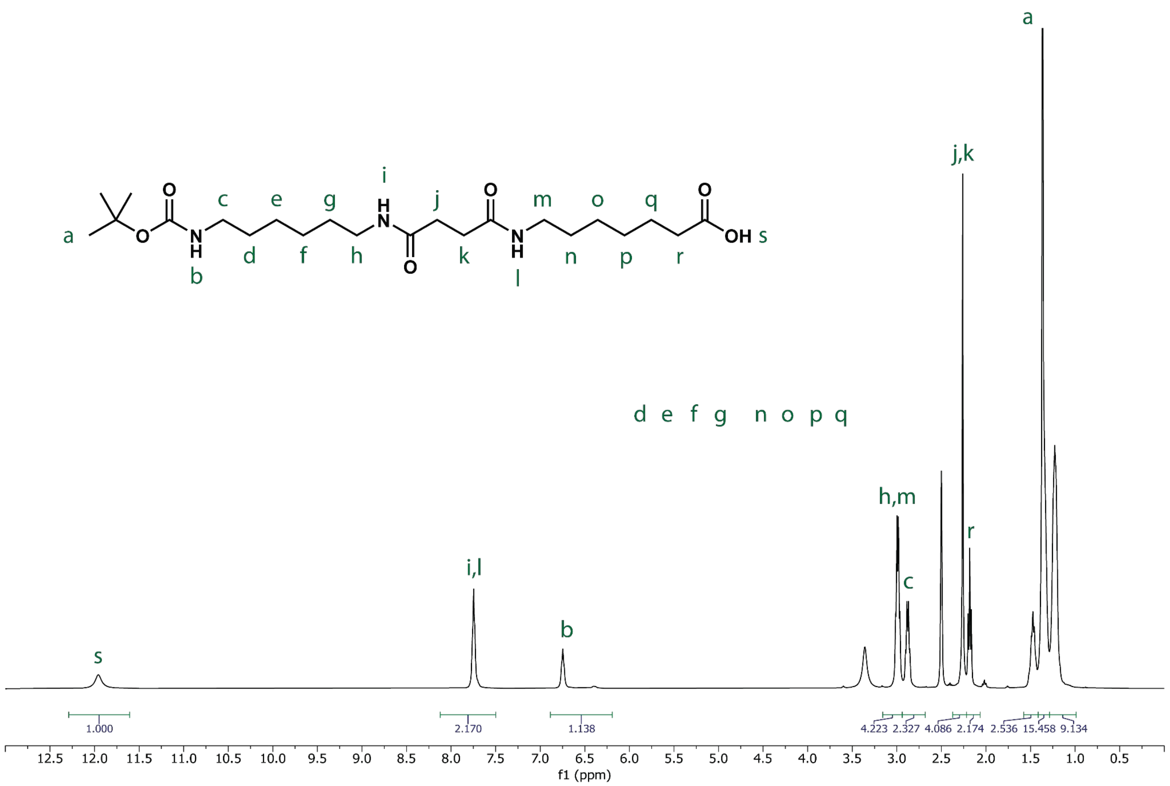


**Figure S63: $^{1}$H NMR spectrum of Boc-MSP in DMSO-d$_6$.**

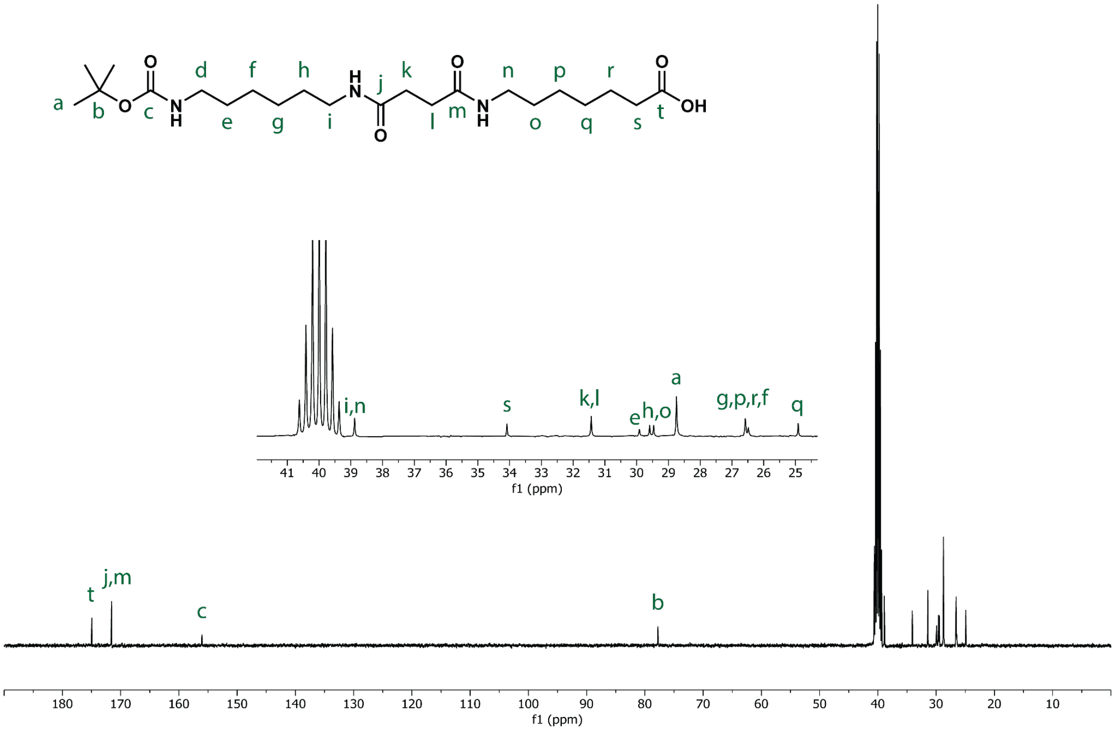


**Figure S64: $^{13}$C NMR spectrum of Boc-MSP in DMSO-d$_6$.**

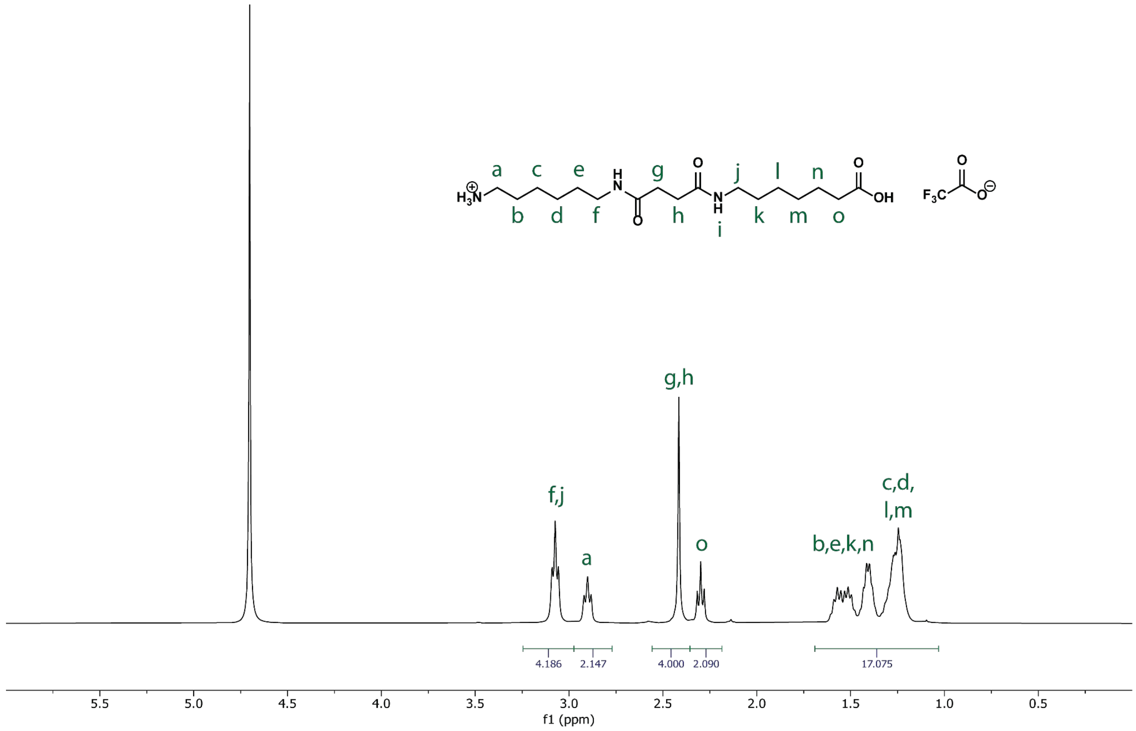

**Figure S65: $^{1}$H NMR spectrum of MSP trifluoroacetate in $D_2O$.**

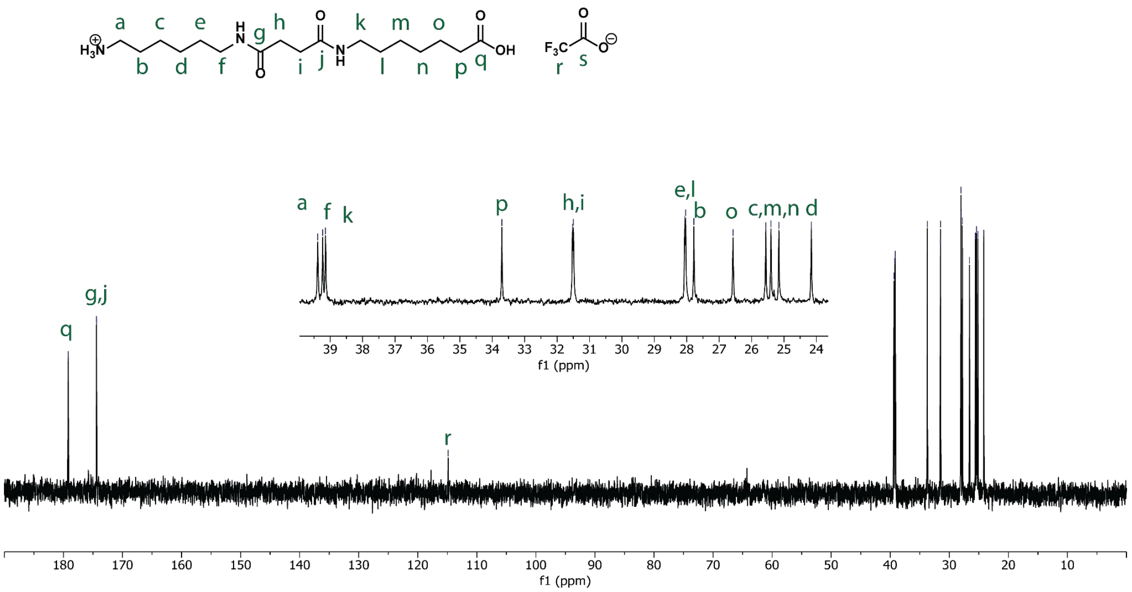

**Figure S66: $^{13}$C NMR spectrum of MSP trifluoroacetate in $D_2O$.**

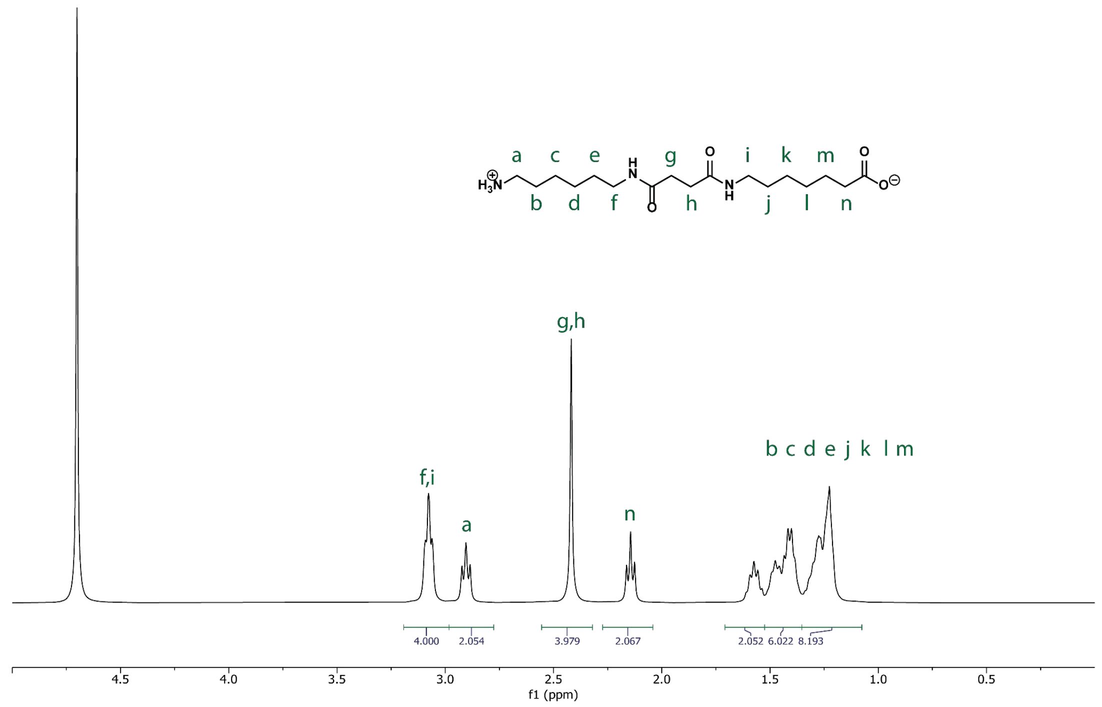


**Figure S67: $^{1}$H NMR spectrum of MSP in $D_2O$.**

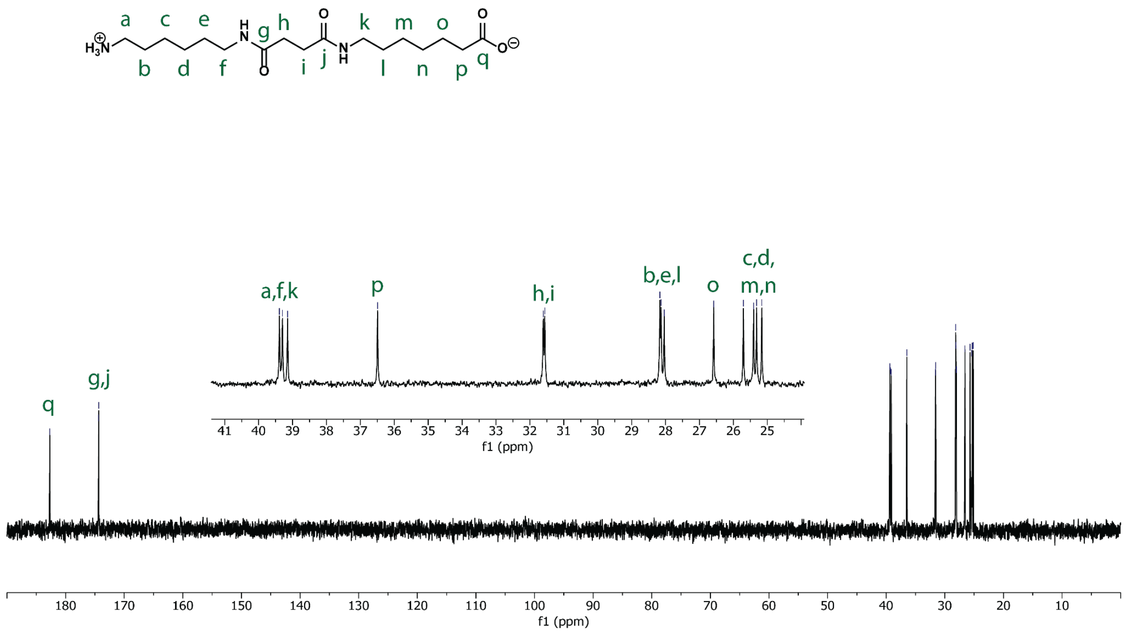


**Figure S68: $^{13}$C NMR spectrum of MSP in $D_2O$.**

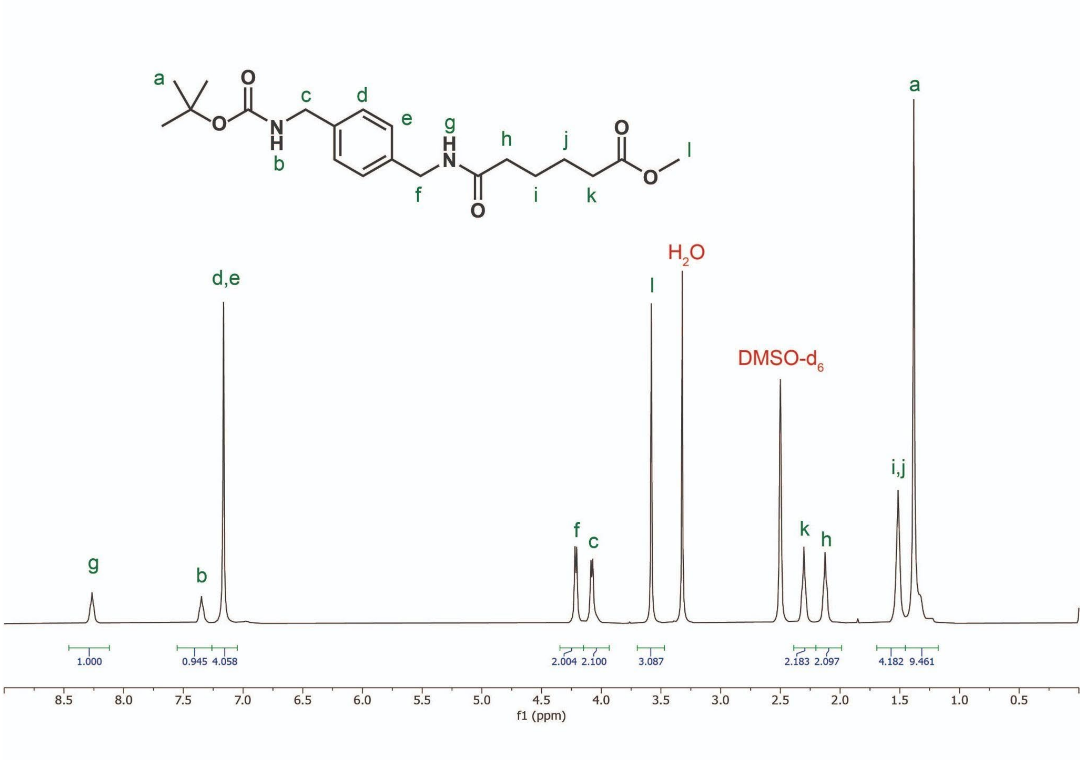


**Figure S69: $^1$H NMR spectrum of Boc-XA-OMe in DMSO-d$_6$.**

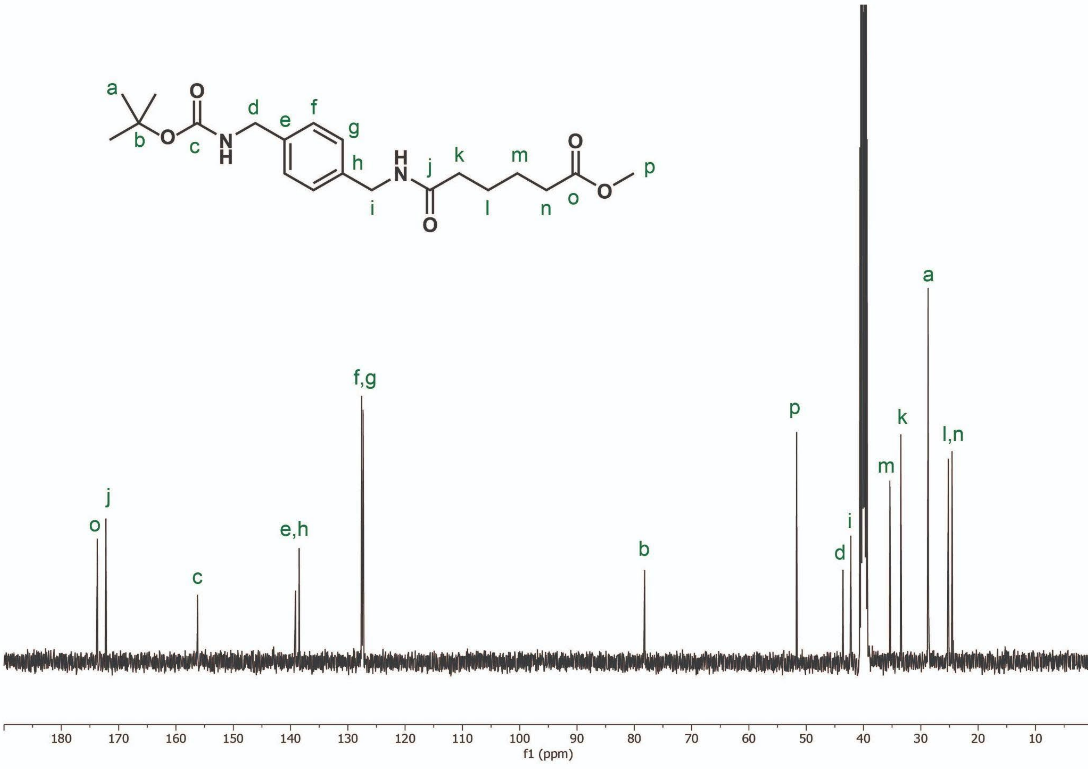


**Figure S70: $^{13}$C NMR spectrum of Boc-XA-OMe in DMSO-d$_6$.**

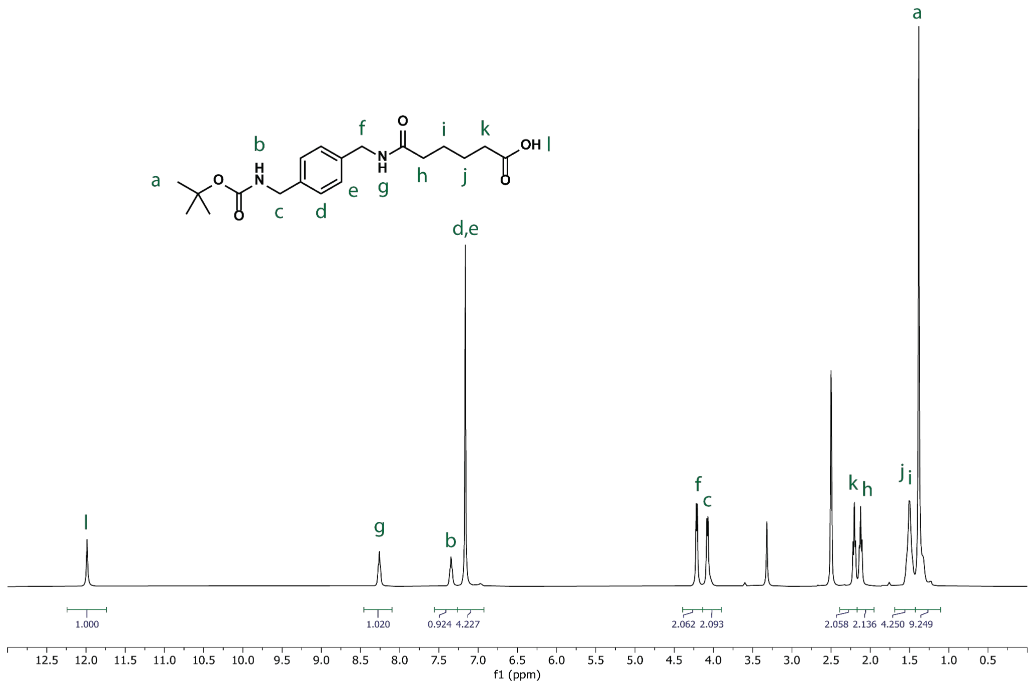


**Figure S71: $^{1}$H NMR spectrum of Boc-XA in DMSO-$d_6$.**

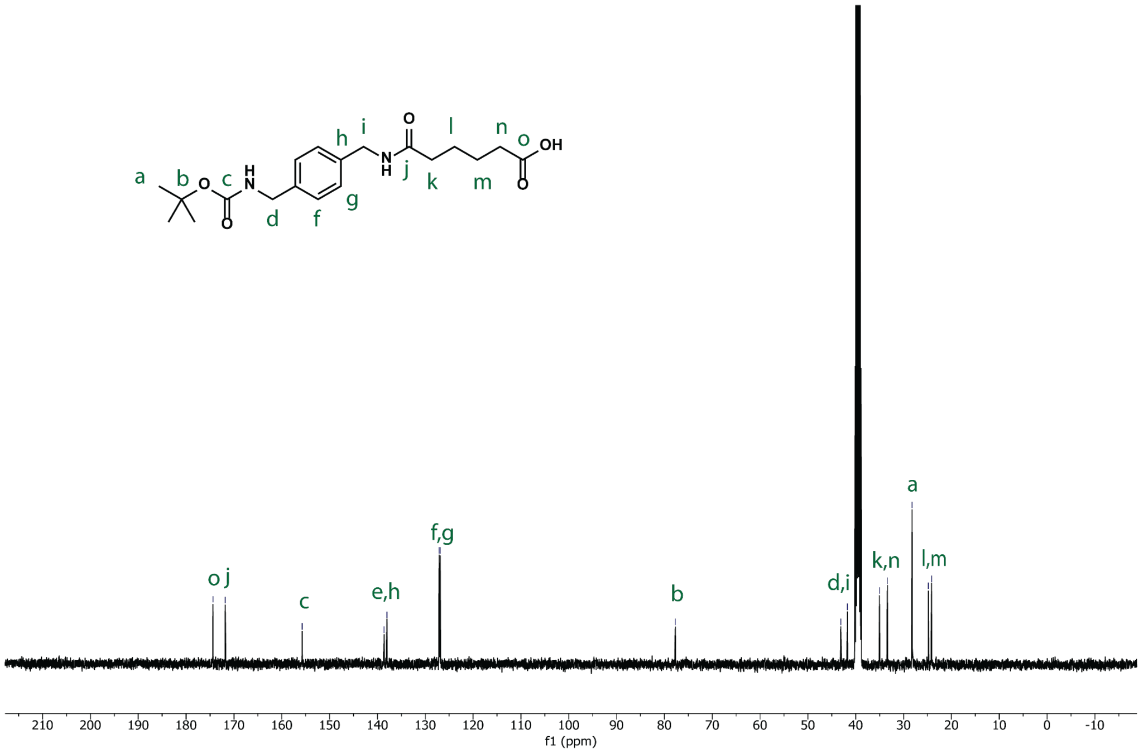


**Figure S72: $^{13}$C NMR spectrum of Boc-XA in DMSO-$d_6$.**

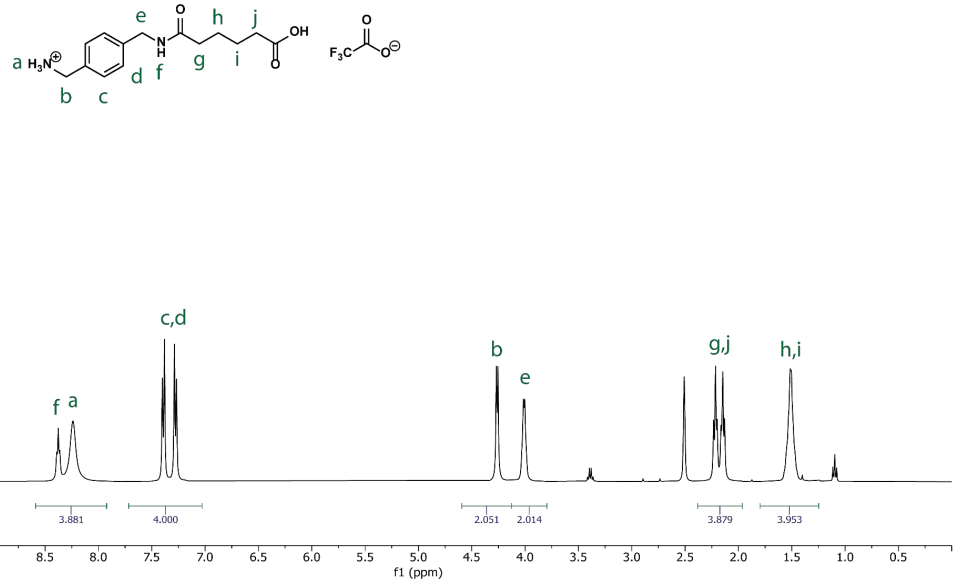


**Figure S73: $^{1}$H NMR spectrum of XA trifluoroacetate in DMSO-d$_6$.**

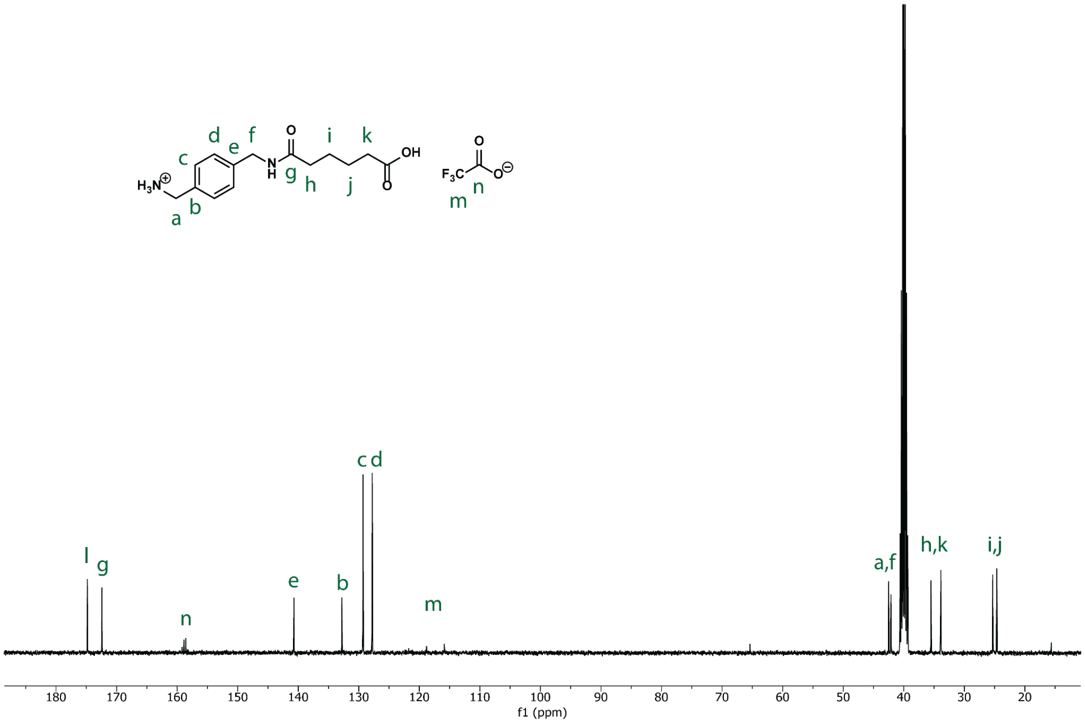


**Figure S74: $^{13}$C NMR spectrum of XA trifluoroacetate in DMSO-d$_6$.**

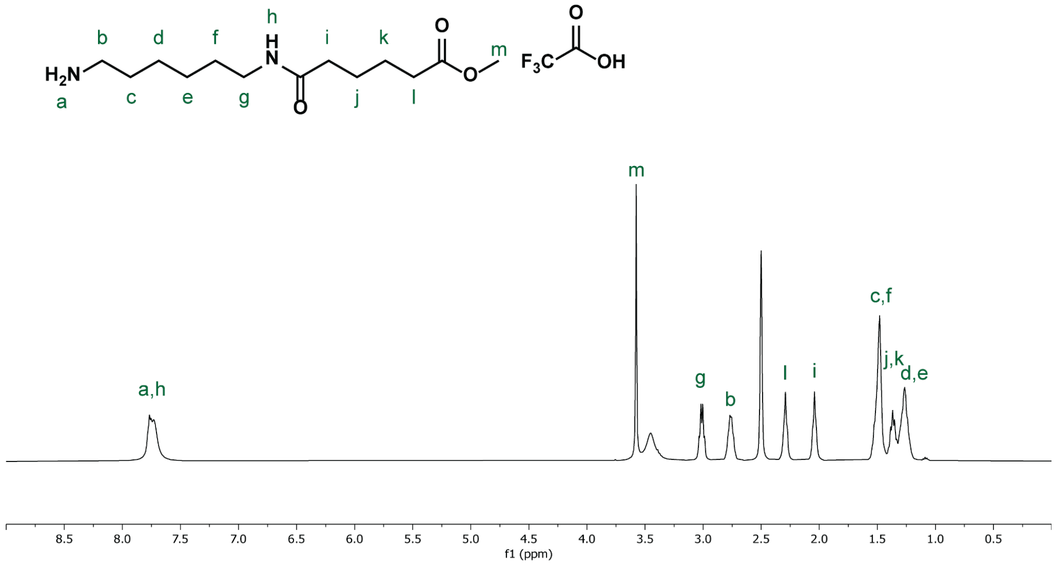


**Figure S75: $^{1}$H NMR spectrum of MA-OMe in DMSO-d$_6$.**

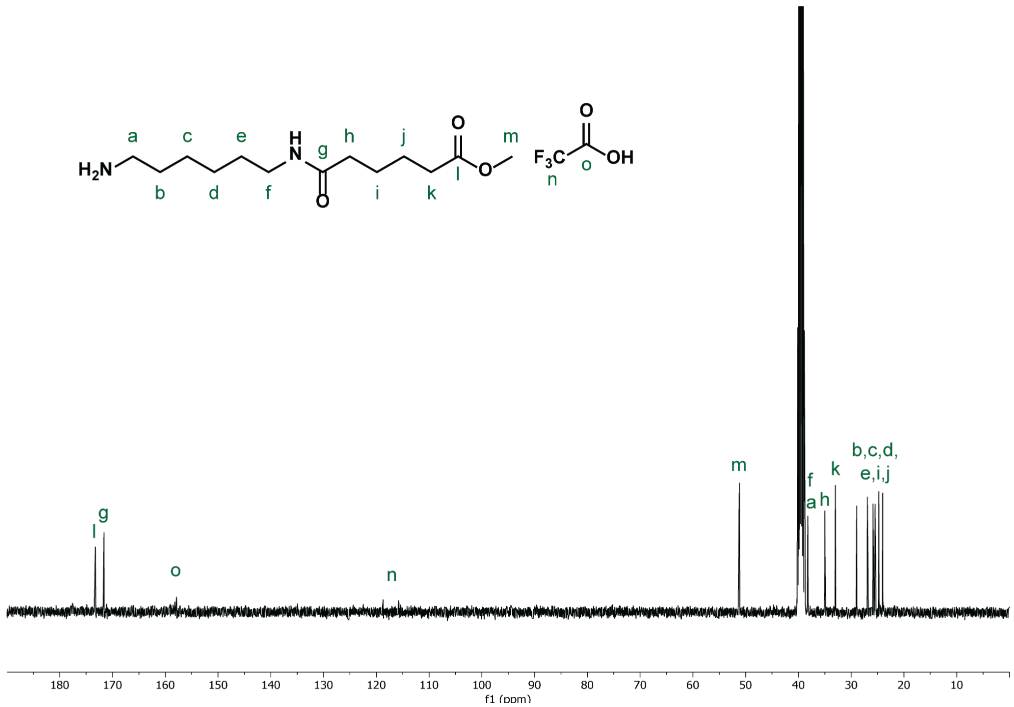


**Figure S76: $^{13}$C NMR spectrum of MA-OMe in DMSO-d$_6$.**

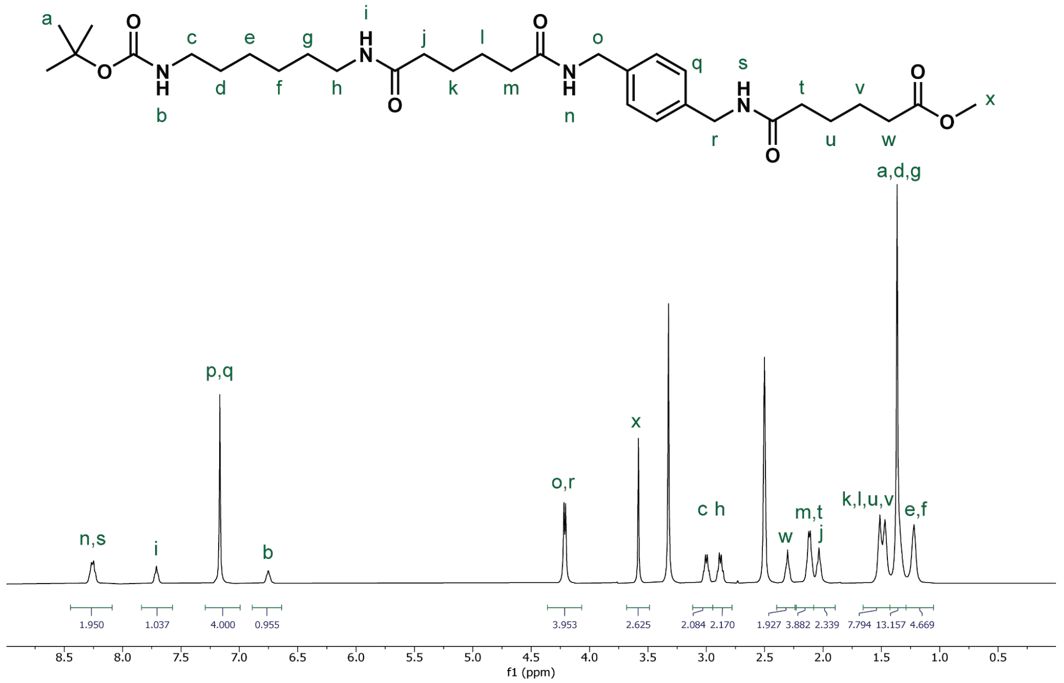


**Figure S77: $^{1}$H NMR spectrum of Boc-MAXA-OMe in DMSO-d$_6$.**

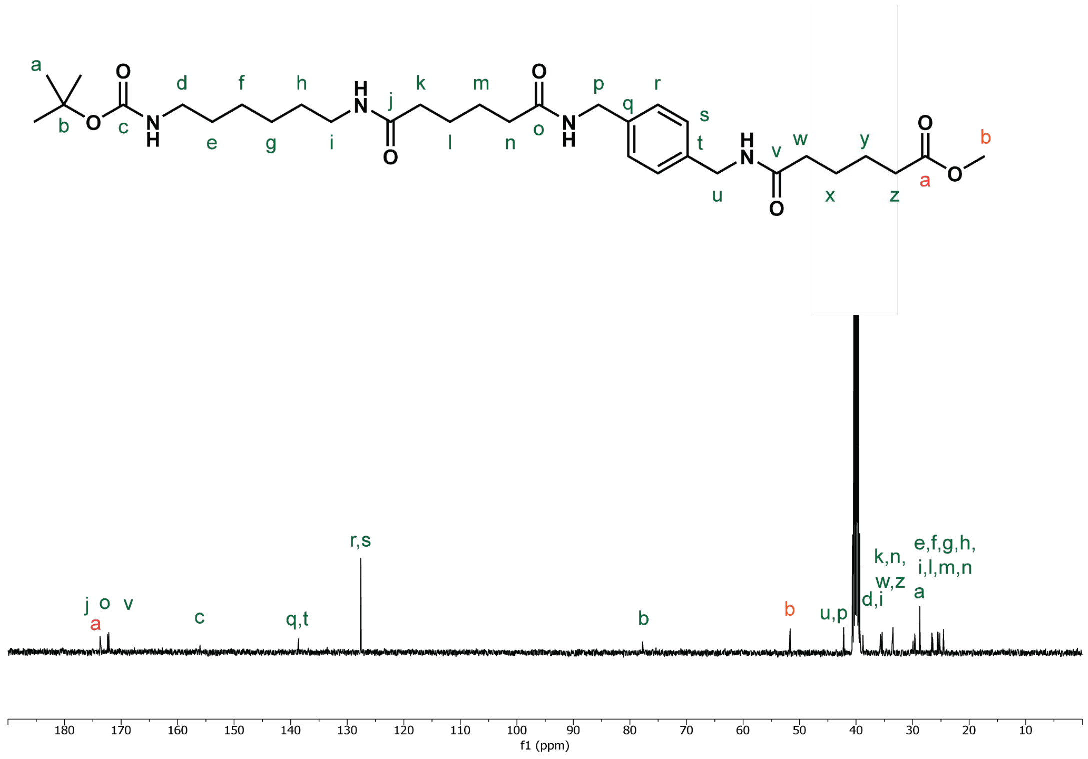


**Figure S78: $^{13}$C NMR spectrum of Boc-MAXA-OMe in DMSO-d$_6$.**

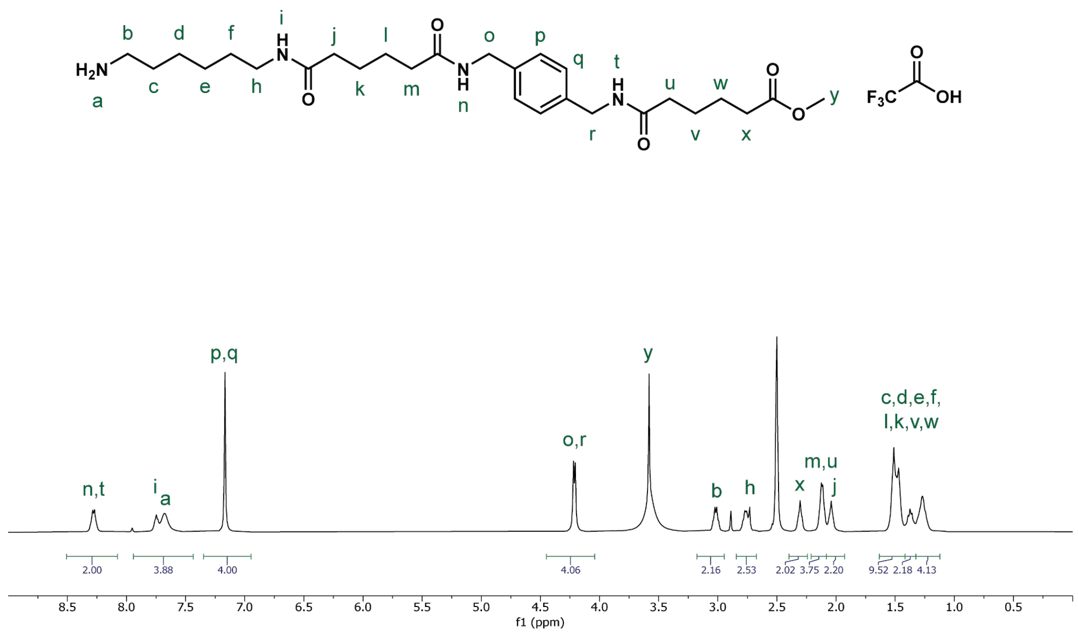

**Figure S79: $^{1}$H NMR spectrum of MAXA-OMe in DMSO-d$_6$.**

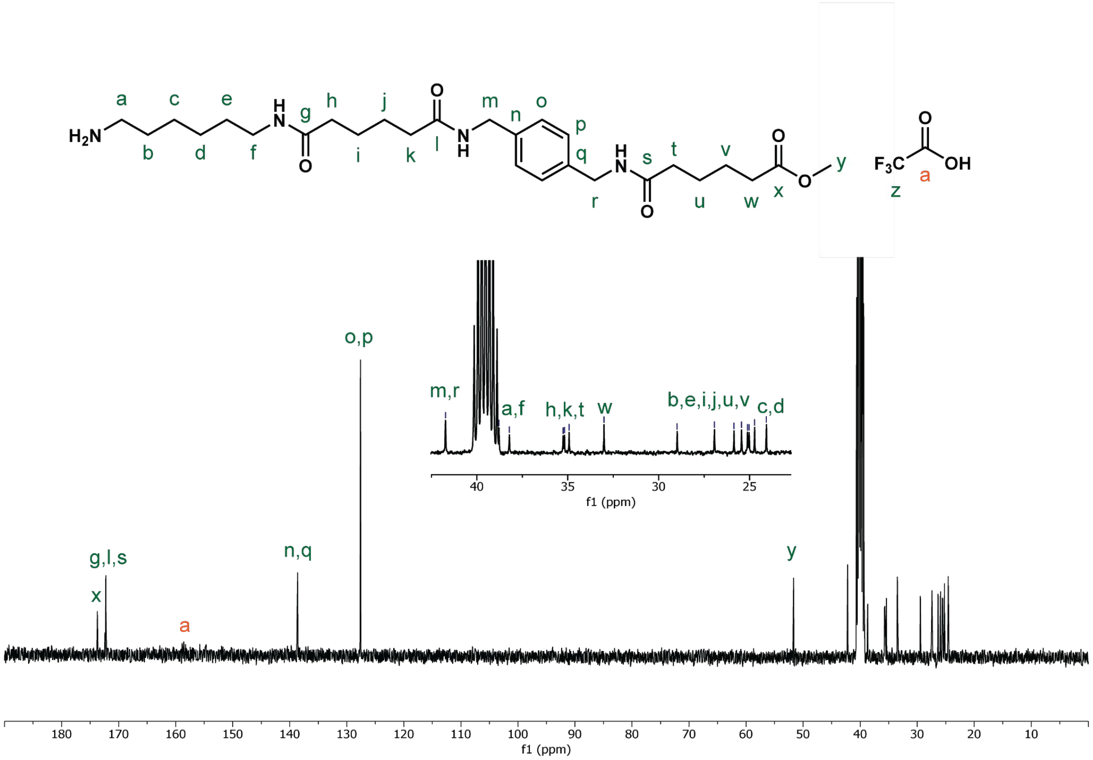

**Figure S80: $^{13}$C NMR spectrum of MAXA-OMe in DMSO-d$_6$.**

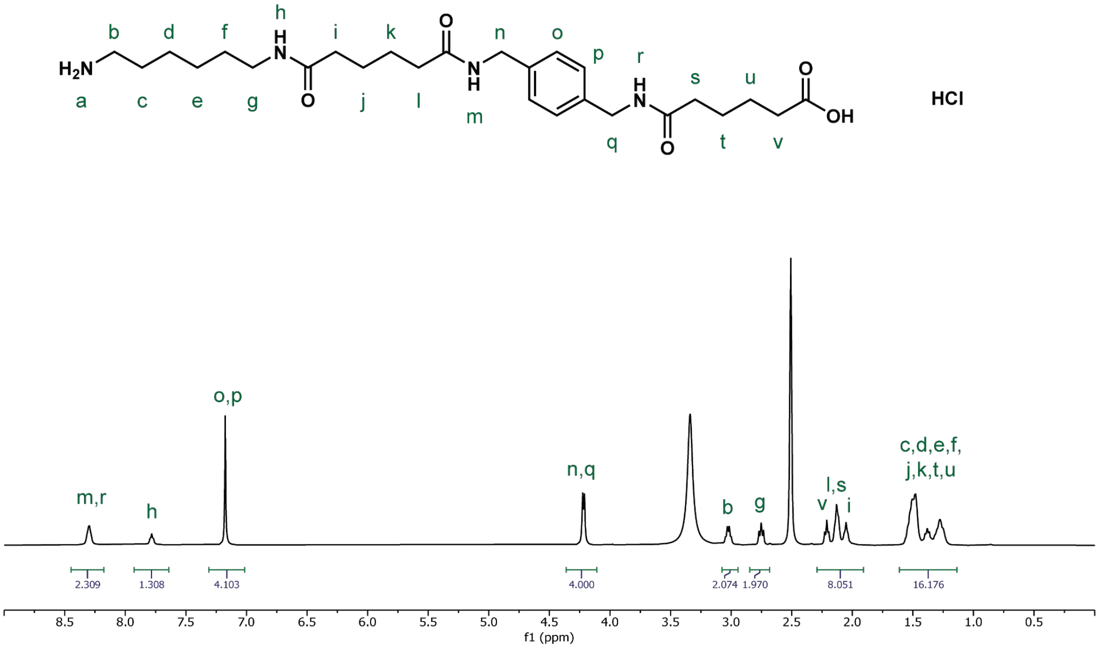

Figure S81: [1]H NMR spectrum of MAXA HCl in DMSO-$d_6$.

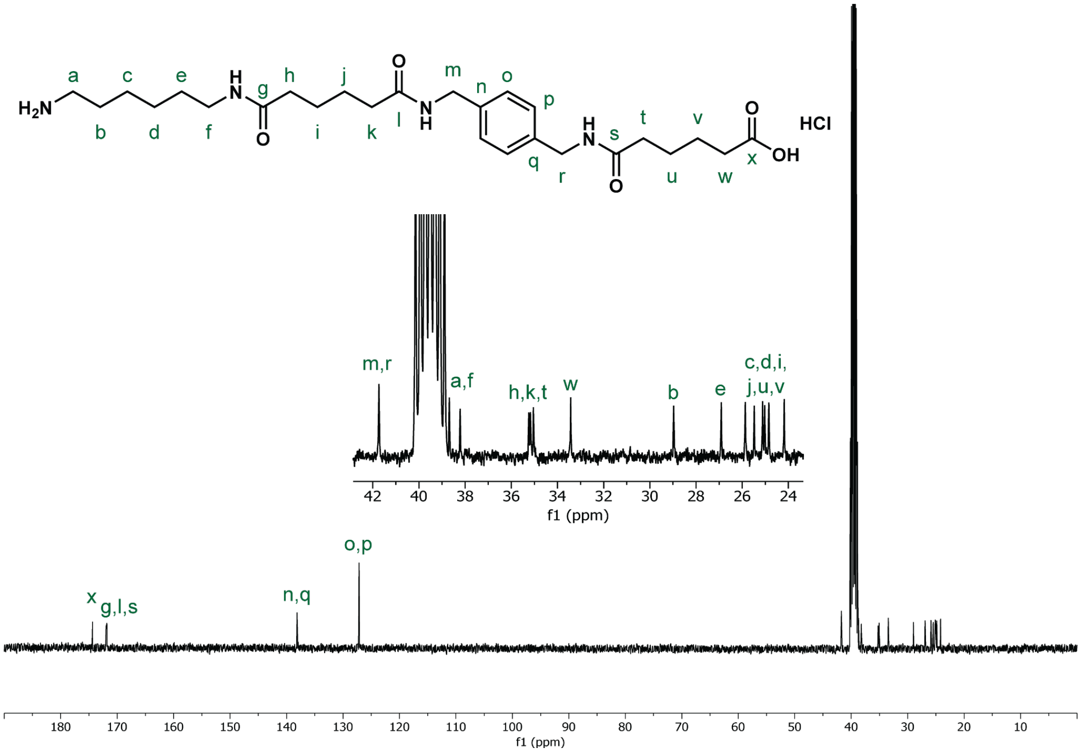

Figure S82: [13]C NMR spectrum of MAXA HCl in DMSO-$d_6$.

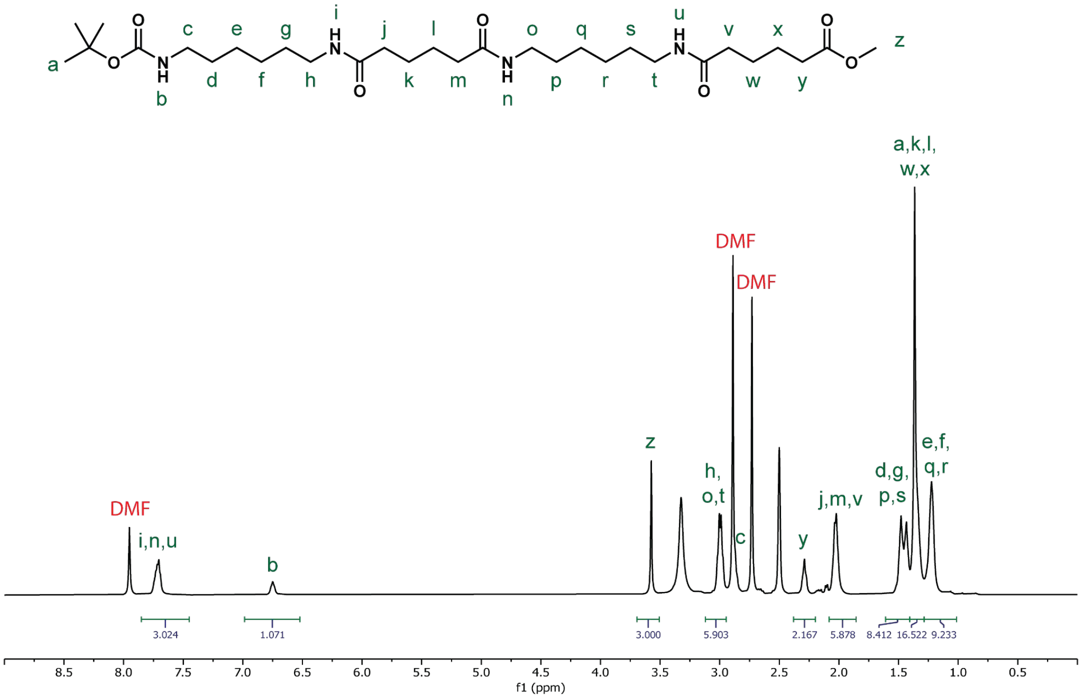


**Figure S83: $^{1}$H NMR spectrum of Boc-MAMA-OMe in DMSO-$d_6$.**

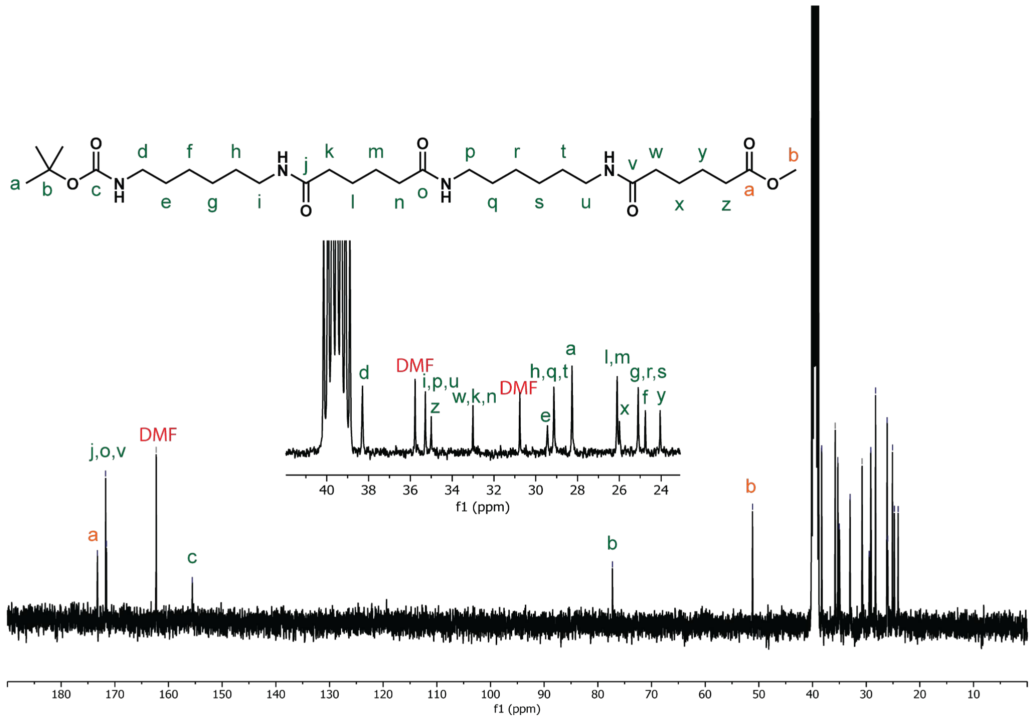


**Figure S84: $^{13}$C NMR spectrum of Boc-MAMA-OMe in DMSO-$d_6$.**

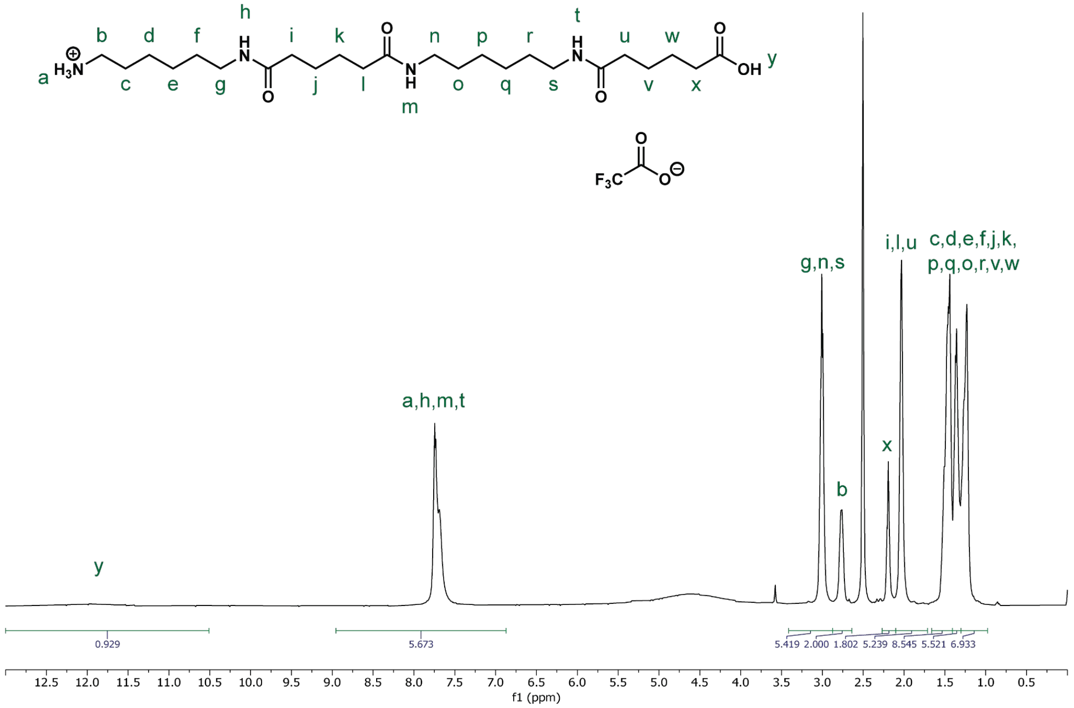


**Figure S85: $^{1}$H NMR spectrum of MAMA trifluoroacetate in DMSO-d$_6$.**

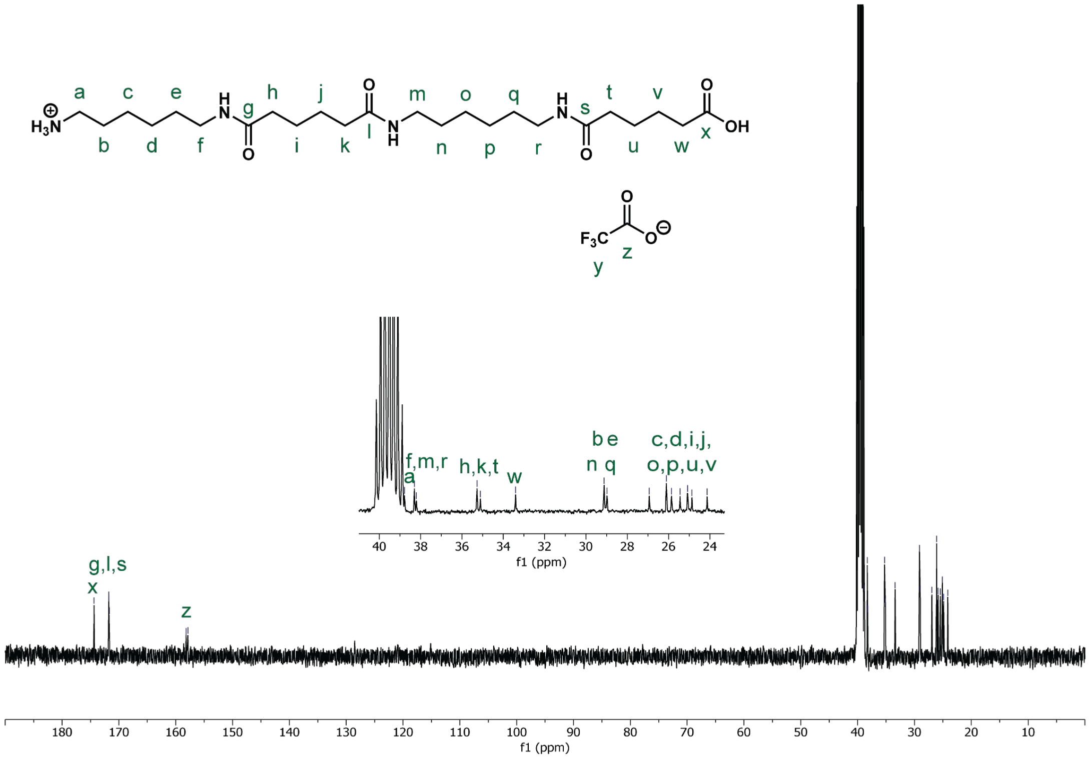


**Figure S86: $^{13}$C NMR spectrum of MAMA trifluoroacetate in DMSO-d$_6$.**

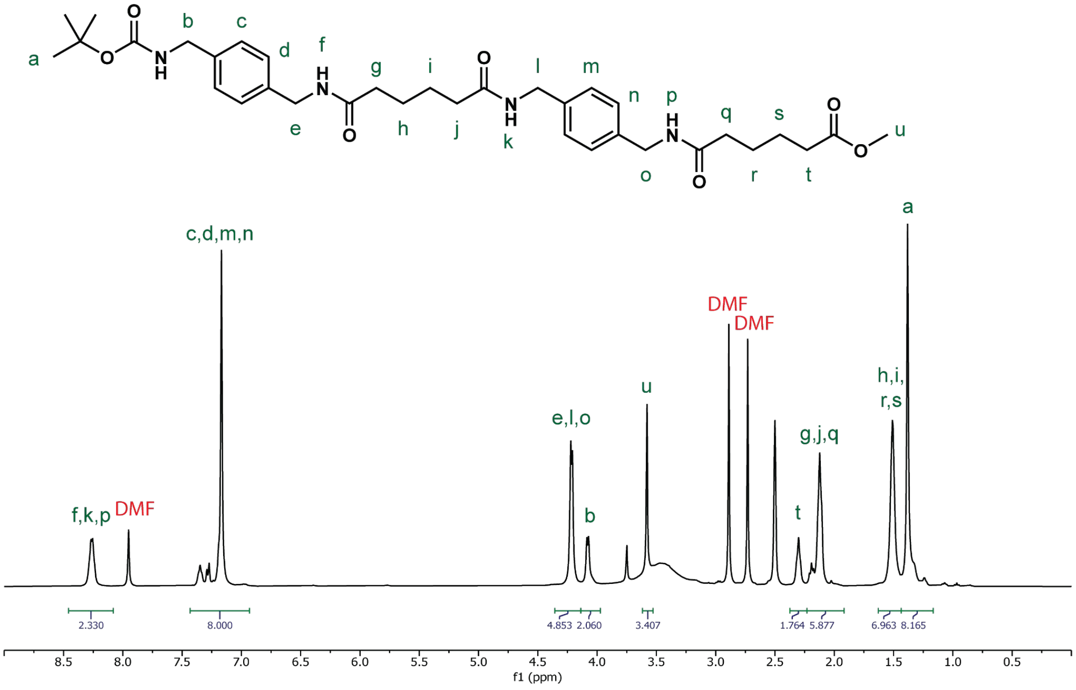


**Figure S87: $^{1}$H NMR spectrum of Boc-XAXA-OMe in DMSO-d$_6$.**

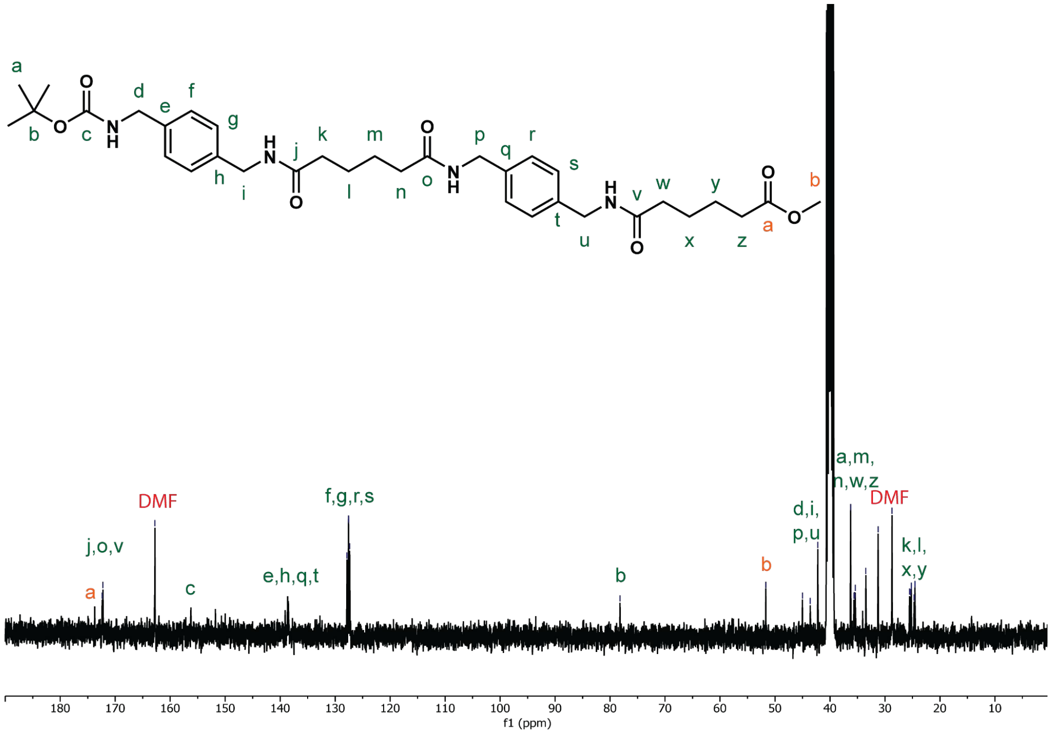


**Figure S88: $^{13}$C NMR spectrum of Boc-XAXA-OMe in DMSO-d$_6$.**

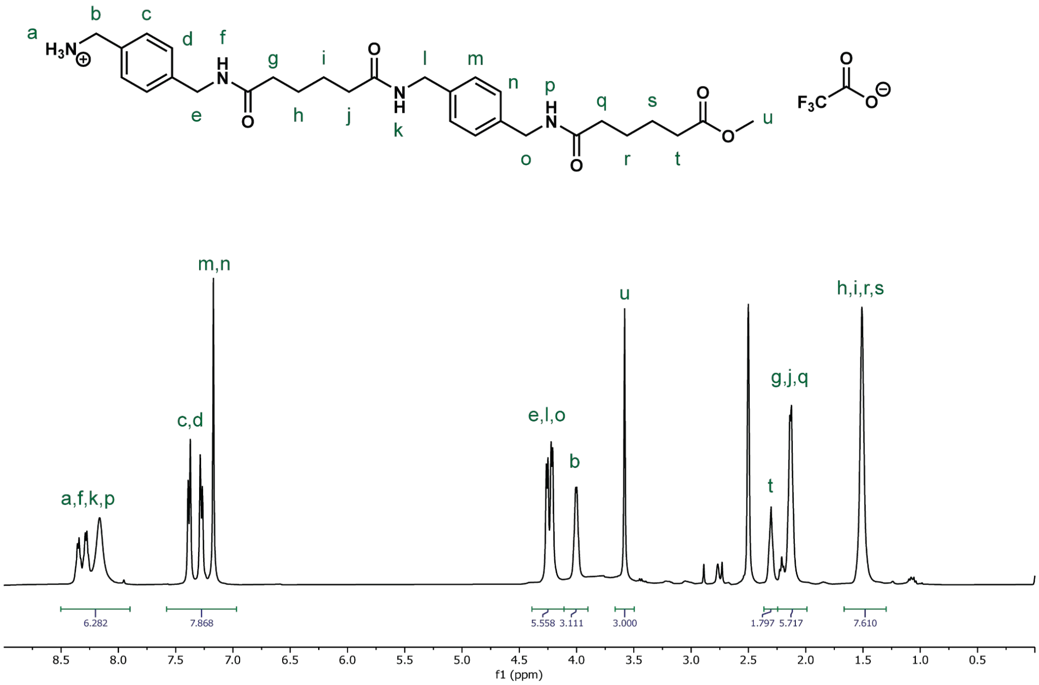


**Figure S89: $^{1}$H NMR spectrum of XAXA-OMe in DMSO-d$_6$.**

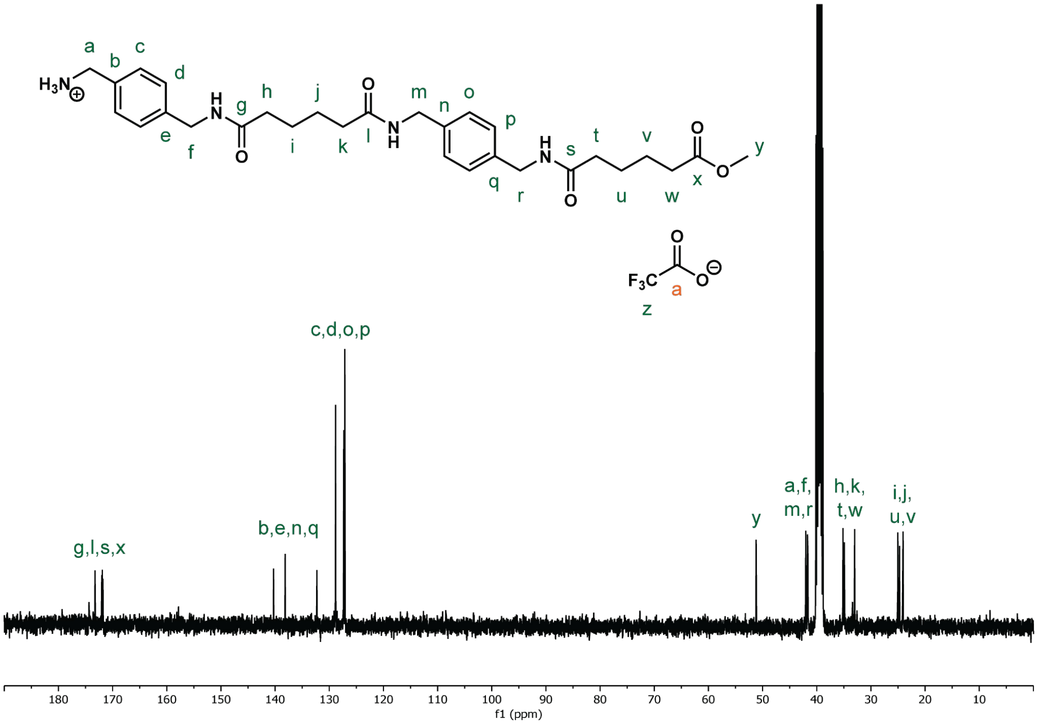


**Figure S90: $^{13}$C NMR spectrum of XAXA-OMe in DMSO-d$_6$.**
